# A Game of Thrones at Human Centromeres II. A new molecular/evolutionary model

**DOI:** 10.1101/731471

**Authors:** William R. Rice

## Abstract

Human centromeres are remarkable in four ways: they are i) defined epigenetically by an elevated concentration of the histone H3 variant CENP-A, ii) inherited epigenetically by trans-generational cary-over of nucleosomes containing CENP-A, iii) formed over unusually long and complex tandem repeats (Higher Order Repeats, HORs) that extend over exceptionally long arrays of DNA (up to 8 Mb), and iv) evolve in such a rapid and punctuated manner that most HORs on orthologous chimp and human chromosomes are in different clades. What molecular and evolutionary processes generated these distinctive characteristics? Here I motivate and construct a new model for the formation, expansion/contraction, homogenization and rapid evolution of human centromeric repeat arrays that is based on fork-collapse during DNA replication (in response to proteins bound to DNA and/or collisions between DNA and RNA polymerases) followed by out-of-register re-initiation of replication via Break-Induced Repair (BIR). The model represents a new form of molecular drive. It predicts rapid and sometimes punctuated evolution of centromeric HORs due to a new form of intragenomic competition that is based on two features: i) the rate of tandem copy number expansion, and ii) resistance to invasion by pericentric heterochromatin within a centromere’s HOR array. These features determine which variant array elements will eventually occupy a pivotal region within a centromeric repeat array (switch-point) that gradually expands to populate the entire array. In humans, continuous HOR turnover is predicted due to intra-array competition between three repeat types with an intransitive hierarchy: A < B < C < A, where A = short, single-dimer HORs containing one monomer that binds centromere protein-B (CENP-B) and another that does not, B = moderately longer HORs composed of ≥ 2 dimers, and C = substantially longer HORs that lose their dimeric modular structure. Continuous turnover of proteins that bind centromeric DNA (but these proteins are not constituents of the kinetochore) and polygenic variation influencing position-effect variegation are predicted to cause rapid turnover of centromeric repeats in species lacking HORs and/or CENP-B binding at centromeres. Evolution at centromeres is a molecular ‘*Game-of-Thrones*’ because centromeric sequences ‘reign’ due to an epigenetic ‘crown’ of CENP-A that is perpetually ‘usurped’ by new sequences that more rapidly assemble large ‘armies’ of tandem repeats and/or resist ‘invasion’ from a surrounding ‘frontier’ of percentric heterochromatin. These ‘regal transitions’ occur in a backdrop of slashing and decapitation (fork-collapse generating truncated sister chromatids) in the context of promiscuous sex that is frequently incestuous (out-of-register BIR between sibling chromatids).

## Introduction

The generally accepted model for the evolution of human centromeric DNA (composed of Higher Order Repeats [HORs] = repeats of repeats; Figure 1) was developed by Smith (1976). He used computer simulation to show that out-of-register recombination between sister chromatids can generate repeated sequences that are qualitatively similar to the long arrays of homogeneous repeats seen at satellite DNAs. The Smith model is a neutral model of evolution that predicts that the DNA sequence of the repeats seen in satellite DNAs will be effectively random –except for the avoidance of intrinsically harmful sequences, such those that form secondary structure that interferes with DNA replication. In the companion paper (Rice 2019), I showed that the human HORs found at active centromeric repeat arrays across all 24 chromosomes are highly structured at multiple levels, and that this structure could feasibly come from some sort of functional constraint that is not predicted by the Smith (1976) model. I also showed that the exceptionally large sizes of centromeric HOR arrays (far larger than required for cellular functioning), and their observed patterns of length variation on the sex chromosomes, indicate that some process other than unequal crossing over (between sister chromatids) is responsible for generating most length variation at centromeric arrays and driving the rapid evolution of centromeric HORs.

**Figure 1.**
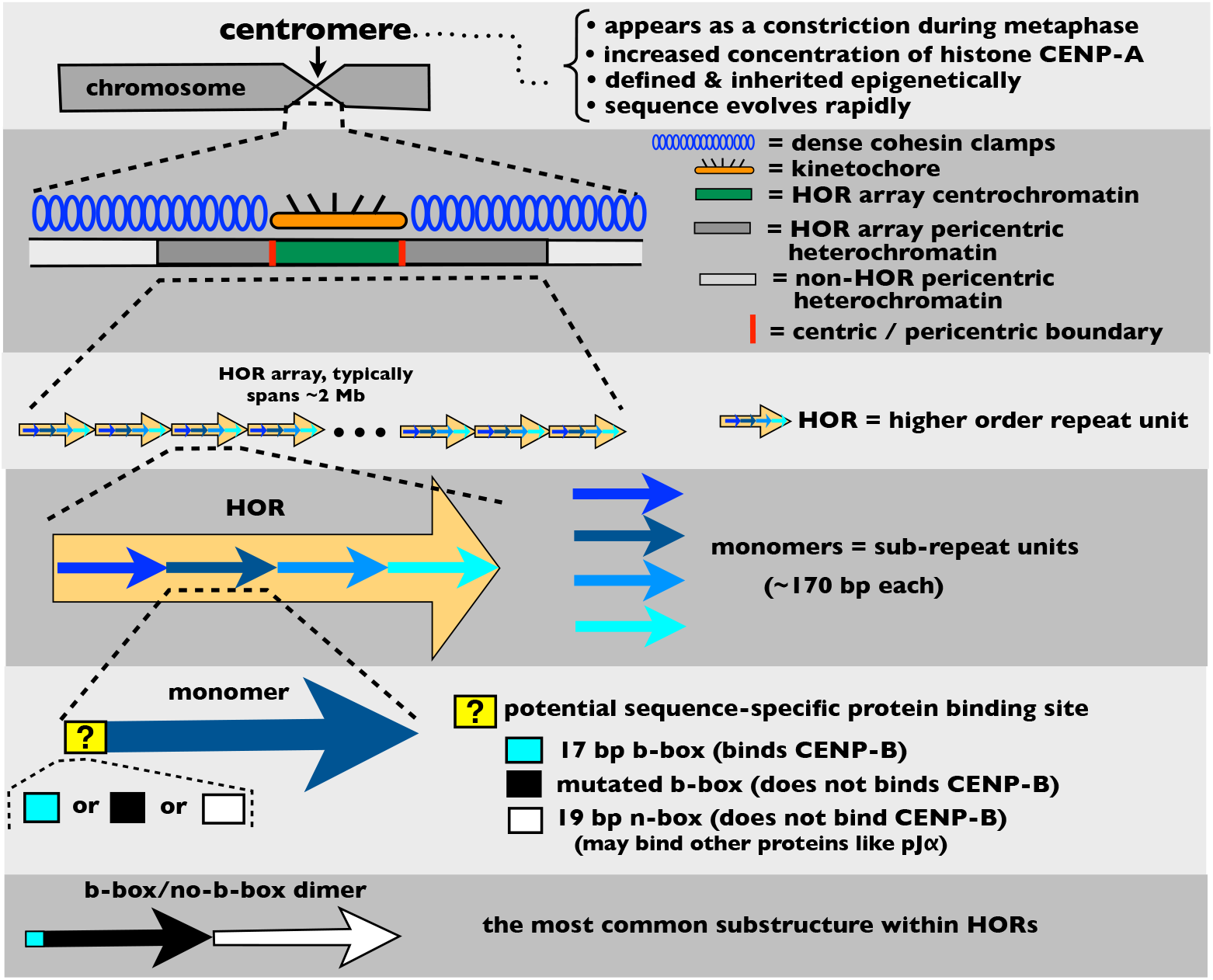
Summary of the structure of human centromeres and Higher Order Repeat HOR) arrays.

Here I develop an alternative to the Smith model that is based on out-of-register re-initiation of DNA replication after fork-collapse. This process is assumed to generate most of the extensive length variation seen at human centromeric HOR arrays, and may apply to other species with regional centromeres. The model focuses on sequence variation within a centromeric HOR array that influences: i) the rate at which repeat units laterally expand (tandem increase in copy number) within the array, and ii) the rate of the invasion of pericentric heterochromatin into the centrochromatin that nucleates the kinetochore (Figure 1). These two rates determine the ability of alternative HOR sequences –that are present within the same centromeric repeat array– to compete for proportional representation, and hence which sequence will eventually predominate within the array. At human centromeres, both of these rates are predicted to be strongly influenced by the binding of CENtromere Protein B (CENP-B) and how this binding is distributed among neighboring monomers within an array. The model predicts the rapid –and sometimes punctuated– evolution seen at human centromeric HORs, as well as many aspects of their complex structure. Although the model was developed for human centromeric HORs, I show that its foundational features should apply more broadly to other species that have epigenetically defined, regional centromeres.

The molecular biology of some of the steps in the model is in a state of discovery, development and uncertainty. As a consequence, I will combine information from many model systems to construct feasible molecular underpinnings of the model. My objective is not to provide a model for centromere evolution that is correct in all details – I expect some details of the model to evolve as new information accrues over time. Instead, I will motivate a feasible foundation (starting point) for a new model of centromere evolution that is based on fork-stalling, fork-collapse and Break-Induced Repair (BIR) during DNA replication (rather than the unequal crossing over of the Smith model) and that is consistent with the many levels of structure that I identified in the companion paper (Rice 2019).

## Hypothesis for an evolutionary cycle of HOR expansion and replacement at human centromeres

The multifarious structure seen at the centromeric HORs of human chromosomes was used to generate a hypothesis for the evolution of centromeric HORs (Figure 2; see Figure 10 in Rice 2019 for a fuller description). This hypothesis describes a cycle that has has five steps:

i. HORs begin as simple b/n-box dimers (a pair of monomers, one containing a b-box sequence (that binds CENP-B) in the linker separating nucleosomes, and the other containing a n-box sequence (that does not bind CENP-B) at this position,
ii. HORs then grow (add monomers), but only by adding additional b/n-box dimer units,
iii. once sufficiently large, HORs continue to grow, but start losing modular b/n-box dimer structure by adding lone monomers and mutating b-boxes so that they no longer bind CENP-B,
iv. after losing substantial dimeric structure, HORs are replaced (by a new b/n-box dimer HOR) because they recruit substantially less CENP-C, and
v. replaced, inactive HOR arrays ultimately go extinct due to recurrent deletion pressure

**Figure 2.**
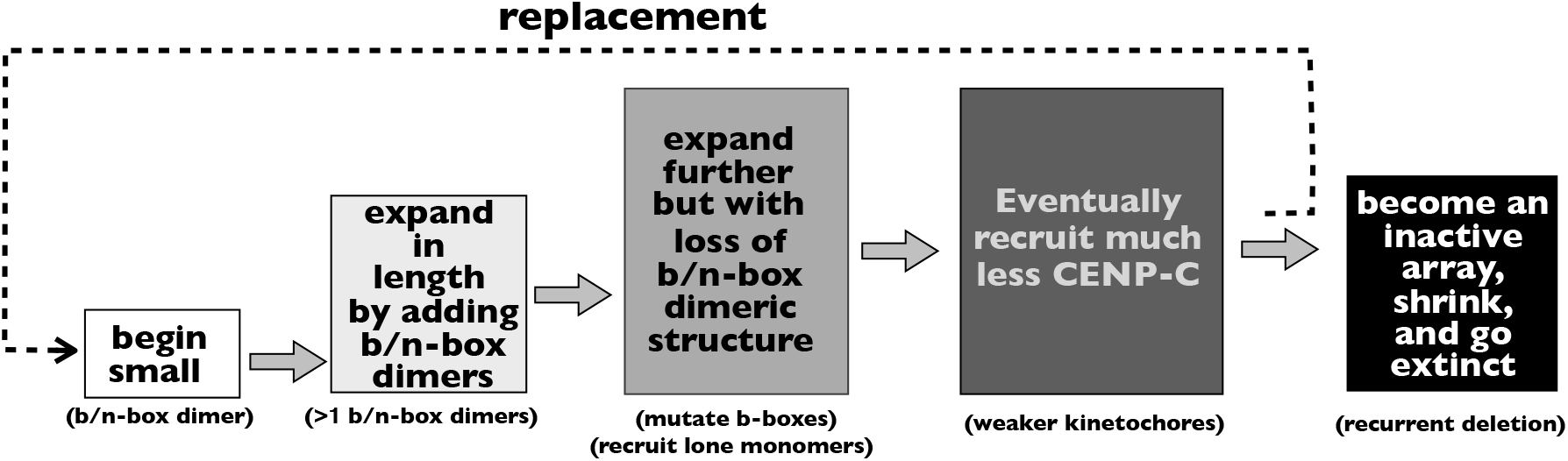
Hypothesis: the inferred pattern of change at human HORs across time. See Rice 2019 for detailed explanation of the trajectory.

While the main goal of this paper is to integrate information from many sources to deduce a new model for centromere evolution that is consistent will the multifarious structure reported (Rice 2019), a second goal is to evaluate this hypothesis.

## A new foundation for the rapid evolution of human centromeric HORs: replication fork-collapse followed by out-of-register BIR

The Smith model of repeat evolution (via unequal crossover between sister chromatids) does not predict the apparent cycle of HOR structure described in the above section, nor many of the structural characteristics found at human centromeric repeats (see Box 1 in the companion paper [Rice 2019]). An example of one of the inconsistencies between the Smith model and empirical data is the observed large size of HOR arrays (usually 2-3 Mb [Willard 1991], but sometimes exceeding 8 Mb [Miga et al. 2014]), compared to the minimum size required for cellular functioning (50-100 Kb; Lo et al. 1999; Yang et al. 2000; Okamoto et al. 2007). The unequal crossing over process lengthens the HOR array on one sister chromatid by the same amount that it shortens the other: so the very large arrays observed in nature (at all chromosomes) could only be generated when: i) longer crossover products repeatedly and fortuitously drift to fixation on all 23 chromosomes simultaneously, or ii) natural selection favors the large size of centromeric HOR arrays. The first explanation is statistically improbable and the second is insufficient to explain why the average size of HOR arrays far exceeds what is required for normal cellular functioning. The multimegabase size of human centromeric HOR arrays is all the more perplexing because tandem arrays are expected to be continually eroded by SSA repair of DSBs (Supplemental Figure 1; Ozenberger et al. 1991; Muchova et al. 2015; Bhargava et al. 2016; Warmerdam et al. 2016). So the observed large size of human HOR arrays is perplexing if crossover between sisters is the major factor generating length variation and homogenization at human HOR arrays. As an alternative to unequal crossovers, I searched for a molecular mechanism that could homogenize long HOR arrays and amplify them to a size of many megabases, despite no selection for this extreme size and despite their continual erosion by SSA repair of DSBs.

## An alternative to the unequal crossing over model

I began my search by looking for biochemically well-characterized long tandem repeats (on a size scale similar to centromeric HORs, i.e., with repeated units much longer than those found at telomeres and micro- and mini-satellites) that were known to be strongly homogenized and also capable of deterministic expansion over time. This immediately led me to rDNA tandem repeats which have been extensively studied at the molecular level in many model organisms –but especially budding yeast (reviewed in Kobayashi 2014). Unequal crossing over at rDNA repeats can lead to both deletions and insertions, and DNA double strand breaks (DSBs) repaired via single strand annealing (SSA) are expected to lead to small deletions of one repeat per DSB (Supplemental Figure S1; Muchova et al. 2015; Bhargava et al. 2016; Warmerdam et al. 2016). Although expected to be much rarer, repair of pairs of DSBs via **N**on-**H**omologous **E**nd-Joining (NHEJ) will also sometimes generate large deletions (Supplemental Figure S2). The length of a rDNA tandem repeat will shrink stochastically (when sampling error leads to the accumulation of more expansions than contractions from unequal crossing over) and deterministically (via SSA repair of DSBs [Ozenberger et al. 1991] and deletions via NHEJ repair of pairs of DSBs) in a cell lineage over time. The deterministic components make shrinkage inevitable over time. Such shrinkage could be prevented by natural selection against shorter repeat arrays, but nature has solved the problem via a molecular regeneration mechanism (using the BIR pathway) that is activated when repeat copy number within a repeat array is low (Supplemental Figure S3A; Kobayashi et al. 1998, Kobayashi et al. 2004; Kobayashi & Ganley 2005; reviewed in Kobayashi 2014).

The expansion of rDNA repeats is described in detail in Box 1. The key features are listed below and illustrated in Supplemental Figure S3:

i. Protein bound to rDNA causes replication fork-stalling, and some of these stalled forks result in fork-collapse.
ii. Fork-collapse generates a one-ended DSB yielding two strands of DNA: a partially replicated full-length sister chromatid and a truncated, partially replicated sister chromatid (Supplemental Figure S3A).
iii. The truncated chromatid can re-initiate DNA replication via the Break-Induce-Repair (BIR) pathway at multiple points of homology along the HOR array on the partially replicated sister chromatid: i) at the original point of breakage (in-register), or ii) upstream or downstream of this location (out-of-register).
iv. When sisters are tightly bound by dense cohesin, they have strongly constrained movement and replication is consistently re-initiated in-register.
v. When sisters are loosely bound by sparse cohesin, they have higher mobility and replication re-initiation can be out-of-register (up-stream or down-stream).
vi. For reasons not fully understood (but see Box 1), downstream re-initiation predominates, on average, leading to duplications of one or more repeat units (usually one).

Can we apply the information on rDNA expansion via fork-stalling/collapse and BIR replication (hereafter abbreviated ‘fork-stalling/collapse/BIR’) to human centromeres? In both the point centromeres of budding yeast (Greenfeder and Newlon 1992) and the regional centromeres of *Canida albicans* (Mitra et al. 2014) there is empirical evidence that kinetochore proteins bound to DNA (more specifically a subset of these proteins called the **C**onstitutively **C**entromere-**A**ssociated **N**etwork, CCAN) cause fork-stalling/ collapse, rather than it being caused by DNA secondary structure. Similarly, in budding yeast (point centromere; Sakuno & Watanabe 2009), fission yeast (regional centromere; Sakuno & Watanabe 2009), and *Arabidopsis* (regional centromere; Topp and Dawe 2006) there is empirical evidence that cohesin is absent or highly rarefied at the centric DNA that binds the kinetochore and has a unique epigenetic profile (‘centrochromatin,’ Sullivan and Karpen 2004), while flanking pericentric DNA (heterochromatin) has highly concentrated cohesin (see Figure 1).

Human centromeric repeats are constitutively bound to the group of 16 CCAN kinetochore proteins, and this protein binding would feasibly produce a barrier to DNA polymerase and therefore generate fork-stalling/collapse (Beuzer et al. 2014; McKinley and Cheeseman 2016). Fork-stalling/collapse may also be enriched at centromeric HOR arrays due to collisions between DNA and RNA polymerases because this DNA is actively transcribed during S phase (McNulty et al. 2017) and because, unlike rDNA repeats, these satellites are not established to contain replication fork barriers (RFBs) that block bidirectional fork progression.

### Box 1. Expansion of rDNA repeat arrays in budding yeast

The protein Fob-1 tightly binds rDNA. During DNA replication, this bound protein causes stalling of DNA polymerase (fork-stalling) that sometimes leads to fork-collapses (Mohanty and Bastia 2004). Fork-collapse produces a one-ended DSB (a truncated, partially replicated sister chromatid) and a full length partially replicated sister chromatid (Supplemental Figure S3A). The truncated chromatid is resected and homology search along the full-length, partially replicated sister chromatid reinitiates DNA replication via BIR: i) in-register with the original break, or ii) out-of-register at multiple points of homology within the tandem repeat array that occur both upstream (at un-replicated DNA that was ahead of the replication fork) and downstream of the location of the one-ended DSB (Supplemental Figure S3A). Downstream re-initiation of DNA replication causes one or more repeats to be copied twice during replication of the truncated strand, while upstream re-initiation leads to a loss of one or more repeat units (Supplemental Figure S3A).

When copy number of repeats within the rDNA array is high, there is reduced transcription of the bidirectional promoter (E-pro) within the rDNA spacer, which increases the epigenetic mark H3K56-ac (Kobayashi & Ganley 2005). This epi-mark is associated with rDNA (on newly synthesized sister chromatids behind the replication fork) that is tightly bound by cohesin during DNA replication. The dense cohesin-binding of sister chromatids behind the replication fork constrains the truncated chromatid (produced by fork collapse) to initiate BIR replication in-register on the full length sister chromatid: leading to no expansion or contraction of the tandem repeat. But when the rDNA repeat array is short, transcription of the bidirectional promoter (E-pro) within the rDNA spacer is increased, which decreases the H3K56-ac epi-mark, and cohesin is reduced. With reduced cohesin density, out-of-register initiation can occur because of the higher mobility of the truncated chromatid produced by fork-collapse: so repeat length can expand or contract (Supplemental Figure S3B). For reasons not fully understood, expansion predominates. This predominance might feasibly occur because downstream, newly replicated DNA has more open chromatin structure (increased DNAse 1 accessibility; Poot et al. 2005) which feasibly makes down-stream DNA more accessible to homology search. The key molecular features required for tandem repeat expansion are: i) protein-bound DNA (the predominant factor leading to fork-stalling and collapse, Mohanty and Bastia 2004; Beuzer et al. 2014), and ii) low levels of bound cohesin (that generates high mobility of the truncated chromatid and thereby permits out-of-register re-initiation of replication via BIR; Kobayashi & Ganley 2005).

Empirical evidence for fork-stalling/collapse in humans was provided by Crosetto et al. 2013. They showed that aphidicolin treatment of human HeLa cells (which slows the progression of DNA replication forks and amplifies DSBs in response to fork-stalling/collapse in regions that are natively high in fork-stalling) led to strong and significant enrichment of DSBs at centromeric repeats in the context of a genome-wide scan for DSBs. Additional evidence for fork-stalling/collapse at human centromeric DNA comes from the work of Aze et al. 2016. They compared the replication of human centromeric DNA to CG-matched control DNA with a *Xenopus laevis* egg extract assay. Compared to the control DNA, they found the replication of centromeric DNA to be slower, to recruit more DSB repair enzymes and MMR (mismatch repair) enzymes. All of these findings are consistent with fork-stalling and collapse during the replication of human centromeric DNA (Aze et al. 2016).

I have not found a study comparing cohesin levels at centric and pericentric regions of human HOR arrays (see Figure 1), but cohesin is expected to be concentrated within the pericentric heterochromatin because it is well established to recruit exceptionally dense cohesin (Sakuno & Watanabe 2009) due to its high concentration of the cohesin-loading H4K20 methyl transferase Suv4-20h2 (Hahn et al. 2013). This high concentration of cohesin is not expected within the centric core which is packaged as centrochromatin and lacks the H4K20 epigenetic mark. Another feature supporting highly rarefied cohesin within the centric core is the substantial separation of sister kinetochores (but not the flanking pericentric heterochromatic regions) when tension is applied to sister chromatids by the spindle fibers during mitotic metaphase (Tanaka 2010). In addition, the data described above from the more tractable species (budding yeast, fission yeast, and *Arabidopsis*) support the conclusion that cohesin is feasibly absent (or rarefied compared to the flanking percentric heterochromatin) at the kinetochore-recruiting regions (centric core) of human HOR arrays. BIR is the predominant repair mechanism following fork-collapse in yeast (reviewed in Anand et al. 2013) and there is extensive evidence that it is commonly used during the repair of collapsed replication forks in humans (reviewed in Leffak 2017; Sakofsky and Malkova 2017).

## Predictions and ramifications of fork-stalling and collapse followed by re-initiation of DNA replication via BIR

The studies described up to this point collectively support the conclusion that fork-stalling and collapse followed by re-initiation of DNA replication via BIR (fork-stalling/collapse/BIR) is feasibly an integral part of DNA replication at human centromeres. This phenomenon is expected to have widespread ramifications when combined with the diverse molecular information that has accumulated concerning the structure and functioning of human centromeres. The remainder of this paper explores these numerous predictions and ramifications (Table 1).

**Table 1.**
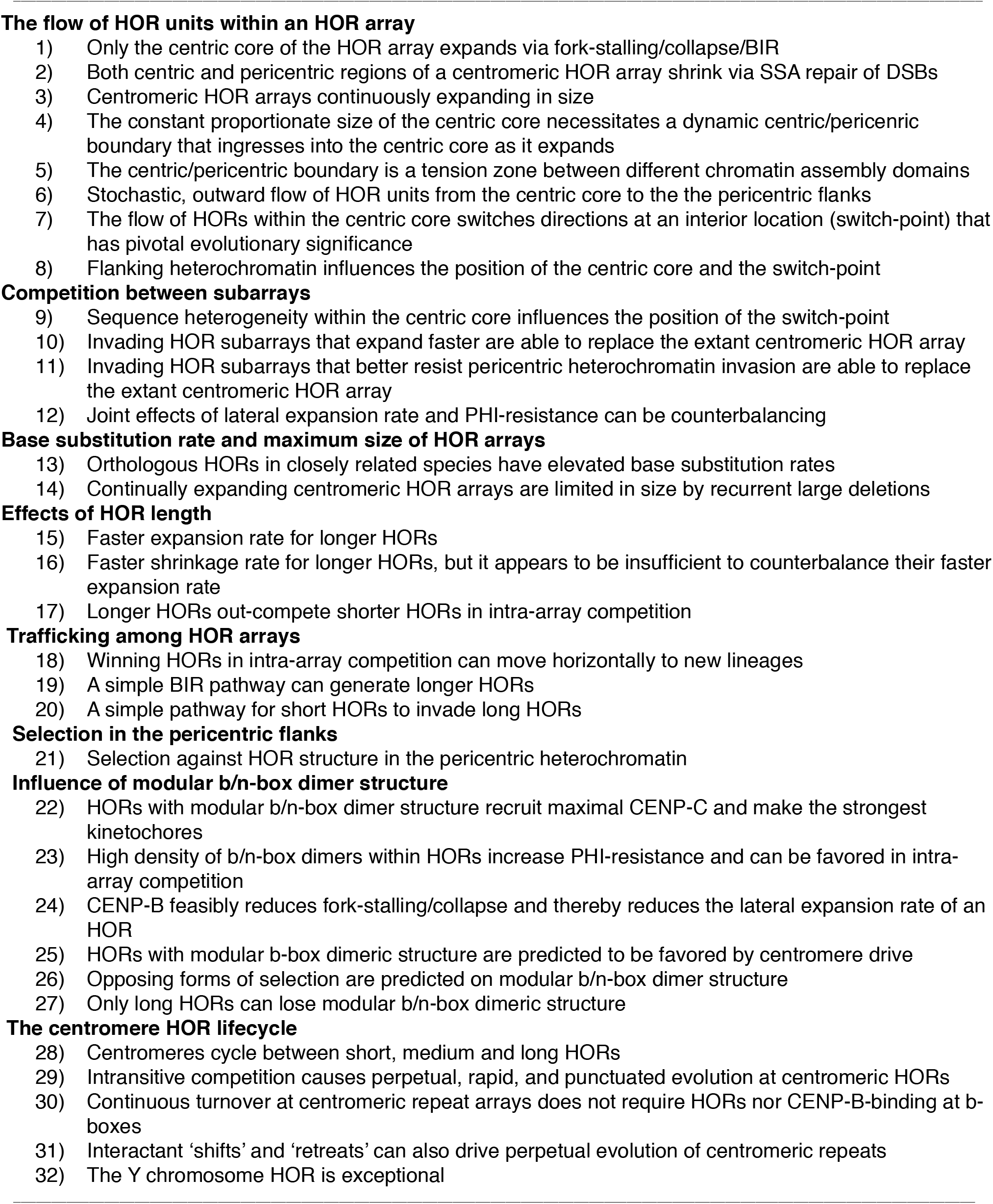
Diverse set of predictions and ramifications that stem from the operation of fork-stalling/collapse/ BIR at human centromeric HOR arrays when combined with their empirically established molecular characteristics.

The first group in Table 1 (the flow of HOR units within an HOR array) focuses on how the combination of fork-stalling/collapse/BIR and several empirically established molecular features of human centromeric repeats leads to the prediction of a stochastic, bidirectional flow of HOR units from a central position toward the two edges of their repeat array –and how the ‘switch-point’ where this flow reverses direction has a pivotal role in HOR evolution. The second group (competition between subarrays) concerns two phenotypes produced by HOR sequences (lateral expansion rate and PHI-resistance) that influence competition between subarrays within the same centromeric repeat array. The third group (base substitution rate and maximum size of HOR arrays) focuses on how recurrent fork-stalling/ collapse/BIR accelerates the rate of sequence divergence between species and also generates array sizes that are much larger than needed for cellular functioning. The fourth group (effect of HOR length) explores how the length of an HOR (number of monomers per repeat unit) influences: i) the rate of expansion and contraction of HOR arrays, and ii) competition between different subarrays within the same centromeric repeat array. The fifth group (trafficking among HOR arrays) focuses on the movement of HORs (and their subunits) between centromeric arrays on different homologs and different chromosomes. The sixth group (selection in the pericentric flanks) contrasts selection on sequence composition with in the pericentric heterochromatin compared to the centric centrochromatin. The penultimate group (the influence of modular b/n-box dimer structure) concerns the influence of b/n-box dimeric structure on: i) the recruitment of foundational centromeric proteins (CENP-A, −B, and −C), and ii) how these phenotypes influence fork-stalling/ collapse/BIR, the recruitment of new monomers during HOR evolution, and intra-array competition between subarrays. The final group (the centromere HOR lifecycle) focuses on i) a form of intransitive competition between HOR subarrays located on the same chromosome that leads to a cycle of perpetual and rapid turnover of HOR size and sequence, ii) how intra-array competition can occur in other species in the absence of both CENP-B and HOR structure, and iii) the special case of centromeric repeat evolution on the male-limited Y chromosome.

## The flow of HOR units within an HOR array

### Only the centric core of the HOR array is predicted to expand via fork-stalling/collapse/ BIR

Centromeric DNA sequences must provide two critical cellular functions during mitosis and meiosis: i) they must bind the kinetochore proteins that attach to spindle fibers, and ii) they must recruit dense cohesin clamps that keep sister chromatids attached until anaphase (mitosis) or anaphase-II (meiosis) (see Figure 1; Supplemental Figure S4). In the laboratory mouse (*Mus musculus domesticus*), these two functions are carried out by separate, neighboring arrays: the minor and major satellites that bind the kinetochore and recruit cohesin, respectively (Guenatri et al. 2004; Note: recent work suggests that the outer flanks of the minor satellite do not recruit kinetochore proteins and may recruit cohesin [Iwata-Otsubo et al. 2017]). In humans, a single HOR array is partitioned and used for both cellular functions (Figure 1). The HOR array partitions are: i) the centric core (centrochromatin, with unique epigenetic marks; Figure 3) that recruits the histone H3 variant CENP-A at a small –but 50-fold elevated proportion– of its nucleosomes (~4% during G1 and Searly stages of the cell cycle and ~2% during Slate, G2, and M stages; Bodor et al. 2014) and binds kinetochore proteins, and ii) the pericentric flanks (constitutive heterochromatin, with different, unique epigenetic marks; Figure 3) does not recruit elevated levels of CENP-A but recruits dense cohesin clamps (Sullivan et al. 2011; Hahn et al. 2013; Supplemental Figure S4). The active HOR itself is embedded within a region of pericentric heterochromatin (Figure 1) composed primarily of unordered monomeric repeats that are unrelated to it (and may also include smaller, inactive HOR arrays) (Shepelev et al. 2009).

**Figure 3.**
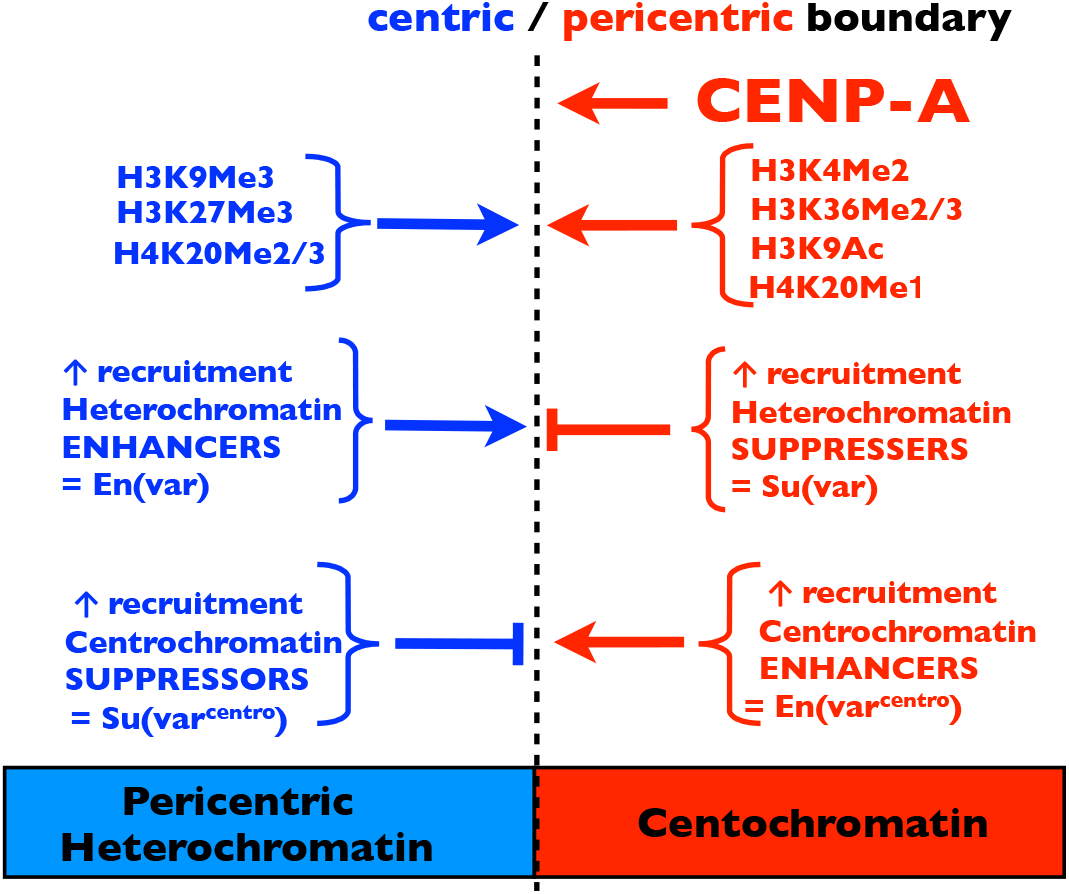
The boundary between a pericentric flank (heterochromatin) and the centric core (centrochromatin) of a centromeric HOR array is a tension zone between two alternative chromatin assembly domains. Centrochromatin is defined epigenetically by a 50-fold increase in the density of CENP-A nucleosomes (about 200 nucleosomes per centromere in cell cycle stage G1, which is about 4% of the nucleosomes of the centric core of a typical centromeric HOR array [Bodor et al. 2014]) and the H3 and H4 histone tail modifications shown in the figure in red (e.g., H3K4Me2). Pericentric heterochromatin is defined by the absence or low density of CENP-A nucleosomes and the heterochromatin-associated histone tail modifications shown in the figure in blue (e.g., H3K9Me3). Empirical data from position-effect variegation indicates that the recruitment and spreading of heterochromatin at the pericentric flank feasibly can be enhanced or suppressed by polygenic variation at many loci (collectively called En(var) and Su(var), respectively; Elgin and Reuter 2013). Although not empirically established, similar variation for the enhancement and suppression of centrochromatin recruitment and spreading (En[var^centro^], Su[var^centro^) is hypothesized to be present. Empirical data on the boundary between heterochromatin and euchromatin at pericentric inversions indicates that DNA flanking the centromeric HOR array may also influence this boundary (not shown on figure).

Within the two functional parts of the active HOR array, only the CENP-A-enriched, centric core is expected to be continually expanding in length via fork-stalling/collapse/BIR (as occurs in rDNA). Expansion is restricted to the centric core because: i) CCAN proteins bind only this region (predicted to cause fork-stalling/collapse/BIR), ii) RNA and DNA polymerase collisions feasibly occur in this region (predicted to cause fork-stalling/collapse/BIR), and iii) and cohesin is not concentrated in this region. In contrast to the centric core, the pericentric flanks of the array are not expected to expand because they do not bind CCAN proteins and recruit dense cohesin (Supplemental Figure S4). Even if some level of fork-stalling/collapse/BIR occurs within the pericentric flanks (e.g., due to collisions between RNA and DNA polymerases), the data from rDNA repeats in budding yeast (Kobayashi 2014) indicated that their high cohesin levels would be expected to suppress out-of-register BIR and hence prevent expansion.

### Both centric and pericentric regions of a centromeric HOR array are predicted to shrink via SSA repair of DSBs

I found no studies comparing repair of DSBs in centric and pericentric regions of human centromeric HOR arrays. Nonetheless, studies of other genomic regions in humans (George and Alani 2012; Geuting et al. 2013; van Sluis and McStay 2015; Bhargava et al. 2016) indicate that SSA repair of DSBs would be expected to cause deletions in tandemly repeated sequences located in both centric and pericentric regions (Supplemental Figure S1). This conclusion is supported by measures of DSB repair at mouse major (pericentric heterochromatin) and minor (centric centrochromatin) satellites (Tsouroula et al. 2016). This study found evidence for SSA repair of DSBs at both the minor and major satellites (although SSA repair was not the predominant repair pathway at either satellite), with a higher rate at the centrochromatin-containing minor satellite (occurs at G1, S, and G2 stages of cell cycle) compared to the pericentric, heterochromatic major satellite (occurs at S and G2 stages) (Tsouroula et al. 2016). Because the repair of DSBs is strongly influenced by chromatin structure (Mladenov et al. 2016) and because distinctive epigenetic marks for the centric and pericentric chromatin are similar between mice (Chan and Wong 2012), flies and humans (Sullivan & Karpen 2004), SSA repair of at least some DSBs (and the deletions they generate) feasibly occurs in humans across the entire centromeric HOR array.

### Centromeric HOR arrays are predicted to be continuously expanding in size

Empirical evidence reviewed in the previous two sections indicates that centromeric HOR arrays are both expanding (by fork-stalling/collapse/BIR in the centric core) and contracting (via SSA repair across the entire array). Contraction may be relatively rare relative to expansion, however, because data from mouse centromeres indicates that SSA repair was not the predominant DSB repair pathway. In addition, as described in an earlier section, HOR arrays are typically more than an order of magnitude larger than needed for normal cellular functioning (Lo et al. 1999; Yang et al. 2000; Okamoto et al. 2007) and sometimes achieve extreme sizes > 8 Mb (Miga et al. 2014). The observation that the typical size of centromeric HOR arrays at all chromosomes is far-larger-than-needed indicates a net excess of expansions over deletions at centromeric HOR arrays: causing them to be continually expanding. In a later section I consider how genetic drift of infrequent mega-deletions would be expected to limit the maximum size of persistently expanding centromeric HOR arrays.

### The constant proportionate size of the centric core necessitates a dynamic centric/ pericentric boundary that ingresses into the centric core as it expands

A collection of observations in humans and mice indicates that the boundary between the centric core and the pericentric flanks of an active HOR array is in a state of flux during array expansion (and contraction via mega-deletions, as discussed later), and that CENP-A concentration within the centric core influences this boundary. The centric core contains an average of about 2-4% nucleosomes that have CENP-A substituted for histone H3 –which is a 50-fold enrichment compared to other genomic regions (Bodor et al. 2014). Stretched chromosome studies indicate the centric core is a contiguous subset of an HOR array (Zeng et al. 2004; Lam et al. 2006; Mravinac et al. 2009) and that its proportionate size (about one third of the total array) is approximately constant across HOR arrays of highly different sequence (X vs. Y HOR arrays) and vastly different sizes (0.2 − 4.4 Mb; Sullivan et al. 2011; Ross et al. 2016). CENP-A was also detected along only a contiguous subregion of the minor satellite of the mouse (about a fifth; Iwata-Otsubo et al. 2017). In human cells, an increase in cellular CENP-A concentration leads to increased CENP-A deposition at centromeres (Bodor et al. 2014) and a corresponding expansion of the centric core region to cover a higher proportion of the HOR array (Sullivan et al. 2011). These observations indicate that the centric/pericentric boundary expands and contracts in response to changes in both CENP-A abundance and the size of the HOR array.

Because: i) only the centric core is predicted to expand, via fork-stalling/collapse/BIR, and ii) the centric core’s proportionate size remains approximately constant (about a third of the total HOR array), the centric/pericentric boundary is predicted to continually ingress into the centric core as it alone expands: causing 2/3^rds^ of each unit of its expansion to be moved out at its edges into the pericentric flanks and a net one third of its expansion to be retained within the centric core. This inward invasion of the pericentric flanks is feasibly caused by a reduced density of CENP-A within the centric core as it expands (Sullivan et al. 2011). The reduction occurs because all centromeres contain a similar amount of of CENP-A (~200 molecules in G2 and ~400 molecules in G1), irrespective of the size of their HOR arrays (Bodor et al. 2014): so expansion of the centric core must reduce the concentration of CENP-A per unit DNA. In Supplemental Figure S5, I propose a simple mechanism leading to the observed constant proportionate size of the centric core.

### The centric/pericentric boundary is predicted to be a tension zone between different chromatin assembly domains

Because the same centromeric HOR array is partitioned into pericentric flanks (heterochromatin) and a centric core (centrochromatin), and the relative size of these compartments (2 pericentric: 1 centric) remains stable over vastly different array sizes, an insulator sequence separating the compartments (as occurs in fission yeast; Scott et al. 2006) is almost certainly absent. Sullivan et al. (2016) examined a naturally occurring deletion of human chromosome 17 that removed the heterochromatic boundary on one side of the centromere. The deletion removed: i) about three fourths of the active centromeric HOR array, ii) all of the flanking pericentric heterochromatin (composed of an inactive flanking HOR array and unordered monomeric DNA), and iii) about 10Mb of the euchromatic arm (q-arm side). About 45% of the centrochromatin (identified by its elevated concentration of CENP-A) spread from the remaining HOR array and penetrated ~ 300 kb into the newly adjacent euchromatin. This finding indicates that centrochromatin can spread far into adjacent, non-heterochromatic DNA. Within the newly adjacent euchromatin that was penetrated by centrochromatin, strong heterogeneity of the level of CENP-A density was observed: indicating that local sequence could quantitatively influence centrochromatin spreading.

This spreading of centrochromatin into euchromatin resembles the spreading of pericentric heterochromatin into euchromatin after a pericentric inversion newly juxtapositions euchromatin and heterochromatin without an insulator (which may simply be a long stretch of DNA without any special sequence) separating them. The spreading of heterochromatin into adjacent euchromatin silences embedded euchromatic genes: a phenomenon called **P**osition-**E**ffect **V**ariegation (PEV). The vast body of work on PEV is summarized in a detailed review by Elgin and Reuter (2013), and I summarize the information relevant to centric/ pericentric boundary in the following paragraph.

At the boundary of a pericentric inversion, the level of heterochromatin spreading into adjacent euchromatin varies between cell lineages within a tissue –leading to a variegated phenotype produced by a mosaic of tissue patches with active or heterochromatin-suppressed genes. Mutagenesis studies indicate that heterochromatin spreading is influenced by the level of expression of at least 150 genes (some of which directly participate in heterochromatin assembly) which either increase gene silencing (enhancers of variegation, En[var]) or reduce it

(suppressors of variegation, Su[var]). Addition or deletion of large blocks of heterochromatin (e.g., adding or deleting a Y chromosome) strongly suppresses or enhances PEV silencing, respectively: indicating that dilution or enrichment of heterochromatin assembly factors strongly influence heterochromatin spreading. On average, closer proximity of a gene to the heterochromatin boundary produces a higher density of the H3K9Me3 heterochromatic epigenetic mark and a greater level of gene inactivation: but some genes closer to the breakpoint exhibit less silencing than other, more-distal genes. Also, the same gene engineered to have different promoters can have strongly different sensitivities to PEV silencing. These observations collectively indicate that both the concentration of trans-acting heterochromatin assembly factors and cis-acting DNA sequence can strongly influence the level of heterochromatin spreading. Although I have found no genetic screens for suppressors and enhancers of centrochromatin spreading, the large number of loci that influence pericentric heterochromatin spreading (i.e., influencing PEV), makes it plausible that there are also enhancers and suppressors of centrochromatin spreading. I will denote this hypothesized genetic variation that positively and negatively influences the spreading of centrochromatin as En(var^centro^) and Su(var^centro^), respectively.

Lack of an insulator sequence separating the centric core (centrochromatin) from its pericentric flanks (heterochromatin) is expected to generate a dynamic boundary (Figure 3). The CENP-A-enriched centric core is expected to spread centrochromatin (with its characteristic epigenetic marks; Figure 3) outward and the pericentric flanks are expected to spread heterochromatin (with its characteristic epigenetic marks; Figure 3) inward at the boundary between the two domains. The relative strength of these two mutually opposing remodeling domains at the boundary between pericentric heterochromatin and centrochromatin of the centric core is expected to be influenced by any sequence-specific effects of the HOR in recruiting: i) CENP-A, ii) centrochromatin-specific histone tail modification and chromatin assembly factors, iii) heterochromatin-specific histone tail modifications and chromatin assembly factors, iv) suppressors and enhancers of heterochromatin spreading (En[var] and Su[var]), and possibly v) suppressors and enhancers of centrochromatin spreading (En(var^centro^) and Su(var^centro^). These influences are shown collectively in Figure 3. I will use the term **P**ericentric **H**eterochromatin **I**nvasion **resistance** (PHI-resistance) to describe the degree to which the sequence of an HOR impedes the invasion of the centric/pericentric boundary as the centric core expands. As described in later sections, PHI-resistance will have important evolutionary consequences when sequence heterogeneity causes it to differ between the two centric/pericentric boundaries of an HOR array.

### Stochastic, outward flow of HOR units is predicted from the centric core to the the pericentric flanks

Consider an HOR array that is expanding via fork-stalling/collapse/BIR within its centric core (Figure 4). As the centric core expands, the boundaries between the centric core and its pericentric flanks are expected to move inward: causing expansion within the centric core to produce growth in both the centric core and the pericentric flanks. As described above, this inward movement of the centric/pericentric boundaries must occur if the relative size of the centric core remains constant (~ one third of the total array) as the array expands in size. As a corollary during array expansion: i) HOR units within the centric core are expected to be continually pushed outward when new HOR units are formed (by fork-stalling/ collapse/BIR) in more central locations within this region, while ii) the centric/pericentric boundary moves inward in response to the expansion, feasibly due to dilution of CENP-A within the centric core (Figure 4). These two features generate a stochastic, outward flow of HOR units from the centric core (centrochromatin) where they are tandemly replicated (‘born’ via fork-stalling/collapse/BIR; Supplemental Figure S3) into the pericentric flanks (heterochromatin) where they are no longer tandemly replicated but will ultimately be deleted (‘die’) by recurrent SSA repair of DSBs (Supplemental Figure S1) and rarer deletions from NHEJ repair of pairs of DSBs (Supplemental Figure S2).

**Figure 4.**
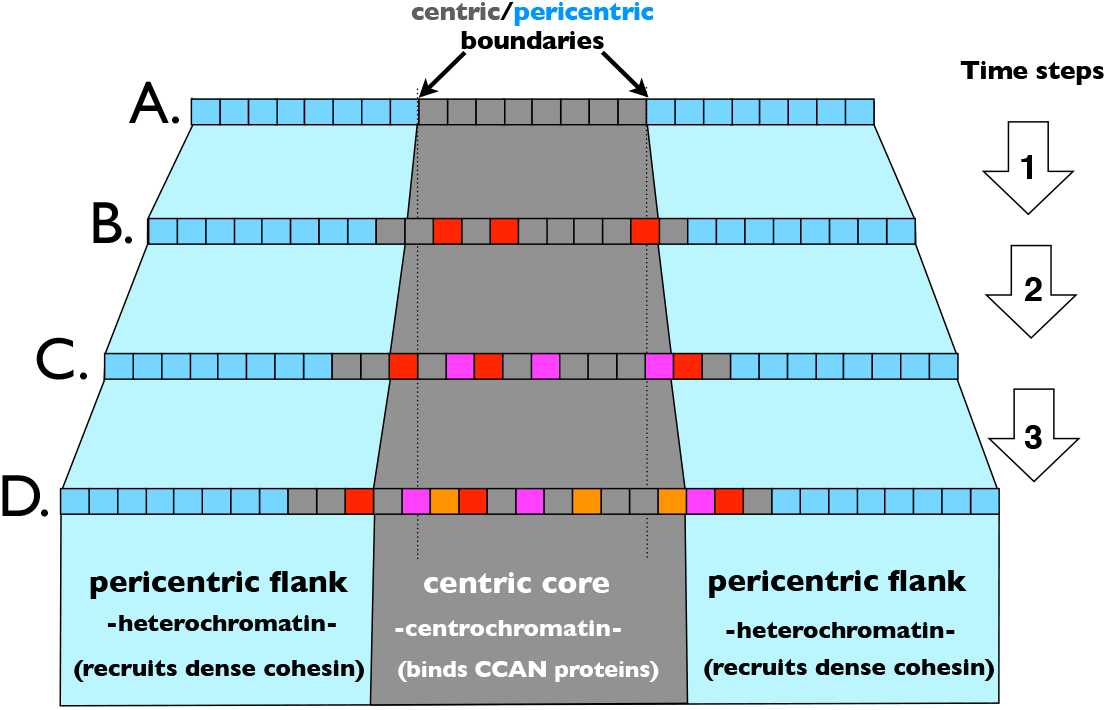
The hypothesized net, stochastic outward movement of repeat elements from the centric core (where they are ‘born’ via fork-stalling/collapse/BIR; Supplemental Figure S3) into the pericentric flanks (heterochromatin) where they will ultimately ‘die’ by being deleted by recurrent SSA repair of DSBs and rarer deletions from NHEJ repair of pairs of DSBs (see Supplemental Figure S2). **A.** Each box represents an HOR unit. One third of the units are packaged as centrochromatin within the centric core and one third each are packaged as pericentric heterochromatin at the two flanks. **B.** Over time, three new repeat units (red) are generated at random positions only within the centric core region of the HOR via fork-stalling/collapse/BIR (for simplicity, rarer deletion events are ignored). This expansion causes each pericentric flank to move inward (feasibly because it dilutes the CENP-A concentration within the centric core; Figure 3), keeping the proportions of the HOR array at ~1/3 centric core and ~2/3 pericentric flanks. As a consequence, two repeats (one on each side) have been moved from the centric core into the pericentric flanks. **C.** Over additional time, three more repeat elements (purple) are randomly added to the centric core via fork-stalling/collapse/BIR. In response, the centric/pericentric boundaries move inward, and two more repeats (one on each side) that originated (i.e., were ‘born’) in the centric core are moved into the pericentric flanks, where over time they will gradually be lost (‘die’) via deletions (e.g., by SSA repair of DSBs). **D.** With more time, three more repeats (orange) are added to the centric core via fork-stalling/collaps/BIR, leading to the movement of two more repeat units (one on each side) to the pericentric flanks. The restriction of fork-stalling/ collapse/BIR to the centric core, when coupled with the inward migration of the centric/pericentric boundary, generates a net flow of repeat elements from the centric core to the pericentric flanks.

### The flow of HORs within the centric core is predicted to switch directions at an interior location (switch-point) that has pivotal evolutionary significance

Consider a centromeric array with a homogeneous HOR sequence across its length. During array expansion via fork-stalling/ collapse/BIR within the centric core, the direction of outward flow of HORs toward the pericentric flanks (Figure 4) depends on position. For reference, let the ‘right’ side of the array be the one closest to the long-arm (q-arm) of a chromosome. HORs within the centric core that are to the right-of-center experience more lateral expansion on their left side compared to their right side, on average, so their direction of flow is to the right-of-center (and vice versa for HORs located to the left-of-center). At the midpoint of this homogeneous centric core, the average outward flow of HORs reverses direction. I will refer to this point of reversed flow as the ‘switch-point’ (Figure 5).

**Figure 5.**
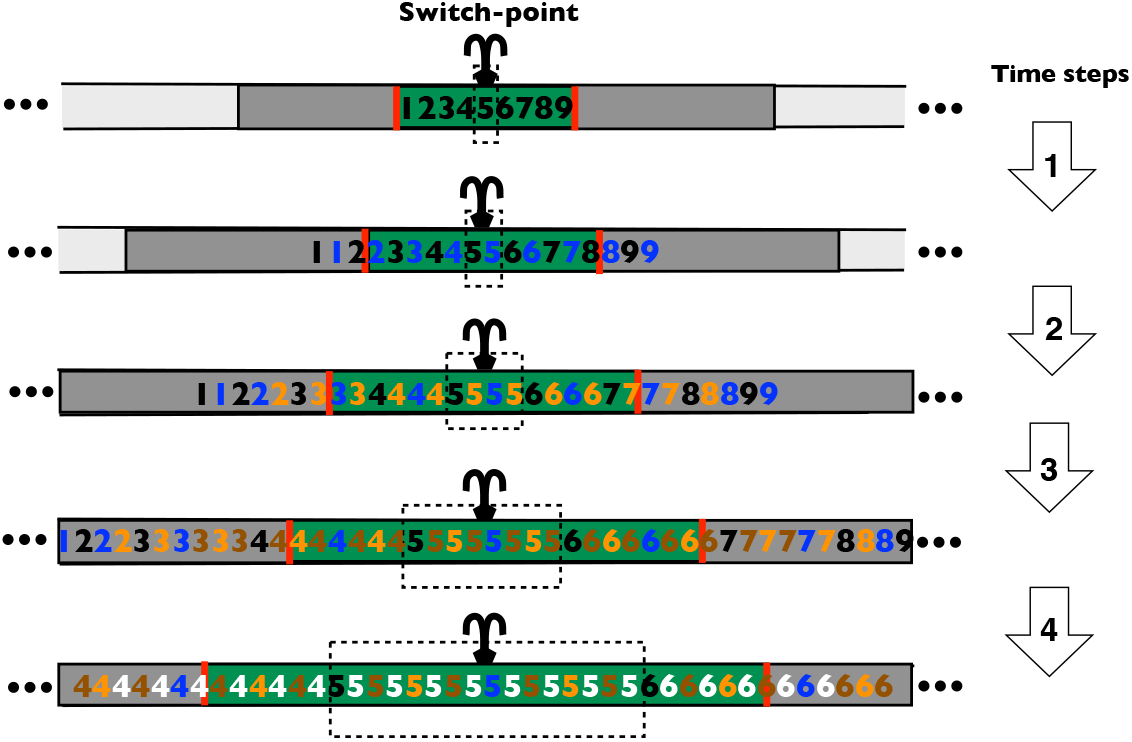
The outward flow of HOR elements (from the centric core to the pericentric flanks) reverses direction at a position called the ‘switch-point’ (indicated by a fountain icon). Bidirectional flow from this switch-point causes new sequences (e.g., point mutations) that originate near here –and that eventually reside in tandem copies than span the switch-point– to spread laterally toward both sides of the array, and eventually encompass the entire centric core and ultimately the entire array once recurrent deletion pressure has removed all older HOR elements. The centric core (centrochromatin, green background) of the HOR array is surrounded by the pericentric flanks (heterochromatin, dark grey background) and the boundaries between these two domains is shown by a red line. For clarity in this illustrative example, there are only 9 copies of the HOR within the centric core (numerals 1-9) and 9 + 9 = 18 in the pericentric flanks (to improve clarity, numerals for these initial 9 + 9 flanking copies are not shown throughout the figure). Fork-stalling/collapse/BIR tandemly duplicates array elements within the centric core, but not those within the pericentric flanks. In this deterministic, illustrative example, fork-stalling/ collapse/BIR within the centric core is assumed to simultaneously replicate each of its HOR elements in tandem one time per time step. After the first time step, 9 new HOR elements are generated in tandem (shown by blue numerals). These new tandem repeats would cause the centric core to expand by 9 repeat units but most of this gain is lost at its margins because the centric/pericentric boundary moves inward as the centric core expands (to maintain a proportional size of the centric core of one third). This inward migration of the centric/pericentric boundaries causes 6 (67%) of the 9 units of expansion to exit the centric core at its outer edges (3 units [33%] on each side) and become part of the pericentric flanks, and 3 units (33%) to remain within the centric core. New HOR elements generated by each time step are shown by a new numeral color. The combination of i) expansion of the centric core, and ii) inward movement of the pericentric flanks, generates a net flow of HOR elements from the middle of the centric core (the switch-point, where the direction of flow reverses) toward the pericentric flanks. Over time, the copies of the sequence that spans the switch-point increase and gradually spread to a progressively larger proportion of the centric core (dashed rectangles). The same flow process occurs, on average, when fork-stalling/collapse/BIR incrementally generates tandem copies of individual HOR elements at random locations, but the position of the switch-point is a random variable centered at the middle of the centric core.

Deletions due to SSA repair of DSBs will contribute to a flow of HORs in a direction that also depends on position. On average, HORs to the right-of-center of the centric core experience more deletions to their left side compared to their right side: so their direction of flow (due to deletions alone) is leftward (inward), toward the center. The same logic causes an average rightward (inward) flow on the left side. Assuming that expansions substantially exceed deletions (as described previously), there will be a net outward flow of HORs despite the inward flow generated by less common deletions.

Because the breakpoints generated by replication fork-collapse within the centric core are expected to be random (Figure 4) rather than deterministic (as are deletions due to SSA repair of DSBs), the position of the switch-point is probabilistic rather than deterministic. For example, stochastic fluctuations in the location of expansions and deletions might cause a repeat element slightly to the right of the midpoint of the centric core to ultimately exit the centric core on the left side. However, once a repeat element is sufficiently far from the mid-point, such a reversal will not occur. So the switch-point is actually a narrow region, within which the direction of flow of an HOR element is ambiguous. Throughout the remainder of this paper I will define the switch-point to be the narrow band with ambiguous direction-of-flow of HOR elements that separates the two sides of the centric core that flow in opposite directions.

The HOR element with copies spanning the switch-point (i.e., has copies on both sides) has important evolutionary significance (Figure 5). Any novel mutation in this HOR element will spread outward in both directions and eventually be included in all HOR elements within the centric core, and ultimately the entire HOR array (after recurrent deletion pressure removes all older HOR elements within the pericentric flanks). As a consequence, any sequence that makes its way to (and spans) the switch-point is expected to eventually spread to the entire centromeric array.

In later sections I will describe how phenotypes produced by the sequences of different HORs within the same active centromeric repeat array can give them a competitive advantage that allows them to spread and eventually encompass the entire array. Competition between different HOR subarrays within the same active centromeric array represents a new form of intragenomic reproductive competition that is predicted to occur within tandem centromeric repeats (a form of molecular drive; Dover 1982). This intra-array competition favors HORs that produce phenotypes that enable them to ‘capture’ (to make their way into and span) the switch-point.

### Flanking heterochromatin is predicted to influence the position of the centric core and the switch-point

The centric core is not necessarily expected to be located in the center of the centromeric HOR array, nor is the switch-point expected to always be located in the center of the centric core. Sequence heterogeneity that influences local lateral expansion rate (due to fork-stalling/collapse/BIR) and/or the strengths of centrochromatin and heterochromatin spreading (PHI-resistance) is predicted to influence the position of both the centric core within the HOR array and the switch-point within the centric core, as described later in the following four sections.

An additional factor that can feasibly influence the positions of both the centric core and its switch-point is an asymmetrical influence of neighboring heterochromatin (on either side of the centromeric HOR array) on the level of heterochromatin spreading into the two sides of the centric core as it expands (due to fork-stalling/collapse/BIR). Studies of PEV indicate that the spreading of heterochromatin into euchromatin can be strongly influenced by the amount (and possibly the composition) of flanking pericentric heterochromatin, and also the three d i m e n si on al d i s tan ce fro m n u cl e ar ‘heterochromatin compartments’ that are expected to contain a higher concentration of factors used to assemble heterochromatin (e.g., HP1alpha and H3K9-specific methyltransferases) (reviewed in Elgin and Reuter 2013). I will refer to these influences of heterochromatin near the centromeric HOR array as ‘neighboring heterochromatin effects’.

If heterochromatin spreading is stronger on one side of the centric core (due to neighboring heterochromatin effects), this asymmetry is predicted to displace the centric core toward the opposite side of the array. For example, suppose that resistance to the inward spread of pericentric heterochromatin was 5x weaker on one side (e.g., the p-arm side) of the centric core due to a larger block of neighboring heterochromatin on this side. In this case, as the centric core expands (due to fork-stalling/collapse/BIR) and the centric/ pericentric boundary ingresses (maintaining a ratio of approximately 33% centric core to 67% pericentric flanks), the ingression will be asymmetrical: for every one unit (e.g., a monomer) moved into the flanking percentric heterochromatin on the q-arm side there will be five units moved onto the opposite flank (the p-arm side). Put another way, as the centric core expands, its DNA is flowing out into the pericentric flanks 5x faster on the p-arm side. Over time, this asymmetry will generate a centric core that is displaced toward the q-arm side of the centromeric HOR array. Because the repeat units are moving into the pericentric flanks faster on the p-side compared to the q-side, the switch-point will be displaced from the center of the centric core toward the q-arm side.

There is empirical evidence for a non-central position of the centric core. Motivated by earlier studies that found the centric core on the X chromosome to be consistently biased toward the p-side (i.e., the short-arm side) of the HOR array (Schueler et al. 2001; Spence et al. 2002), detailed measurements by Ross et al. 2016 found that the position of the X chromosomes’ centric core was strongly biased toward the p-arm side of the HOR array in both of two unrelated cell lines. Although less rigorously documented, the position of the centric core of the Y chromosome also has been reported to be biased (across unrelated cell lines) toward the p-arm side of the HOR array (Floridia et al. 2000). These observations indicate that sequence variation across an HOR array and/or an influence of flanking DNA (e.g., a neighboring heterochromatin effect) may strongly influence the position of the centric core (Ross et al. 2016). Studies of sequence variation among different copies of the same monomer element (e.g., differing sequences of the i^th^ monomer of the X chromosome’s 12 monomer HOR) on both the X and Y chromosomes found very low divergence (Durfy and Willard 1989; Jain et al. 2018). This high sequence uniformity observed at both HOR arrays indicates that neighboring heterochromatin effects, rather than sequence variation within the HOR array, are more likely responsible for the observed strong bias in the position of the centric core toward one side of the array on both of these sex chromosomes.

## Competition between subarrays

### Sequence heterogeneity within the centric core is predicted to influence the position of the switch-point

When the HOR sequence of the centric core is uniform across its length (and there are no neighboring heterochromatin effects) the p-arm (short-arm) and q-arm (long arm) halves of the centric core are expected to have, on average: i) equal lateral expansion rates, and ii) equal rates of outflow from the centric core into the pericentric flanks (as the centric core expands and the centric/pericentric boundaries ingress). These two uniformities generate a switch-point that is positioned at the center of the centric core (Figure 6A). Some sequence non-uniformities, however, are expected to move the position of the switch-point.

**Figure 6.**
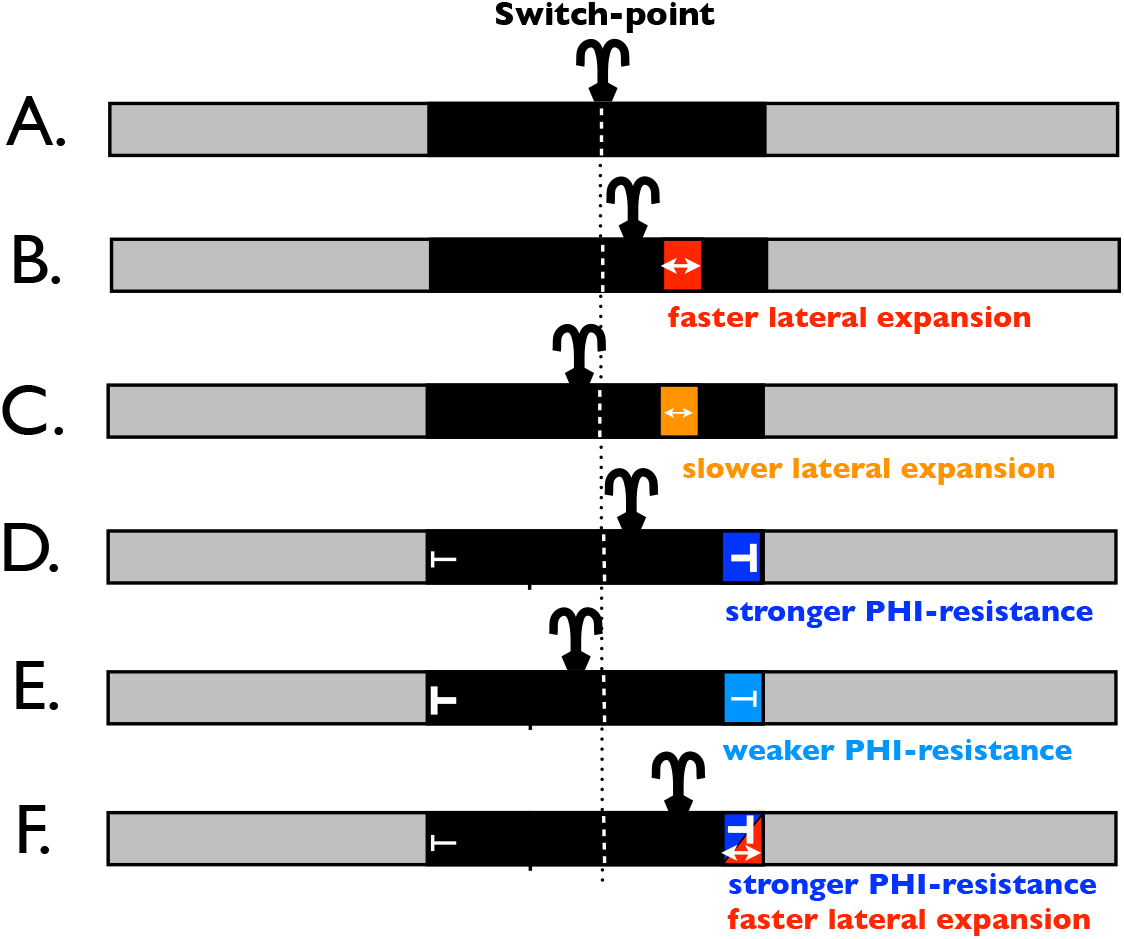
A new subarray recruited to a centromeric HOR array can influence the position of the switch-point. **A.** A centromeric HOR array composed of pericentric flanks (grey) and a centric core (black). The array contains no substantial sequence heterogeneity and no substantial asymmetric neighboring heterochromatin effects. This uniformity generates a switching-point (where outward flow of HOR elements reverses direction, on average) in the center of the array. **B.** The array in panel-A has been invaded (e.g., via transposition) by a small, new HOR subarray that has a faster lateral expansion rate. The presence of the faster subarray causes the right half of the centric core to expand faster than its left half, which moves the switch-point to the right, i.e., toward the side containing the faster expanding subarray. **C.** Same as panel-B except here the new subarray has a slower lateral expansion rate, causing the right half of the array to have slower lateral expansion rate. This asymmetry shifts the position of the switch-point toward the opposite side of the centric core (to the left in the figure). **D.** Same as panel-B except the new subarray causes stronger resistance to the invasion of pericentric heterochromatin into the centric core as it expands (stronger PHI-resistance). This phenotype is only expressed when the subarray is at or near a centric/pericentric boundary. As the centric core expands due to fork-stalling/collapse/BIR, two thirds of the expansion of the centric core is pushed into the pericentric flanks (outflow). The stronger PHI-resistance of the new subarray causes more of the outflow to occur on the opposite (left) side of the centric core. This asymmetry in outflow moves the switch-point toward the side of the centric core with the slower outflow rate, i.e., to the side containing the new subarray. **E.** Same as panel-D but the new subarray causes weaker PHI-resistance, which moves the switch-point to the left, i.e., away from the side containing the new subarray with weaker PHI-resistance. **F.** The new HOR array could feasibly influence both PHI-resistance and lateral expansion rate in a manner that is reinforcing (as shown in this example) or opposing (not illustrated).

Consider a centromeric HOR array with high sequence homogeneity along its length (and with no neighboring heterochromatin effects) that is “invaded” (to the right of center in Figure 6B) by a short subarray that has a different lateral expansion rate. I will use the term “invade” to refer to the establishment of a new subarray with a different sequence that originates by mutation, transposition, or a combination of these two processes. I will discuss this invasion process in a later section. If the expansion rate of the subarray is higher (Figure 6B), the side of the centric core containing the subarray (right side in Figure 6B) will have a higher lateral expansion rate. This asymmetry causes the switch-point to be moved toward the side containing the new subarray (Figure 6B). The position of the switch-point changes because the expansion is faster on the right half of the centric core (containing the new subarray) compared to the left half: so the point of equal expansion on each side (the switch-point) is shifted toward the side containing the faster expanding subarray. By the same logic, a new subarray that has a lower lateral expansion rate will move the switch-point toward the side that does not contain it (Figure 6C).

Next consider a newly recruited subarray that influences PHI-resistance (the capacity to i mpede the invasion of pericentric heterochromatin into the centric core at the centric/pericentric boundary as the centric core expands). This phenotype is not expected to be expressed until the subarray is pushed to (or near) a centric/pericentric boundary. First consider the case where the subarray has higher PHI-resistance compared to the larger array that it invaded (5-times higher; Figure 6D). As the centric core continually expands via fork-stalling/ collapse/BIR, two-thirds of the expansion is pushed into the pericentric flanks as the centric/ pericentric boundary ingresses into the centric core: which I will refer to as the ‘outflow.’ A new subarray with higher PHI-resistance will slow the outflow rate on the side of the centric core where it resided (right-side in Figure 6D). This asymmetry changes the switching-point at which the average outward flow of HORs reverses direction, because at the center of the centric core the net flow of HORs is leftward (toward the side with lower PHI-resistance). As a consequence, the switch-point is moved from the middle of the centric core toward the side that contains the new subarray with stronger PHI-resistance (Figure 6D). Because the outflow rate is 1-unit-right to 5-units-left, the switch-point will be positioned 100[1/(1+5)]% of the way from the right side of the centric core. By the same logic, the switch-point will be moved away from the side of the centric core containing a new subarray that has weaker PHI-resistance (Figure 6E).

### Invading HOR subarrays that expand faster are predicted to be able to replace the extant centromeric HOR array

Consider a homogeneous centromeric HOR array that is invaded by a short piece of a different HOR (the new subarray) that has a faster lateral expansion rate (e.g., it produces more fork-stalling/collapse/BIR per cell cycle)(Figure 7). For simplicity, further assume that there are no asymmetrical neighboring heterochromatin effects that influence the rate of pericentric heterochromatin spreading. For these reasons, the switch-point is located in the center of the centric core prior to the invasion of the new subarray (Figure 7A). If the new subarray invades either pericentric flank, it will not expand via fork-stalling/collaps/BIR and it will eventually be lost via recurrent deletion pressure (e.g., by SSA repair of DSBs). If the new subarray invades within the centric core but far to one side, it rapidly will be pushed into the flanking pericentric heterochromatin and have the same fate as if it began in this position. But if the new subarray invades the centric core sufficiently close to the switch-point, it will eventually spread to the entire HOR array.

**Figure 7.**
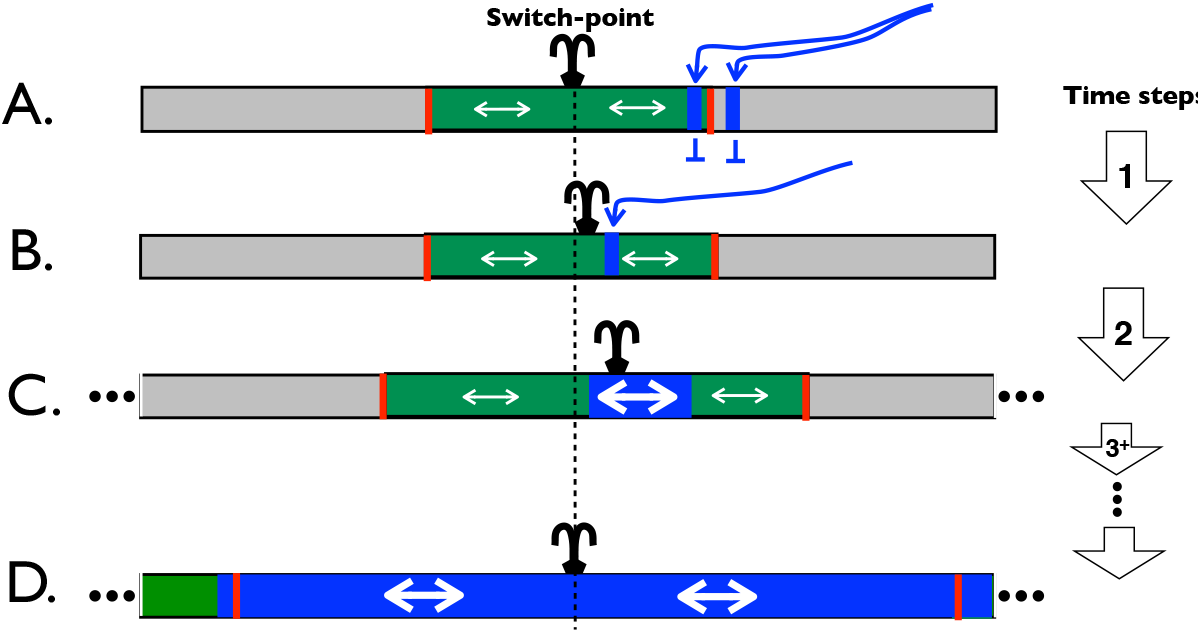
Invasion of a new HOR subarray with a faster lateral expansion rate (blue rectangle, with thicker arrow) into an established centromeric HOR array (combined green [centric core] and grey [pericentric flanks] rectangles). **A.** If the new subarray enters the established HOR array in one of the pericentric flanks, it will not expand via fork-stalling/collapse/BIR and eventually be lost due to recurrent deletion pressure. The same outcome occurs when the subarray enters the centric core -but too close to one of the centric/pericentric boundaries (red lines) because it will rapidly be pushed into a pericentric flank as the centric core expands and the pericentric flank encroaches into the centric core. **B.** When the subarray invades sufficiently close to the switch-point (where the net outward flow of HOR elements reverses direction), it can eventually expand to encompass the entire centric core, and ultimately the entire HOR array. The presence of the small, newly invading subarray immediately causes the right side of the centric core to expand faster, on average, than the left side -and this asymmetry moves the switch-point slightly to the right of the middle of the centric core. As the relative size of the faster-expanding subarray increases with time, the switch-point moves progressively further to the right (toward the new array). **C.** If the subarray enters sufficiently close to the switch-point, it will encompass the switch-point before being pushed off the right side of the centric core. At this point the new HOR subarray will be pushed in both directions as the centric core expands, and eventually spread to the entire centric core. **D.** Once recurrent deletion pressure removes all of the original array from the pericentric flanks (where dense cohesin clamps associated with pericentric heterochromatin prevent lateral expansion via fork-stalling/collapse/BIR), the new HOR with faster lateral expansion rate will encompass the entire centromeric array.

For example, suppose that the new subarray invades on the q-arm side of the centric core (the right side in Figure 7) but not too close to the edge. The presence of the new, small subarray (that has a faster lateral expansion rate) will make the right half of the centric core expand slightly faster than the left half, and this asymmetry will move the switch-point slightly to the right. As the new subarray expands in size, the asymmetry between the expansion rate of the right and left halves of the array will increase and the switch-point will move progressively closer to the subarray (Figure 7B,C). If the switch-point enters the subarray (deep enough to have its HOR copies span both of its sides) before it is completely pushed into the flanking pericentric heterochromatin, the new subarrays will begin expanding in both directions and eventually spread to the entire centric core. The same logic applies to the case where the switch-point is initially displaced from the center due to neighboring heterochromatin effects on the rate of pericentric heterochromatin spreading: if the faster spreading subarray starts close enough to the switch-point so that it spans it (captures it) before being pushed out of the centric core, the new, faster-expanding subarray is predicted to eventually replace the sequence of the entire centromeric array.

Competition between different HOR subarrays within the same active centromeric array represents a new form of intragenomic reproductive competition that is predicted to occur within tandem centromeric repeats (a form of molecular drive; Dover 1982). It favors the HOR with the higher lateral expansion rate, but with a caveat: starting position matters, because faster-expanding HORs only replace slower-expanding ones when they invade sufficiently near the switch-point of the centric core (i.e., not too close to the centric/pericentric boundary nor within the pericentric flanks).

### Invading HOR subarrays that better resist pericentric heterochromatin invasion are predicted to be able to replace the extant centromeric HOR array

As described previoulsly, to maintain a constant size of the centric core of about one third of the HOR array, the centric/pericentric boundaries on either side of the centric core must continually move inward (ingress) as the centric core expands. As a consequence, asymmetries between the two flanks of the centric core in resistance to pericentric heterochromatin invasion (PHI-resistance) can influence the position of the switch-point, and ultimately the fate of a newly recruited subarray.

Again, consider a homogeneous centromeric HOR array that is invaded by a short subarray of a new HOR, but in this case the new subarray has the same lateral expansion rate as the original array but has higher PHI-resistance (five times higher; Figure 8A,B). This higher resistance causes a slower ingression of pericentric heterochomatin into the centric core as it expands due to fork-stalling/collapse/BIR – compared to the opposite side of the centric core. For simplicity, further assume that there are no neighboring heterochromatin effects that influence the rate of pericentric heterochromatin spreading, so that before the new subarray invaded, the switch-point was located in the center of the centric core (Figure 8A). The new subarray can replace the original array when it invades close enough to the switch-point: where close enough depends on how much better the new sequence is at resisting being pushed into the pericentric heterochromatin.

**Figure 8.**
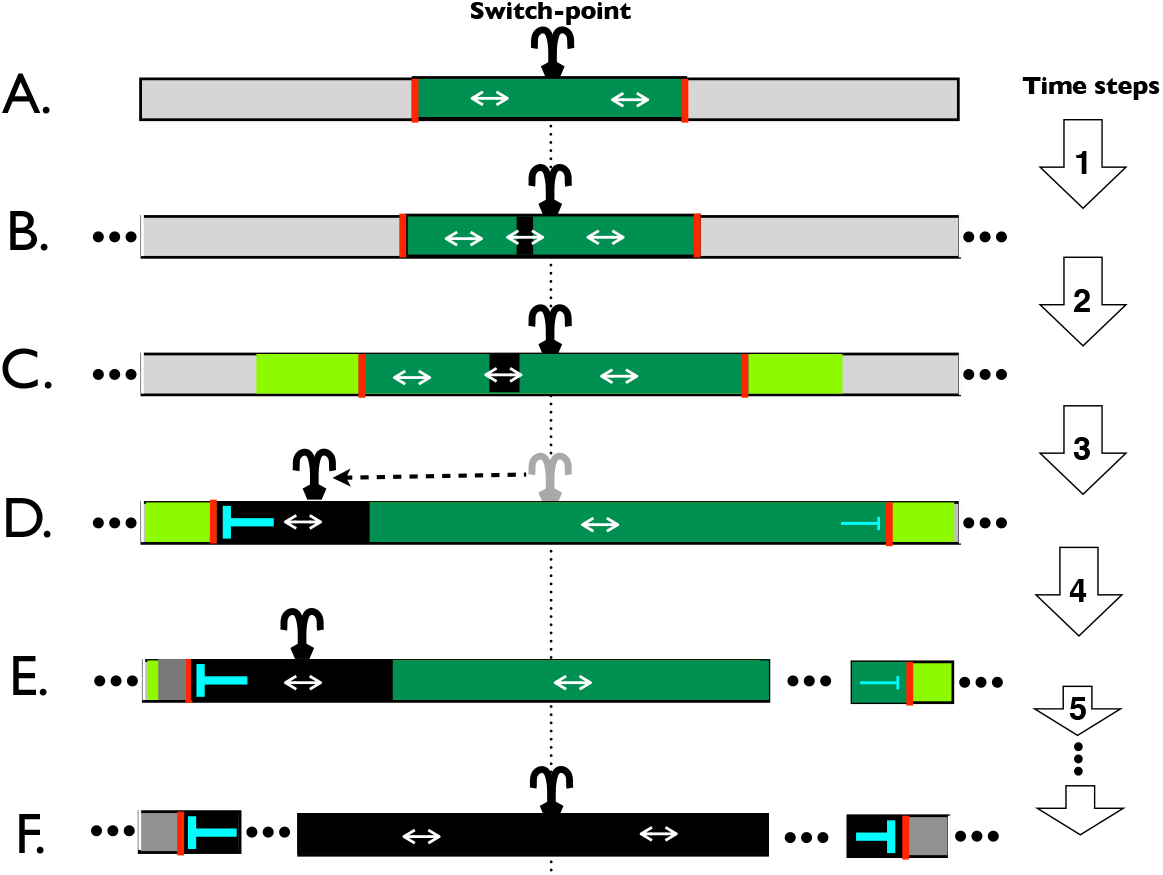
Invasion of a new HOR subarray with stronger PHI-resistance (black rectangle) into an established centromeric HOR array (combined green [centric core] and grey [pericentric flanks] rectangles). **A-B.** The subarray enters the centric core of the established HOR array at position far from either boundary (red lines) between the centric core and the pericentric flanks. **C.** The new array expands in absolute and proportionate size with time due to recurrent fork-stalling/collapse/BIR. The proportionate size increases because, unlike the surrounding (established) HOR array, none of its expansion is lost to the inward-moving pericentric boundary (light green). **D.** When the new subarray reaches the left centric/pericentric boundary, it will begin to express its stronger resistance to the recurrent invasion of pericentric heterochromatin (higher PHI-resistance) that occurs as the centric core expands. As the pericentric flanks ingress into the edges of the centric core (while the core continually expands via fork-stalling/collapse/BIR), most of this invasion is on the right side of the centric core due to its weaker PHI-resistance. When the new subarray reaches the left centric/pericentric boundary, the switch-point immediately moves toward the new array, to a position proportional to the ratio of PHI-resistance values. For example, if the new array has five times higher PHI-resistance (so that on average, for every one base pair moved into the pericentric flank on the left, five base pairs are moved into the right percentric flank), then when the new subarray reaches the left pericentric boundary, the switch-point will immediately move to a position one sixth of the way from the left boundary. **E.** If this repositioned switch-point is located within the expanded subarray when it reaches the left boundary, the new HOR will be pushed in both directions as the centric core continually expands via fork-stalling/collapse/BIR. **F.** Eventually the new HOR subarray will spread to the entire centric core, and also into the pericentric flanks. Once recurrent deletion pressure removes all of the original array from the pericentric flanks (where dense cohesin clamps associated with pericentric heterochromatin prevent lateral expansion via fork-stalling/collapse/BIR), the new HOR with faster lateral expansion rate will span the entire centromeric array.

For example, suppose the new subarray invades far from the edge of the centric core (left side in Figure 8A,B). Because the new sequence does not change the expansion rate of the left side of the array, the switch-point remains in the center of the centric core. With time, the new array is pushed outward to the left and it expands as it does so (Figure 8C). Eventually it is pushed to the left edge of the expanding centric core (Figure 8D). At this point onward, HOR elements are pushed into the pericentric flanks five times as fast on the right side compared to the left side. This asymmetry instantaneously moves the switch-point to the left, to a position one sixth of the way between the left and right sides of the centric core (because the rate of movement into the left and right pericentric flanks is 1:5). If the new array began in a position close enough to the center of the centric core, the repositioned switch-point will reside within the new subarray (deep enough for its HORs to span both of its sides) when it encounters the left pericentric flank, and it will eventually spread to all positions within the centric core –and ultimately the entire HOR array once deletions have removed all older repeat elements in the pericentric flanks. The same logic applies to the case where the switch-point is displaced from the center due to neighboring heterochromatin effects on the rate of pericentric heterochromatin spreading: if a subarray with higher PHI-resistance invades close enough to the switch-point so that it encompasses it before being completely pushed out into the pericentric heterochromatin, the new, higher PHI-resistant subarray is predicted to eventually replace the sequence of the entire centromeric array.

### Joint effects of lateral expansion rate and PHI-resistance can be counterbalancing

In the above two sections, I motivated how HOR subarrays that expand faster or have higher PHI-resistance can invade and ultimately replace an extant HOR array. In this section, I provide a rationale for how counterbalancing features of a subarray can allow those with slower lateral expansion or lower PHI-resistance to invade and replace an extant HOR array.

Consider an invading HOR subarray that has (relative to the extant active HOR array): i) somewhat slower expansion rate (e.g., half the rate), and ii) substantially higher PHI-resistance (e.g., ten times higher). The slower expansion alone would predict that the new subarray cannot replace the extant array. However this prediction can be reversed due to the much higher PHI-resistance.

To see why, consider a homogeneous centromeric HOR array, and further assume that there are no neighboring heterochromatin effects that influence the rate of pericentric heterochromatin spreading: so that before the new subarray invaded, the switch-point was located in the center of the centric core (Figure 9A [left panel]). Next suppose that a new, small subarray invades near the center of the extant array (slightly to the left of the switch-point. Figure 9B[left panel]). Despite its slower expansion rate, it initially moves very slowly toward the left edge of the centric core because it is surrounded by nearly equal blocks of the faster-expanding extant HOR array. As a consequence, by the time it reaches the left edge of the expanding centric core, it makes up a substantial part of the total centric core (Figure 9D [left panel]). At this time point, the position of the switch-point, which has been moving continuously to the right due to the slower expansion rate of the new HOR subarray (see Figure 9C [left panel]), is instantaneously repositioned to a place that is to the left side centric core (despite its slower expansion rate because it has much higher PHI-resistance (Figure 9E [left panel]). If the newly positioned switch-point is included within the new subarray (deep enough for its HOR units to span both of its sides) it will begin expanding in both directions and is now predicted to eventually spread to the entire centric core – and ultimately the entire array once recurrent deletion pressure removes all of the older HOR copies from the non-expanding pericentric flanks (Figure 9F [left panel]).

**Figure 9.**
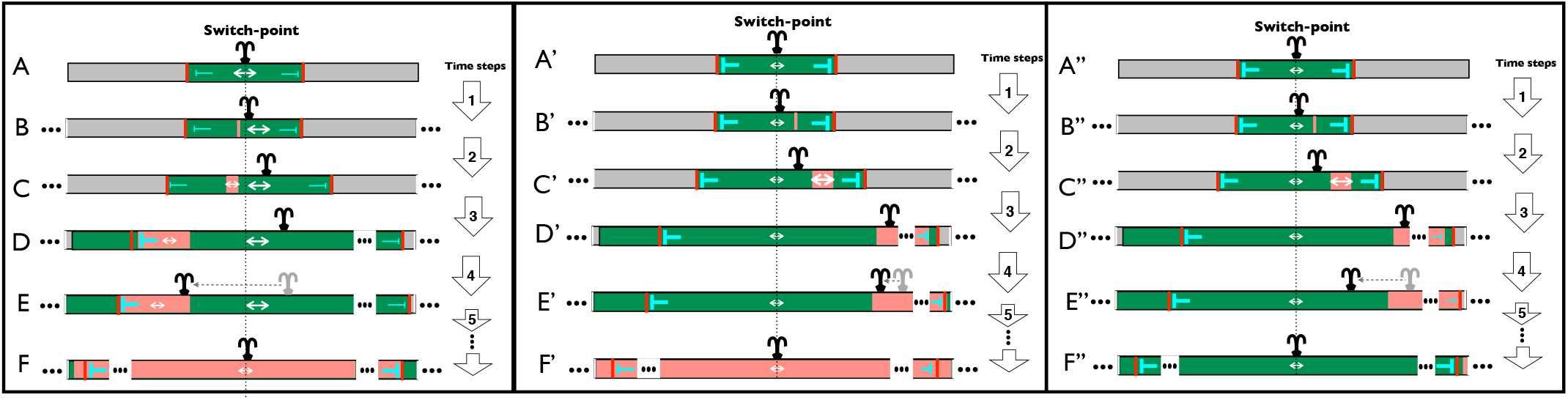
**Left:** Invasion of a new HOR subarray with slower lateral expansion rate but stronger PHI-resistance (pink rectangle) into an established centromeric HOR array (combined green [centric core] and grey [pericentric flanks] rectangles). **A-B.** The subarray enters the centric core of the extant HOR array at a position far from the centric/pericentric boundaries (red lines). **C.** The new array expands in size with time due to recurrent fork-stalling/collapse/BIR. Because it entered the established HOR array near the middle, the new subarray will expand to substantial size before reaching the left pericentric boundary, despite its lower lateral expansion rate. **D.** When the new array reaches the left pericentric boundary, it will begin to express its stronger resistance to the persistent invasion of pericentric heterochromatin, and this higher PHI-resistance moves the switch-point strongly to the left. If the position of the switch-point is included within the new, expanded subarray, then the new HOR will be spread in both directions, eventually spreading to the entire centric core, and ultimately the entire HOR array once recurrent deletion pressure removes the old HOR sequence (no longer expanding because it is in the dense-cohesin pericentric flanks). **Middle:** All steps identical to the left panel except the new subarray has opposite characteristics: faster lateral expansion but slightly weaker PHI-resistance. As the faster expanding subarray expands and moves toward the right edge of the centric core, it eventually captures the switch-point (**D**’). All else being equal this would lead to the subarray eventually expanding in both directions and eventually replacing the original HOR. But the somewhat weaker PHI-resistance potentially changes this outcome. When the new subarray encounters the right centric/pericentric boundary (**E**’), the slightly weak PHI-resistance of the subarray is expressed: causing the switch-point to instantaneously reposition in the direction toward the opposite side of the centric core (that does not contain the new subarray). But when this repositioning is sufficiently weak (because PHI-resistance is only slightly lower), it will not remove the switch-point from the subarray, so its bi-directional spreading will allow it to eventually encompass the entire centric core, and eventually the entire HOR array once recurrent deletion pressure removes the old HOR sequence (no longer expanding because it is in the pericentric flanks with dense-cohesin (**F**’). **Right:** Same as middle panel but a strong reduction in PHI-resistance prevents the new, faster-expanding subarray from displacing the extant array.

Next consider an invading HOR subarray that has (relative to the extant array): i) substantially faster expansion rate (e.g., quadruple the rate), and ii) slightly lower PHI-resistance (e.g., 10% lower). The lower PHI-resistance alone would predict that the new subarray would not replace the extant array. However this prediction can be reversed due to the much faster lateral expansion rate (Figure 9 [middle panel]). The logic here is the same as that used in the above paragraph and the left panel of Figure 9, but in this case the pattern is reversed: the new subarray has much faster lateral expansion rate but slightly low PHI-resistance. When the new subarray invades sufficiently close to the switch-point (Figure 9B’ [middle panel]), its faster lateral expansion rate moves the switch-point in its direction as it expands over time (Figures 9C’ [middle panel]) and eventually captures the switch-point before encountering the centric/pericentric boundary (Figure 9D’). When the subarray eventually does encounter the centric/pericentric boundary, the repositioning of the switch-point (due to the slightly weaker PHI-resistance of the new subarray) moves it toward the other side of the centric core (the side not containing the new subarray)(Figure 9E’). But because this shift in position of the switch-point is small, the switch-point is not moved out of the faster-expanding subarray: thereby enabling it to displace the original HOR array (Figure 9F’ [middle panel]).

Lastly consider an invading HOR subarray that has (relative to the extant array): i) somewhat faster expansion rate (e.g., double the rate), and ii) substantially lower PHI-resistance (e.g., ten times lower). The faster expansion rate alone would predict that the new subarray would replace the extant array if it invaded close to the switch-point. However this prediction can be reversed due to the much lower PHI-resistance (Figure 9 [right panel]). The logic here is the same as that used in the above paragraph and the middle panel of Figure 9: but in this case the pattern is quantitatively different: the new subarray has faster lateral expansion rate but much low PHI-resistance. When the new subarray invades sufficiently close to the switch-point (Figure 9B’’ [right panel]), its faster lateral expansion rate moves the switch-point in its direction as it expands over time (Figures 9C’’ [right panel]) and eventually captures the switch-point before encountering the centric/ pericentric boundary (Figure 9D’’ [right panel]).

When the subarray eventually does encounter the centric/pericentric boundary, the repositioning of the switch-point (due to the much weaker PHI-resistance of the new subarray) moves it toward the other side of the centric core (the side not containing the new subarray)(Figure 9E’’ [left panel]). If this shift in position of the switch-point is sufficiently large, the switch-point will be moved out of the faster-expanding subarray: thereby preventing it from displacing the original HOR array (Figure 9F’’ [right panel]).

## Base substitution rate and maximum size of HOR arrays

### Orthologous HORs in closely related species predicted to have elevated base substitution rates

The predicted bidirectional flow of HORs within an array (from the switch-point toward the pericentric flanks) is fueled by recurrent expansion of the centric core via fork-stalling/collaps/BIR. The elevated use of BIR within the centric core is expected to substantially increase the base substitution rate within HOR elements. This increase is expected because extensive studies in yeast (reviewed in Sakofsky et al. 2012), and more limited work in humans (e.g., Costantino, et al. 2014) indicate that the replication forks generated by fork-stalling/collapse/BIR use error-prone DNA polymerases that increase the base substitution mutation rate over 1,000-fold. This error-prone replication is expected to generate sequence variation among HOR units within the same active centromeric array. Over time, however, the only mutations that accumulate and generate between-species divergence among orthologous HOR arrays (in closely related species) are those that originate near the switch-point (and have copies that eventually span it) and subsequently spread to the entire HOR array. For these reasons, orthologous HORs in closely related species are predicted to exhibit elevated rates of base substitution (compared to the rest of the genome) because of elevated use of BIR during DNA replication (Supplemental Figure S3).

Nearly all centromeric HOR arrays in chimps and humans are exceptionally highly diverged in sequence (Archidiacono et al. 1995): but as described below, this extreme divergence probably reflects HOR replacement rather than high substitution rate within the same HOR clade in sister species. One exception is the X chromosome. The consensus sequence for both humans and chimps is 12 monomers long. Monomers with the highest sequence similarity have co-linear arrangements within their HORs, and their sequence divergence averages 6.2% (Supplemental Figure S6). The similar ordering of closely related monomers in both species indicates that they are orthologs. Their level of sequence divergence, however, is more than five times greater than the average sequence divergence seen at both single copy and repeated sequences elsewhere in their genomes (1.2%; Brittan 2002). The elevation in divergence rate at the X-linked centromeric HORs occurs despite the X chromosomes spending two-thirds of their time in females –which have an order of magnitude fewer germ-line mitoses per generation (Wilson Sayers and Markova 2011). The observed high sequence divergence of the orthologous HORs is consistent with: i) elevated mutation rates due to increased levels of DNA replication via BIR replication forks that have markedly higher substitution rates, and ii) recurrent homogenization of each species’ HOR array due to the continuous flow of HOR units from the switch-point to the rest of the array.

### Continually expanding centromeric HOR arrays are predicted to be limited in size by recurrent large deletions

Unlike rDNA repeat arrays, that have a mechanism to terminate tandem expansion once a sufficient array size is achieved (the E-pro bidirectional promoter; Kobayashi 2014; Box 1), no evidence for such a mechanism is evident in the structure of human centromeric HOR arrays. Lack of such a braking mechanism would be expected to lead to exceptionally long HOR arrays at centromeres: and correspondingly, exceptionally long centromeric HOR arrays (>8Mb) have been observed (Miga et al. 2014). But as described in the next paragraph, rare, recurrent, large-scale deletions may put a cap on the expansion of centromeric HOR arrays and limit the maximum size they actually achieve in nature.

Consider an HOR array that is increasing in length (more tandem HOR units) on an arbitrary chromosome over time because expansion via fork-stalling/collapse/BIR exceeds contraction via SSA repair of DSBs. At a single point in time, different chromosomes would have different histories of stochastic fork-stalling/collapse/BIR (and SSA deletions), and these differences would generate a distribution of array sizes. However, rare large deletions would also recurrently occur (e.g., Lo et al. 1999), many of which are expected to be neutral because, although smaller, these HOR arrays with deletions are nonetheless sufficiently large to nucleate a kinetochore with normal functioning (i.e., at least 50-100 Kb; Lo et al. 1999; Yang et al. 2000; Okamoto et al. 2007). In this case, over a long period of time a succession of neutral deletions would be expected to drift to high frequency, and eventual fixation. Each deletion that fixed by drift would displace the larger HOR array that generated them (Kimura 1983) and gradually expand to longer size (more tandem HOR units) (Supplemental Figure S7). This drift-based replacement process would generate transient polymorphism (of larger and smaller average array sizes) and produce a bimodal distribution of array sizes (Supplemental Figure S7A). If the large deletion rate (Udel) were sufficiently high relative to effective population size (Ne), so that 4NeUdel ≥ 2, more than one size allele would be expected to be segregating simultaneously at most time points, and a persistent bimodal or multimodal distribution of HOR sizes would be expected (Kimura 1983; Supplemental Figure S7C). Bimodal distributions for array size have been observed in large samples of X and Y chromosomes (Miga et al. 2014), indicating that large, neutral deletions are sufficiently common to generate at least transient polymorphisms for larger and smaller size classes. This same study found the tail of the larger mode of the distribution of HOR array sizes on the X to extend to 8.3 Mb long. Because most HOR arrays are much shorter than 8.3 Mb (Willard 1991), large deletions to neutral length-alleles are feasibly sufficiently common to keep most HOR arrays far from their maximal size and in a state of perpetual expansion.

The restriction of expansion via fork-stalling/ collapse/BIR to the centric core of the active HOR array motivates a testable prediction concerning inactive HOR arrays (found on at least half of human chromosomes; UCSC genome browser [GRCh38]): inactive HOR arrays will be smaller. Unless they were recently replaced by a new active HOR array, inactive HOR arrays are expected to be much smaller than the active HOR array because they have been shrinking for a protracted time span toward eventual loss via recurrent SSA-induced deletions (and rarer NHEJ repair of pairs of DSBs) (Supplemental Figures S1-S2) while not expanding via fork-stalling/ collapse/BIR (Supplemental Figure S3). This prediction is supported by the smaller size of all inactive HOR arrays on all chromosomes that have them, compared to the active arrays (UCSC genome browser [GRCh38], see also Supplemental Table 4 of Nechemia-Arbely et al. [2017] and panel H3 in Supplemental Figure S7 in the companion paper, Rice 2019).

## Effects of HOR length

### Faster expansion rate predicted for longer HORs

Another consequence of fork-stalling/collapse/BIR at human centromeres is that –all else being equal– subarrays of longer HORs (i.e., those containing more monomers) are predicted to have a lateral expansion rate that is higher (i.e., the number of monomers within a subarray increases faster) than subarrays of the same length but with shorter HORs. As described above, the data from rDNA tandem repeats in yeast indicate that out-of-register BIR after fork collapse is more commonly downstream (array expansion) than upstream (array contraction), leading to BIR-induced array expansion when the array is small and cohesin is not too dense (Kobayashi 2014; Supplemental Figure S3). The repeat units in rDNA are highly homogeneous in length, presumably due to natural selection against indels that change their sequence and hence reduce their cellular functioning. In sharp contrast, HORs across the 24 human centromeres are exceptionally variable –ranging in length by over an order of magnitude (from 2 to 34 monomers). Does this length variation have consequences for the rate of expansion of HOR arrays?

Consider the shortest active human HOR which is a simple 1-dimer HOR (a b/n-box dimer, Figure 10A,B). After a fork-collapse, the truncated, partially replicated sister chromatid can find rad51-mediated homology (and reinitiate DNA replication via BIR) downstream at −1 dimer, −2 dimers, −3 dimers, and so on (Figure 10B). Studies of ectopic recombination in yeast indicate that proximal homology sites are found and used during homology search and DNA repair far more often than more distal sites (e.g., Wang et al. 2017). This observation indicates that most expansion events will add only one dimer to the HOR array. Next consider the second smallest active HOR in humans which is 2 dimers long. After fork-collapse, downstream homology for BIR re-initiation of DNA replication can be found at −2 dimers, −4 dimers, −6 dimers, and so on. For this HOR, most expansion events will add two dimers to the HOR array (Figure 10C). Extending this logic, the longer the HOR (Figure 10D), the faster the lateral expansion rate of the HOR array, all else being equal, i.e., an expansion benefit to longer HORs is that they grow by more monomers per out-of-register, downstream BIR event after fork collapse.

**Figure 10.**
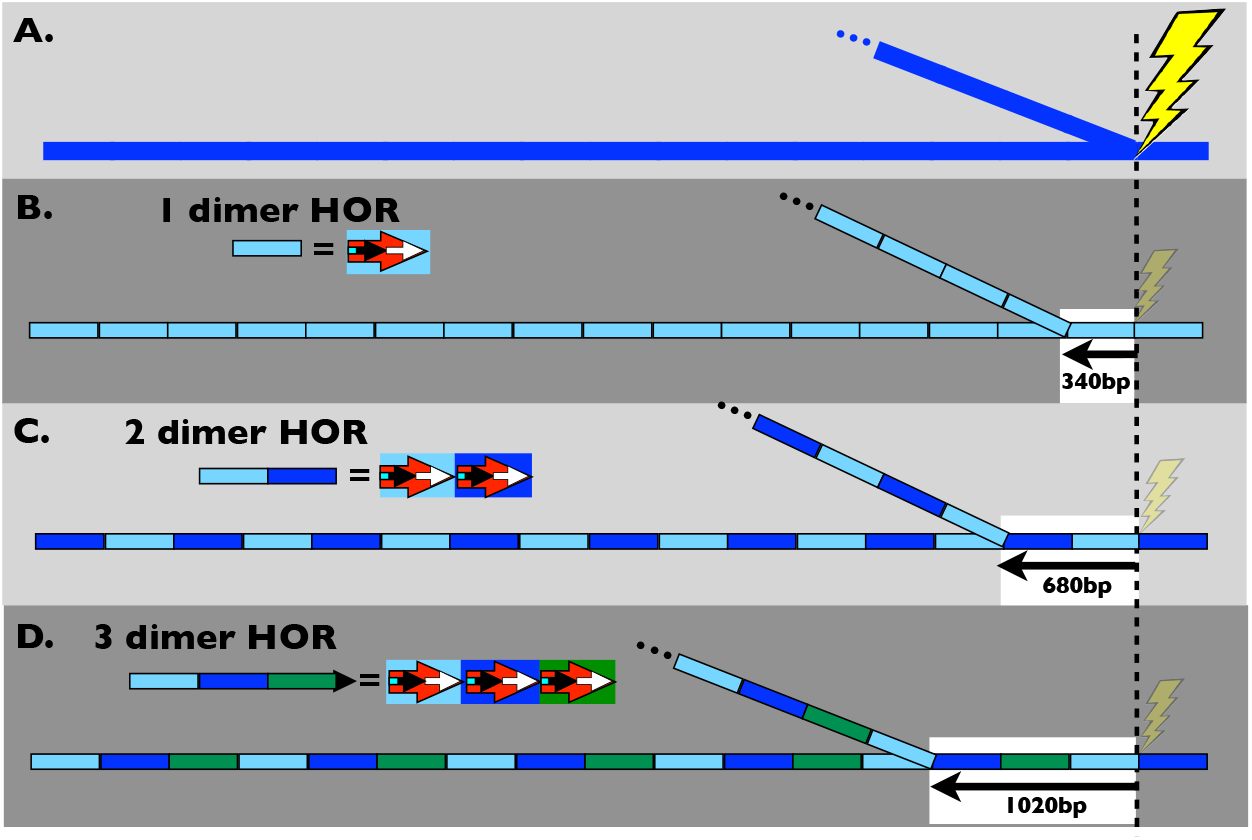
Longer HORs are expected to expand more per fork-stalling/ collapse/BIR event because the closest downstream homology to reinitiate replication is further away. Small black arrow with blue tail = b-box monomer (blue tail = b-box); small white arrow = n-box monomer; large red arrows depict dimers containing a pair of b-box and n-box monomers. In later figures, dashed large arrows denote dimers with mutated b-boxes (blue tail missing on small black arrows) that no longer bind CENP-B.

But there is also an expected expansion cost to a longer HOR: they have a greater distance separating a fork-collapse-induced, one-ended DSB (at the end of the truncated sister chromatid, Supplemental Figure S3) and its closest point of downstream homology within the HOR array on the full-length, sister chromatid (Figure 10). As pointed out above, in yeast there is substantial evidence that longer intervals separating a DSB and it closest point of homology reduces the efficacy of rad51-based homology search (e.g., Wang et al. 2017), which in the context of BIR after fork-collapse in a tandem repeat, would make in-register re-initiation of DNA replication via BIR more likely and expansion less likely. When I modeled the combined expansion costs and benefits of longer HORs, I found that longer HORs have an expansion advantage over a broad range of the parameter space (Supplemental Figure S8). I also found quantitative data on the reduction in the efficacy of homology search with distance from a DSB (Renkawitz et al. 2013) that indicates an expansion cost that is too small (i.e., efficacy of homology search declines too slowly with increasing distance between the DSB and the point of homology) to offset the expansion benefits of longer HORs (Supplemental Figure S8). On balance, the available information indicates that longer HORs will feasibly have a faster lateral expansion rate.

An additional advantage of a longer HOR is that the distance separating homologous sections of neighboring repeats is longer. Longer distances between homologous repeat units reduces the rate of deletion via SSA repair of DSBs (Schildkraut et al. 2005) and thereby increases the net expansion rate of an HOR array –as described in the following section.

### Faster shrinkage rate predicted for longer HORs, but it appears to be insufficient to counterbalance their faster expansion rate

The faster lateral expansion rate of longer HORs will be at least partially counterbalanced by a higher rate of contraction by deletion (shrinkage) during repair of DSBs via SSA (Supplemental Figure S9). Hereafter, I will use the term DSB to mean the typical form of double-stranded DNA break that generates two ends: not including the one-ended DSBs (producing a truncated sister chromatid) that are generated during replication fork collapse.

The SSA repair pathway deletes the intervening sequence between neighboring repeats (Sugawara et al. 2000; Supplemental Figure S1). In the context of HORs, SSA repair is expected to delete ‘n’ monomers per DSB repaired (= the length of the HOR = the distance between homologous monomers), where n is the number of monomers in the HOR (Supplemental Figure S9). So when an HOR is longer, more sequence is lost per DSB repaired via SSA, and longer HORs may be at a shrinkage disadvantage, i.e., they shrink faster than shorter HORs. However, longer HORs separate regions of homology (homologous monomers) by a longer distance and this greater separation reduces repair by the SSA pathway and increases repair by deletion-free pathways (Schildkraut et al. 2005), as described below. If this reduction is sufficiently large, longer HORs could have a shrinkage advantage because they utilize the SSA repair pathway sufficiently less frequently than shorter HORs, despite the fact that they shrink more per SSA repair of a DSB (Supplemental Figure S10). In the following paragraph I consider the net consequence of these two opposing effects of HOR length on the shrinkage rate due to SSA repair of DSBs.

Data on the repair of DSBs at centric and pericentric repeat arrays in mice *(Mus musculus domesticus)* indicate that there are three major pathways (Tsouroula et al. 2016). DSBs generate a pair of DNA ends that must be rejoined (repaired). Un-resected DNA ends are repaired via NHEJ, which generates little or no shrinkage via deletion. When the DNA end are resected, the SSA repair pathway competes with a group of deletion-free repair pathways leading to gene conversion and collectively called **H**omology **D**irected **R**epair (HDR, including repair by **S**ynthesis-**D**ependent **S**trand **A**nnealing [SDSA], and repair via double Holiday structures; Fishman-Lobell et al. 1992; Schildkraut et al. 2005). In Supplemental Figure S10 I quantify the shrinkage costs and benefits of longer HORs during repair of DSBs and show that the probability of SSA repair (instead of repair via HDR when DNA ends are resected) must drop precipitously with distance between repeats in order for longer HORs to have a lower rate of shrinkage. Data in humans measuring the decreased use of SSA (vs. HDR) as distance between repeats increases (Schildkraut et al. 2005), indicates that the decline is too slow to cause longer HORs to have a shrinkage advantage. However, this work was not done with human HORs. Nonetheless, most of the parameter space (Supplemental Figure S10) leads to longer HORs having a shrinkage disadvantage (i.e.,they shrink faster) during the repair of DSBs. Data from the repair of DSBs in the centric and pericentric chromatin of mice (Tsouroula et al. 2016) indicates that while SSA repair of DSBs is sometimes used, the predominant pathways for repair are NHNJ and the HDR pathways: so the shrinkage cost of longer HORs probably is small, relative to the expansion advantage via fork-stalling/collapse/ BIR, because the SSA repair pathway is sufficiently uncommon at centromeric HORs.

In sum, the available data indicate that expansion events (fork-stalling/collapse/BIR) favor longer HORs and one type of shrinkage event (SSA-repair of DSBs) disfavors longer HORs. Data from mice indicates that the strong predominance of HDR and NHEJ over SSA repair at centromeric repeats reduces substantially the shrinkage cost of SSA repair of DSBs. The fact that many HORs have evolved very long length (e.g., 34 monomers for the Y chromosome) –despite recurrent deletion pressure that would continually generate embedded, shorter-HOR subarrays– is consistent with the assumption that the expansion advantage of longer HORs prevails over their shrinkage cost and that longer HORs have a net expansion advantage. I next explore the consequences of a faster lateral expansion rate of longer HORs.

### Longer HORs predicted to out-compete shorter HORs in intra-array competition

Longer HORs with a faster lateral expansion rate are expected to have a replicative advantage (i.e., more rapidly expand the length of their tandem sub-array) when they invade an array composed of a shorter HOR. If a small subarray of a longer HOR invades an extant centromeric array (and they differ in no other factors that influence lateral expansion rate or PHI-resistance), the longer subarray is expected to eventually displace the shorter one whenever it invades close enough to the switch-point (Figure 7). It can ultimately replace the shorter HOR array because its faster lateral expansion rate will move the switch-point increasingly closer as its subarray expands while being pushed toward the pericentric flank (Figure 7B-C). If it captures the switch-point before being pushed off the side of the centric core (Figure 7C), copies of the new HOR will span both sides of the switch-point and spread it in both directions: ultimately encompassing the entire centric core. Over time, recurrent deletions are expected to remove the older (shorter) HOR from the pericentric flanks (Figure 7E).

Chromosomes 1, 5, and 19 share the same (or very similar) short HOR sequence at their active HOR arrays (a 1-dimer HOR; UCSC genome browser [GRCh38]). This short HOR may be in the process of expanding to a longer HOR. Long PaBio reads of a homozygous genome indicate that the HOR arrays of these chromosomes contain at least one long, homogeneous region of the predominant 1-dimer HOR. However, many isolated regions of longer HORs (2-, 3-, and 4-dimer repeat units that include and extend the predominant 1-dimer repeat unit) can be found (see Supplementary Figure S4 of the companion paper, Rice 2019) which are feasibly the early steps of the expansion of the predominant 1-dimer HOR to a longer HOR. Most remarkable are long, homogeneous runs of a 4-dimer expansion of the predominant 1-dimer HOR. The logic developed here predicts that, over time, these contiguous patches of the expanded HOR (4 dimers long) will displace the predominant shorter HOR (1 dimer long) and will become the sole HOR in future generations –so long as it resides within the centric core of the HOR array and did not originate too near the centric/pericentric boundary.

## Trafficking among HOR arrays

### Winning HORs in intra-array competition are predicted to move horizontally to new lineages

The lateral expansion advantage of a longer HOR subarray will only lead to the longer HOR replacing the shorter one within the chromosomal lineage that it invaded. To spread population-wide, assuming no natural selection or transmission advantage (as might occur from centromere drive; see below), the stochastic and slow process of random genetic drift would be required –unless some other mechanism spreads the longer HOR horizontally to other chromosomal lineages. The concerted evolution that has homogenized the active HORs on chromosomes 1, 5, and 19 (and also a different HOR on chromosomes 13 and 21 and another on chromosomes 14 and 22; Ziccardi et al. 2016) demonstrates that some mode of horizontal transfer must be occurring at a nontrivial rate between different chromosomes sharing the same HOR. For the case of horizontal transfer between homologs, work by Roizes 2006 and Pironon et al. 2010 provides strong empirical evidence for substantial horizontal transfer of ‘diagnostic variant nucleotides’ (DVNs) between HOR arrays found in different lineages –despite strong evidence against meiotic and/or mitotic crossovers between homologs. This noncrossover exchange among homologous HOR arrays is also supported by substantial additional evidence that is reviewed in Talbert and Henikoff (2010). Ectopic gene conversion via the SDSA pathway and/or BIR with template switching are feasible candidate mechanisms for this horizontal transfer without crossover (Mladenov et al. 2016). I will refer to this non-crossover, horizontal transfer between homologs as ‘lineage-spreading’.

A non-trivial rate of horizontal movement between homologous HOR arrays would permit a new, longer HOR expanding within one lineage to spread to other lineages, where it could expand and replace the established, shorter HOR as the active centromeric sequence. Note that a new HOR subarray expanding within one lineage shares flanking homology with all other lineages (at its edges where is abuts with the established HOR sequence) that do not contain the subarray: this flanking homology would be expected to facilitate transfer of the new HOR sequence between lineages. In sum, when coupled with horizontal transfer between chromosomal lineages, longer HORs with a faster lateral expansion rate have the potential to (eventually) globally replace shorter HORs across all lineages of a chromosome within a population.

### A simple BIR pathway is predicted to generate longer HORs

Another consequence of fork-stalling/collapse/BIR at human centromeres is that it generates a simple pathway for the enlargement of shorter HORs to longer HORs, i.e. those with more monomers (Figure 11). For example, consider the expansion of a simple 1-dimer HOR (a b/n-box dimer) at a centromeric repeat array to a 2-dimer HOR array. The rationale for short HORs expanding in units of dimers –rather than single monomers– is described in a later section. The first step in producing a longer, 2-dimer HOR is the recruitment of a new repeat element (i.e., a new b/n-box dimer shown in blue in Figure 11) into the active 1-dimer HOR array. Ectopic gene conversion (Benovoy and Drouin 2009; Hastings 2010) and/or mmBIR with template switching (Smith et al. 2007; Zhang et al. 2009; Hastings et al. 2009A,B; note: mmBIR = microhomology-mediated BIR) provide a plausible mechanism for this recruitment. Both of these processes are facilitated by sequence similarity between the donor and recipient HORs (Hastings 2010; Anand et al. 2014), and dimers within the same HOR nearly always share sequence similarity, i.e., they belong to the same cluster (of the four major clusters of dimers; see Figure 5 in Rice 2019). Fork-collapse during DNA replication that occurs one dimer upstream from the newly recruited dimer can reinitiate replication via BIR immediately downstream of the new dimer, and this event leads to a new, longer HOR (a 2-dimer HOR subarray in Figure 11). Because the new 2-dimer HOR is longer, it will feasibly have a lateral expansion advantage (Figure 10), enabling it to gradually displace the entire 1-dimer HOR array over time (Figure 7) within the lineage where it originated. The new dimer could eventually spread population-wide in all lineages due to exchange between homologs via ectopic gene conversion and/or BIR with template switching, i.e via lineage-spreading.

**Figure 11.**
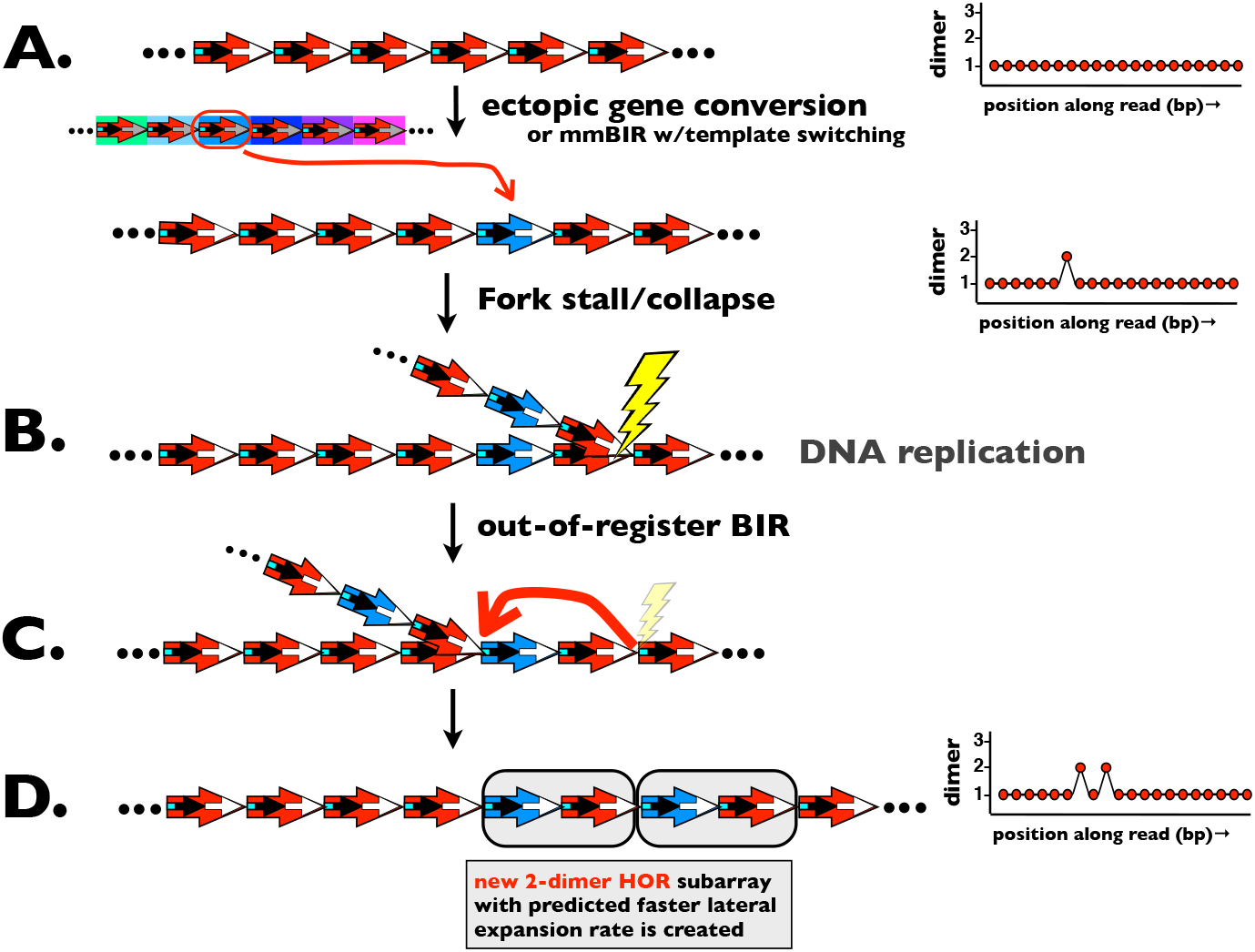
BIR and mmBIR facilitate the creation and expansion of a new, longer HOR with a faster lateral expansion rate. **A**. New dimer from a different HOR array (large blue arrow) enters a 1-dimer HOR array (large red arrows) via ectopic gene conversion or mmBIR with template switching. **B**. Fork-collapse during DNA replication. **C**. Downstream BIR re-initiation of DNA replication. D. A new 2-dimer HOR is created which is predicted to have a faster lateral expansion rate and can thereby eventually predominate within the initially 1-dimer HOR array. Arrow key as in Figure 10.

In the same manner, a new dimer could enter the 2-dimer HOR and give rise to a 3-dimer HOR with a lateral expansion advantage that eventually replaces (population-wide) the 2-dimer HOR array. This process could be reiterated to eventually make long HORs like the one at chromosome 20, which is an 8-dimer HOR. Because ectopic gene conversion and mmBIR with template switching in humans usually insert < 1 kb of new DNA into the recipient strand (Chen et al. 2007; Padhukasahasram and Rannala 2013) and an average of ~370 bp in the case of gene conversion (Benovoy and Drouin 2009), HORs would be expected to expand in length gradually in steps of one dimer at a time (or no more than a few monomers). Evidence for HOR expansion in length can be found within centromeric HOR arrays currently active in the human genome, e.g., interspersed two-, three-, and four-dimer HOR subarrays can be found within the predominantly one-dimer centromeric repeat array that is shared between chromosomes 1, 5 and 19 (see Supplementary Figure S4 in the companion paper, Rice 2019).

### A simple pathway is predicted for short HORs to invade long HORs

About half of the 23 humans chromosomes have more than one centromeric HOR array (UCSC genome browser [GRCh38]; Ziccardi et al. 2016), although current evidence indicates that only one HOR array is active on a single chromosome (e.g., Aldrup-MacDonaldet al. 2016). How does a new HOR array become established on a chromosome? Although there are many possibilities, here I illustrate a simple pathway by which BIR can facilitate this process. I will illustrate this pathway in the context of one-dimer HOR invading (and then expanding within) a long HOR with highly reduced b/n-box dimer modular structure (Figures 12 and 13). This particular case will be important later on when I consider the influence on b/n-box dimer structure on the lifecycle of HORs. The first step is the insertion of a copy of a b/n-box dimer from another HOR into the longer HOR array (with highly reduced b/n-box dimeric modular structure) via ectopic gene conversion or mmBIR with template switching (Figure 7A). There is evidence for dimer insertions that originated at a different chromosome at the HOR shared by chromosomes 1, 5, and 19 (see Supplemental Figure S4 in the companion paper, Rice 2019). The high mutability of mmBIR replication (Deem et al. 2011; Sakofsky et al. 2012), and the fact that only a part of a dimer may be replaced in the new location, will cause the newly formed b/n-box dimer to usually be similar but unique in sequence –rather than an exact copy of a dimer from another HOR. Fork-stalling/collapse followed by downstream, out-of-register, mmBIR next duplicates the dimer and produces a direct repeat of the dimer that is embedded within the longer HOR (Figure 12B-E). Repair of collapsed replication forks via mmBIR is thought to play a major role in generating copy number variation (CNV) in humans (Hastings et al. 2009B; Hsiao et al. 2015). Finally, repeated out of register BIR events can expand the new, tandem repeat (a new one-dimer HOR sub-array that is embedded within the old, longer HOR array) (Figure 13).

**Figure 12.**
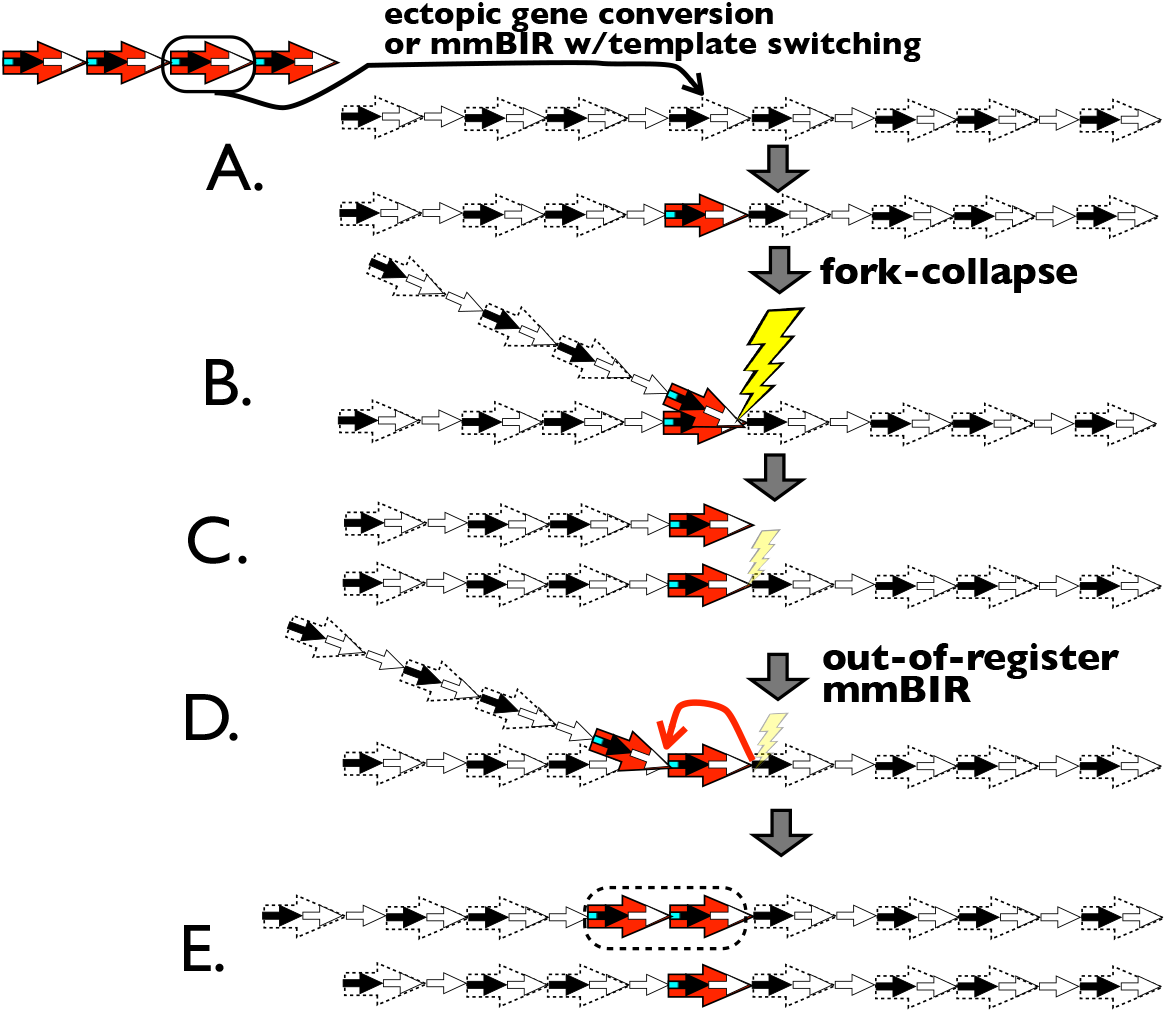
Fork-stalling/collapse/mmBIR facilitates the recruitment of short HORs with modular b/n-dimer structure into a longer HOR array that has lost this modular structure (b/n-dimer-decayed array). **A.** A dimer from a different HOR (a 4-dimer HOR in this example) is recruited to the b/n-dimer-decayed array. **B-C.** Fork-stalling followed by fork-collapse generates a one-ended DSB that represents an incompletely replicated sister chromatid. **D.** DNA replication is reinitiated one dimer downstream via mmBIR. The imported dimer has now formed a tandem duplication (a b/n-dimer subarray embedded within the b/n-dimer-decayed array). Arrow key as in Figure 10.

**Figure 13.**
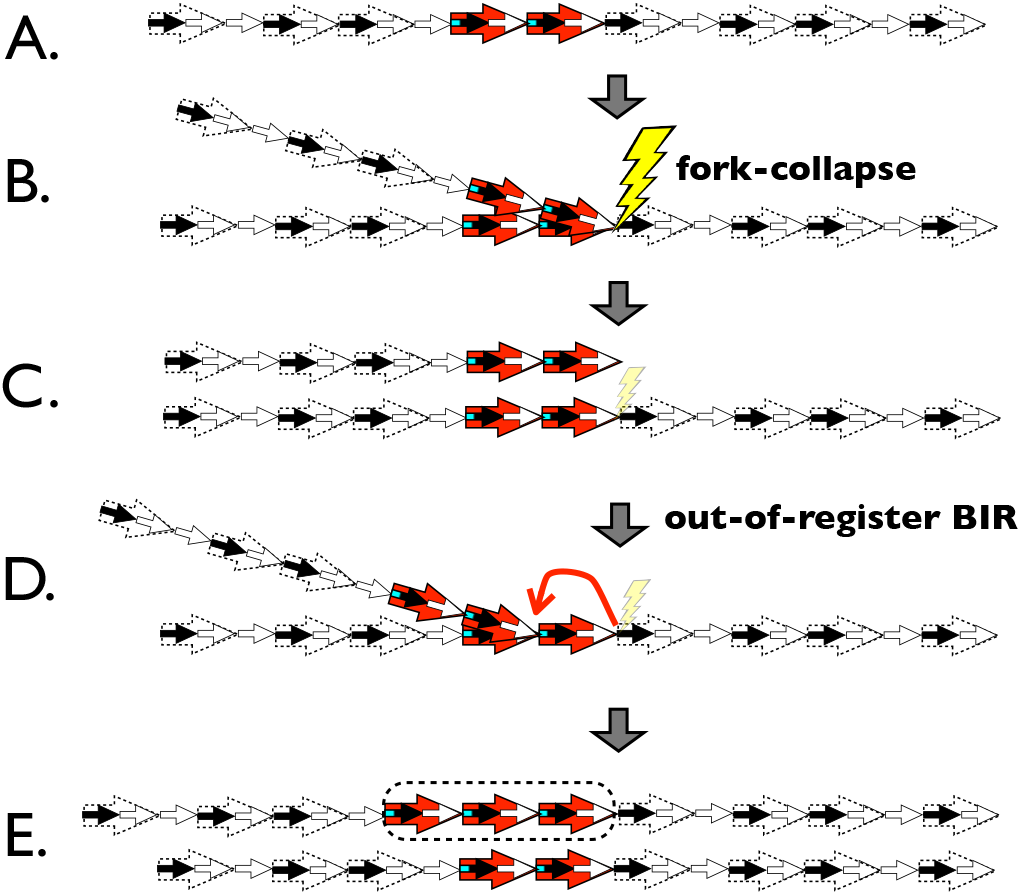
A short 1-dimer HOR subarray that is embedded in an HOR array with reduced b/n-dimer modular structure (b/n-dimer-decayed array) can be expanded via fork-stalling/ collapse/BIR. **A-C.** Fork-collapse during the replication of the dimer subarray generates a one-ended DSB forming a partially replicated (truncated) sister chromatid. **D.** Downstream BIR reinitiates replication one repeat unit downstream (closest downstream homology) via BIR. **E.** Replication of the truncated sister chromatid expands the 1-dimer subarray from two to three tandem repeats. Reiteration of this process can further expand the copy number of the 1-dimer subarray. Arrow key as in Figure 10.

## Selection in the pericentric flanks

### Selection against HOR structure is predicted in the pericentric heterochromatin

Up to this point I have focused on the consequences of fork-stalling/collapse/ BIR within the centric core region of the active HOR array: a sector that does not densely bind cohesin. In this section I focus on the pericentric heterochromatin that does have densely bound cohesin. Selection on sequence composition in this region is expected to be quite different. Expansion via fork-stalling/collapse/BIR is expected to be absent (because densely bound cohesin is expected to prevent out-of-register BIR after fork collapse [Kobayashi 2014]): so the intra-array selection for faster lateral expansion rate and higher PHI-resistance found in the centric core also will be absent. In addition, the pericentric heterochromatin has no known influence on centromere strength (i.e., the level of assembled kinetochore proteins) –and hence, no selection based on centromere drive (Wu et al. 2018). But there will be selection to avoid deletion during SSA-repair of DSBs. Such deletion avoidance would occur by promoting **H**omology **D**irected **R**epair (HDR; via **G**ene **C**onversion [GC], including the SDSA pathway and repair via double Holiday structures; Fishman-Lobell et al. 1992; Schildkraut et al. 2005) over SSA-repair. The primary feature favoring HDR over SSA is a longer distance between repeats (Fishman-Lobell et al. 1992; Schildkraut et al. 2005; Kappeler et al. 2008). This relationship would favor mutations that reduce homology between neighboring repeat elements in an HOR array, such as point mutations, short inversions, and short indels. So in sum, in the pericentric heterochromatin there is selection for sequence divergence because only repeat elements that have diverged from their neighbors avoid deletion via SSA repair of DSBs and persist through time.

Shepelev et al. 2009 provide evidence that repeat elements that are more distal to the active HOR array are generally older. This arrangement predicts that repeat elements that are far from the active HOR array should have lost their HOR structure due to the long time period that was available for the accumulation of selectively favored point mutations, small inversions and indels. This structure is observed in the distal pericentric regions surrounding active HOR arrays (Ross et al. 2005; Nusbaum et al. 2006; Shepelev et al. 2009; Ziccardi et al. 2016): the distal flanks of the pericentric heterochromatin are made up of unorganized monomers that lack HOR structure and contain frequent small inversions and indels.

Influence of modular b/n-box dimer structure HORs with modular b/n-box dimer structure are predicted to recruit maximal CENP-C

The pattern of monomers observed at the full set of human HORs (across all chromosomes) indicates that their composition has structural constraints (Rice 2019). The strongly predominant unit found in active human HORs is the b/n-box dimer, and shorter HORs (those with 10 or fewer monomers) are made up almost exclusively of these dimers (see Figure 7 of the companion paper, Rice 2019). In sharp contrast, flanking, inactive HOR arrays –when present on a chromosome– typically have substantially reduced b/n-box dimer structure (see Figure 9 of the companion paper, Rice 2019). Why are b/n-box dimers so predominant at active HOR arrays?

One possibility is that the b/n-box dimer configuration is somehow optimal for kinetochore functioning. CENP-B binds DNA at a 17 bp b-box sequence that is located in the linker DNA between nucleosomes (Fujita et al. 2015; Hasson et al. 2013; Henikoff et al. 2015). The b/n-box dimer configuration enables a single CENP-B dimer to bind simultaneously to both of the b-boxes in neighboring dimers within an HOR (Yoda et al 1998; Tawaramoto et al. 2003). Work by Fachinetti et al. (2015) found that centromeric HOR arrays that lack b-boxes (and do not bind CENP-B) produce suboptimal kinetochores. These kinetochores contained much less CENP-C: one of the foundational pillars located at the base of kinetochores. This study found that recruitment of maximal CENP-C per nucleosome (2 molecules) requires CENP-B to bind linker DNA near CENP-A nucleosomes: otherwise nucleosomes recruit only one CENP-C molecule (Supplemental Figure S11-12), leading to weak kinetochores with elevated mis-segregation rates.

Evidence for an an additional advantage of b-boxes comes from the work of Basu et al. 2005. They found that placing b-boxes at every monomer in an HOR maximized its ability to recruit CENP-A during the establishment of **H**uman **A**rtificial **C**hromosomes (HACs). These observations lead to the intuitive notion that b-boxes would ideally be found at all the monomers in an HOR, thereby guaranteeing high kinetochore strength (and a high ability to recruit the centromere-defining epigenetic mark –CENP-A): but this is not the case.

Despite the importance of b-boxes in recruiting maximal density of CENP-C per CENP-A nucleosome, long runs of b-box monomers are absent in both the active and inactive HORs (see Figures 4 and 9 in the companion paper, Rice 2019), with the longest being a 3 monomer long run (only one occurrence that is located within the active HOR on chromosome 17). In sharp contrast, long runs of n-box monomers are found in the HOR of the Y chromosome (34 monomers) and on many of the inactive HOR arrays on the acrocentric chromosomes (e.g., one on chromosome 15 [19 monomers] and another shared by some or all of chromosomes 13, 14, 21 and 22: [16 monomers]; see Figures 4 and 9 in the companion paper, Rice 2019). At all of these HORs, the 19 bp n-box has high levels of conservation (e.g., see supplemental Supplemental Figure S11 in Rice 2019). Among the inactive flanking HOR arrays, long runs of no-b-box monomers (n-box monomers plus b-box monomers with mutated b-boxes that do not bind CENP-B) are common (see Figure 9 in the companion paper, Rice 2019). The avoidance of long runs of b-box monomers (but not monomers lacking functional b-boxes) suggests that neighboring b-box monomers are somehow suboptimal.

Empirical evidence for a disadvantage to runs of b-box monomers comes from two sources. First, Hasson et al. 2013 found that a pair of neighboring b-box monomers (a two-monomer run) on the X chromosome had markedly lower canonical nucleosome positioning and reduced recruitment of CENP-A (see their Figure 6). This pattern was also observed at an unusually long n-box monomer (containing a 14 bp insertion that produced a monomer that was 185 bp long), and also at a monomer sandwiched between the neighboring b-box monomers and the longer monomer. Second, Aldrup-MacDonald et al. 2016 identified a variant HOR of the predominant, active HOR on chromosome 17. This variant form is composed primarily of a 13mer that was generated by a 3 monomer deletion (Supplemental Figure S11C; the first three monomers are missing from the typical 16 monomer HOR on chromosome 17, as depicted in Figure 4 of the companion paper, Rice 2019). The variant HOR contains no canonical b/n-box dimers and all of its b-box monomers are in runs (two 2-monomer runs and one 3-monomer run). This 13 monomer variant HOR recruited ~40% less CENP-A and ~60% less CENP-C compared to the average from all other chromosomes. It also had substantially elevated mis-segregation rates. Yet this HOR has a higher density of b-box monomers than any other active HOR in the human genome (see Figure 7 of Rice 2019). These observations indicate that the spacing of b-box monomers within an HOR, and not just their density or presence/absence, is important in kinetochore assembly and function.

One plausible hypothesis for the functional significance of alternating b-box and no-b-box monomers (in b/n-box dimers) is that this structure places both a b-box and an no-b-box sequence adjacent to every nucleosome (Supplemental Figure S12; Yoda et al. 1998; Tawaramoto et al. 2003). The placement of DNA-bound CENP-B adjacent to every nucleosome would permit all CENP-A nucleosomes to recruit maximal levels of CENP-C (and hence make strong kinetochores; Fachinetti et al. (2015). This arrangement would also simultaneously place the no-b-box sequence (including the n-box) adjacent to every nucleosome. The no-b-box sequence is present in the linkers between all nucleosomes at the centromeric HOR of the Y chromosome (Hasson et al. 2013; Henikoff et al. 2015) and it was feasibly present in all monomers before CENP-B invaded the great ape lineage (Shepelev et al. 2015). If i) the no-b-box linker sequence historically had functional significance (see next paragraph) before the b-boxes evolved in the HORs of the great apes, and ii) the presence of CENP-B (a large 160 kD dimer) bound to the b-box interfered with this functioning, then the b/n-box dimer configuration would have preserved the ancestral sequence adjacent to every nucleosome while simultaneously permitting CENP-B to bind adjacent linker DNA and recruit the maximal level of the foundational kinetochore protein CENP-C.

### High density of b/n-box dimers within HORs is predicted to increase PHI-resistance and to be favored in intra-array competition

As described earlier (summarized in Figure 3), one feature expected to promote resistance to invasion by pericentric heterochromatin as the centric core expands via fork-stalling/collapse/BIR (i.e., to promote PHI-resistance) is increased recruitment of CENP-A. CENP-A is recruited to centromeric nucleosomes by its chaperone HJURP (Foltz et al. 2009), which is recruited by the Mis18 complex (Nardi et al. 2016), which is recruited by CENP-C (Moree et al. 2011). For this reason, HORs with a higher density of b/n-box dimers (which are predicted to recruit maximal levels of CENP-C) are predicted to have high PHI-resistance. High PHI-resistance is expected to increase the probability that a new, invading HOR subarray can replace the extant array (Figure 8). This trait is expected to also increase the probability that an extant array can resist replacement by a new, invading subarray (Figure 8).

### CENP-B is predicted to reduce fork-stalling/ collapse and to thereby reduce the lateral expansion rate of an HOR

The hypothesis that longer HORs replace shorter HORs because they have a lateral expansion advantage(Figure 10) cannot explain the observation that short HORs are usually composed entirely of canonical b/n-box dimers whereas longer HORs commonly deviate substantially from this modular structure (mainly due to mutated b-boxes that cannot bind CENP-B; see Figure 7 in Rice 2019). Is there any effect of b-box monomers on lateral expansion rate?

In fission yeast, CENP-B is coded by three loci and their gene products localize to both centromeric DNA and LTR retrotransposons (Nakagawa et al. 2002). The LTR transposons recruit the DNA-binding protein Sap1, and this bound protein causes fork-stalling, a phenotype required for retrotransposon replication and retrotransposon stability during DNA replication. CENP-B opposes this effect by counteracting Sap1 barrier activity and promoting replication fork progression through the LTR retrotransposon. CENP-B is present in most mammal species (Yoda et al. 1992; Casola et al. 2008) despite the fact that its binding site is absent at the centromeres of most of these species (Haaf et al. 1995; Alkan et al. 2011; Kugou et al. 2016). This pattern indicates that CENP-B has cellular functions other than centromere binding in mammals –and indirect evidence indicates that CENP-B mediates surveillance for retrotransposons in mammals (Kipling and Warburton 1997; Casola et al. 2008). On the assumption that CENP-B in humans reduces fork-stalling/collapse at centromeric repeats (as it does in yeast), its presence would diminish the lateral expansion rate of an HOR because of less fork-stalling/collapse/BIR. A reduced density of bound CENP-B (by point mutations within b-boxes or by dilution via an increased proportion of n-box monomers within an HOR) would be expected to increase lateral expansion rate of the HOR. So the loss of functional b-boxes (and consequent loss of modular b/n-box dimeric structure) in HORs may be explained by intra-array selection for a faster lateral expansion rate.

But then why are HORs with low densities of b-boxes only observed in long HORs (see Figure 7 in Rice 2019)? In a later section (immediately after the next section) I explore the influence of CENP-B on the boundary between pericentric heterochromatin and the centric core of an HOR array as a possible explanation. In the next section I explore the possible contribution of centromere drive (when one of the two homologous centromeres of a diploid oocyte has a higher probability of being transmitted to the ovum instead of a polar body).

### HORs with modular b-box dimeric structure are predicted to be favored by centromere drive

Centromere drive is generally assumed to be a recurrent but rare and maladaptive phenotype (at the organismal level, because driving centromeres can cause male sterility) that is usually suppressed (Malik and Henikoff 2002). However, for hundreds of millions of years eukaryotic oocytes have begun meiosis with two homologous centromeres but needed only one of these by the end of the process (assuming anisogamy). So there has been long and persistent selection to choose the better centromere when they differ in functional integrity. Although there will usually be no functional difference between the two homologous centromeres in an oocyte, centromeres can be compromised by structural damage (e.g., large deletions, epigenetic programming errors, oxidative damage, mislocalization at any of the ~100 proteins that make up the kinetochore, and so on): so there would be selection across hundreds of millions of generations of eukaryotic evolution to retain the best centromere in the ovum and divert any damaged centromeres to the polar bodies.

Although the number of studies and species is small, data from monkey flowers (Fishman & Willis 2005) and house mice (Chmatal et al. 2014; Iwata-Otsubo et al. 2017; Wu et al. 2018) indicate that centromere drive favors ‘stronger’ centromeres that recruit more kinetochore proteins. As described above, CENP-B bound to b-boxes located in the linker DNA near CENP-A nucleosomes causes a nucleosome to recruit twice the amount of the foundational CCAN protein CENP-C (Supplemental Figures S11-12; Fachinetti et al. 2015). But as also described above, the spacing of b-box monomers, rather than their density alone, i) strongly influences the recruitment of CENP-C: a high density of runs of two or more b-box monomers is associated with low CENP-C recruitment and weak kinetochores (Supplemental Figure S11C), and ii) feasibly influences the recruitment of additional features required for kinetochore assembly, such as transcription factors and other binding proteins such as pJα (Gaff et al. 1994; Romanova et al. 1996). In sum, modular b/n-box dimer structure at human HOR arrays should feasibly produce the ‘strong’ kinetochores that are expected to be favored by centromere drive.

### Opposing forms of selection are predicted on modular b/n-box dimer structure

Lateral expansion rate and centromere drive are expected to have opposing influences on centromere evolution. Within an HOR lineage, features expected to increase lateral expansion rate are: longer HORs and those with reduced modular b/n-box structure (mutated b-boxes that do not bind CENP-B and/or a higher proportion of n-box monomers). But, in competition between HOR lineages, reduced modular b/n-box dimer structure is predicted to make weaker kinetochores that are disfavored by centromere drive. So centromere drive is expected to oppose one feature that causes faster lateral expansion rate (less CENP-B binding due to reduced b/n-box dimeric structure) but not the other feature (longer HORs).

Lateral expansion rate and PEV-resistance are also expected to have opposing influences on centromere evolution in the context of reduced modular b/n-box dimeric structure. As described previously, loss of b/n-box dimeric structure is expected to increase lateral expansion rate (because of less recruitment of CENP-B) but this same feature is expected to reduce PHI-resistance (due to reduced recruitment of CENP-C, which would feasibly reduce recruitment of CENP-A). So reduced PHI-resistance is expected to oppose loss of b/n-box dimeric structure and the increased lateral expansion rate that this structure generates.

### Only long HORs are predicted to lose modular b/n-box dimeric structure

Consider the case where a mutation at the b-box of a single monomer in a short HOR (e.g., a 1-dimer HOR) prevents CENP-B from binding. As described earlier, a reduced CENP-B density is predicted to increase lateral expansion rate and potentially cause the mutant HOR to replace the original HOR (Figure 7). The reduced CENP-B binding, however, also would be expected to markedly reduce the amount of CENP-C recruited to the kinetochore (by 50%, thereby leading to a weaker centromere and reduced PHI-resistance; Supplemental Figures S11,S13), and this phenotype would feasibly lead to a strong disadvantage in centromere drive and/or the ability to capture the switch-point within a centromeric HOR array (Figure 9 [right panel]). For these reasons, mutations in short HORs that inactivate b-boxes will feasibly be prevented from accumulating because they are selected against in multiple ways.

Once the HOR becomes sufficiently long, however, mutational loss of a functioning b-box at a single b-box monomer position within the HOR would have only a small effect on CENP-C recruitment (Supplemental Figure S13). As a consequence, the reduction in kinetochore strength may be too small to substantively influence centromere drive and counterbalance the within-lineage lateral expansion advantage (Supplemental Figure S13). The same logic applies to the costs and benefits in the context of PHI-resistance (Figures 8,9).

Lineage-spreading would be expected to spread the loss of a single b-box mutation (in a long HOR) across lineages, where it could spread within these array lineages in the same way as in the above paragraph. Sufficiently long HORs can therefore begin to gradually accumulate loss-of-function mutations for b-boxes at one monomer position at a time. In this way the modular b/ n-box dimer structure of an HOR can be gradually eroded over time (nickel-and-dimed).

The same logic concerning the gradual loss of b-boxes in longer HORs applies to the recruitment of single n-box monomers that dilute the density of modular b/n-box dimers and thereby reduce centromere strength and PHI-resistance (Supplemental Figure S14). Lastly, b-box mutations that prevent the binding of CENP-B may be unusually common due to the presence of two hypermutable methylated CpG sites within the critical 9 bp of the 17bp sequence (Gopalakrishnan et al. 2009).

## The centromere HOR lifecycle

### Centromeres predicted to cycle between short, medium and long HORs

From the information surveyed above, there is substantial evidence that longer HORs (Figure 10), and those that recruit less CENP-B (Supplemental Figures S11A-B,C), feasibly have a lateral expansion advantage within a centromeric HOR array (Figure 7). This advantage is depicted by the right histogram in Figure 14. There is also evidence that HORs with a substantial reduction in modular b/n-box dimeric structure recruit less CENP-C (Supplemental Figure S11C). Recruitment of less CENP-C is expected to produce weaker kinetochores and lower PHI-resistance: depicted by the left histogram in Figure 14. These lower-CENP-C HORs would be susceptible to replacement by new, short HORs that produce stronger kinetochores and higher PHI-resistance. Collectively these patterns suggest the HOR cycle depicted in Figure 14 and described below.

**Figure 14.**
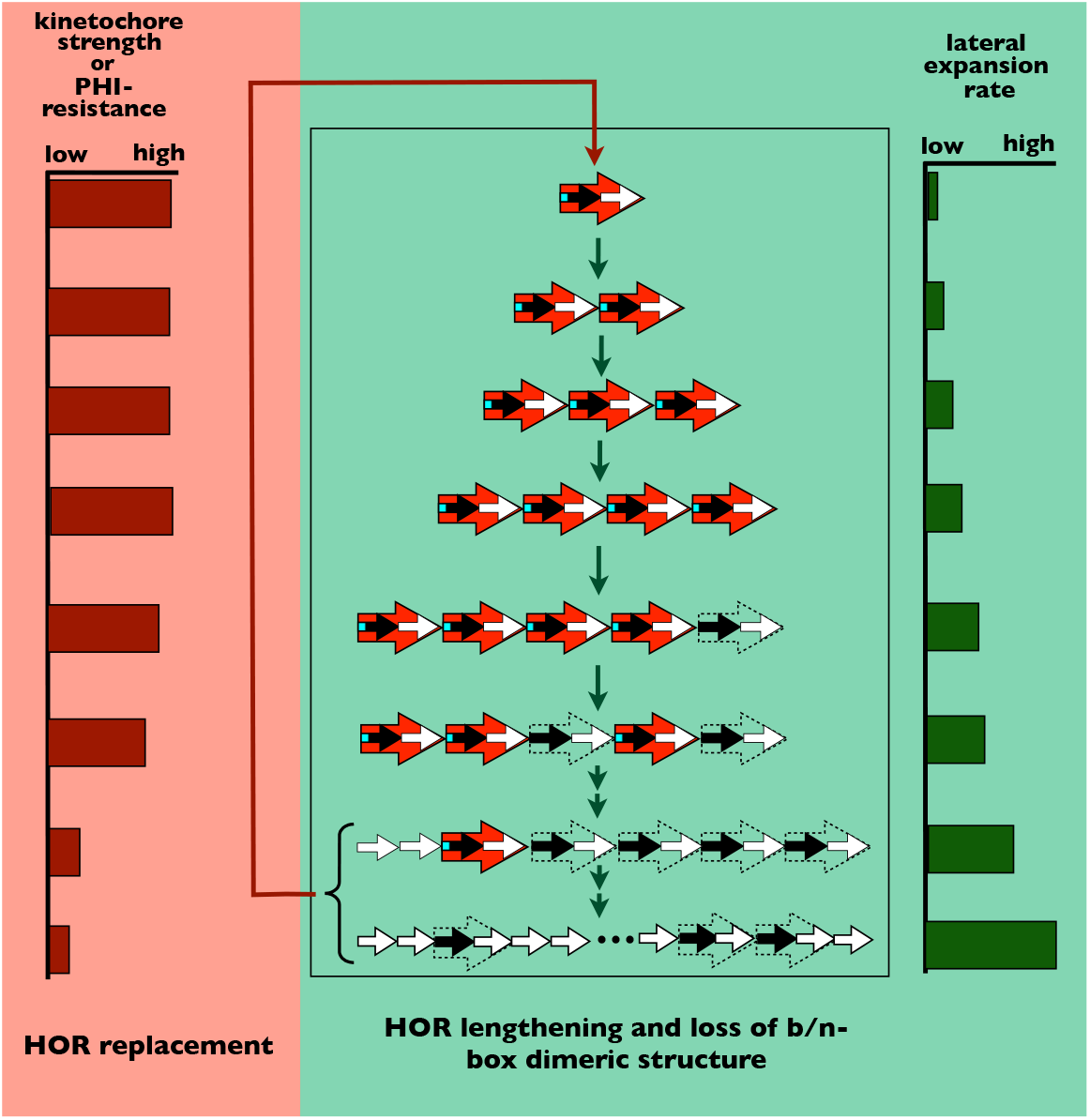
Hypothesis: Life cycle of HORs at human centromeres. Centromeric HORs begin as simple b/n-box dimers (Top) because these short sequences: i) are small enough to be readily recruited to new locations within established HOR arrays via mmBIR with template switching and/or ectopic gene conversion, and ii) produce strong kinetochores and high PHI-resistance. These one-dimer HORs are susceptible to competitive replacement by variant HOR subarrays (having added one or more monomers) that have a lateral expansion advantage due to: 1) being longer (more monomers per HOR) or having reduced density of b/n-box dimers (that reduce recruitment of CENP-B per HOR). Initially HORs become longer exclusively by adding modular b/n-box dimers because loss of even one b-box monomer per HOR leads to a strong reduction in kinetochore strength and/or PHI-resistance (Supplemental Figure S13-S14). Once sufficiently long, HORs begin to also expand by adding (one at a time) single monomers and/or change composition via single loss-of-function mutations within a b-box (so that the b-box no longer binds CENP-B) because these single changes in longer HORs are expected to have only a small effect on kinetochore strength and PHI-resistance (Supplemental Figure S13-S14). These new HOR variants with a lateral expansion advantage are expected to spread among chromosomal lineages due to ectopic gene conversion between homologs and/or BIR with template switching between homologs. Once substantial modular b/n-box modular structure is lost (bottom of figure), the HOR can be invaded and displaced by a short HOR containing a single b/n-box dimer that produces high kinetochore strength and high PHI-resistance (Figure 9, left panel). This replacement (red bottom-to-top arrow) by a single b/n-box dimer occurs via centromere drive favoring lineages with HOR arrays producing stronger kinetochores, and/or lineage spreading (via ectopic gene conversion or BIR with template switching) followed by the dimer’s accumulation within newly invaded lineages due to its strong PHI-resistance.

The HOR life cycle begins with a new HOR array containing tandem copies of a short HOR (a b/n-box dimer). The new HOR is short because this small size is more readily recruited to another chromosome’s active HOR array via ectopic gene conversion and/ or mmBIR with template switching (as described in a previous section). The new HOR next increases in size (adds monomers) by adding new b/n-box dimers (Figure 11), but not by adding other monomer configurations that reduce modular b/n-box dimeric structure, e.g., they do not add single monomers. These expanded HORs replace their shorter ancestors because they retain full b/n-box modular structure (no loss in kinetochore strength nor PHI-resistance) while being able to capture the switch-point of the HOR array (Figure 7) due to their higher lateral expansion rate of longer HORs (Figure 10). Once sufficiently long, HORs continue expanding by sometimes adding monomers that disrupt modular b/n-box dimeric structure (Supplemental Figure S14), e.g., adding single monomers with or without a b-box. They also begin to accumulate mutations in b/n-box dimers that block binding by CENP-B (Supplemental Figure S13). These changes occur gradually: one-at-a-time. This slow, incremental change prevents: i) a strong reduction in kinetochore strength which would lead to their removal by centromere drive (Supplemental Figures S13 and S14), and ii) a strong reduction in PHI-resistance which would prevent them from capturing and retaining the switch-point (Figure 9 [right panel]).

Once an HOR becomes sufficiently degraded in modular b/n-box dimeric structure, it is expected to recruit substantially reduced levels of CENP-C (50% less; Supplemental Figure S11), which produces strongly reduced kinetochore strength and PHI-resistance. The lower PHI-resistance enables a newly invading subarray (of a 1-dimer HOR) to replace the degraded HOR despite its slower lateral expansion rate (Figure 9 [left panel]). The new HOR would be expected to spread beyond its lineage or origin via lineage-spreading. In addition, once the new subarray became sufficiently large within the centric core, it would be expected to generate a substantially stronger kinetochore (leading to a centromere drive advantage): causing lineages with the subarray to replace those without it.

Lineage-spreading would be expected to begin while the new array was too small to produce a centromere drive advantage. One result of this timing is that –once centromere drive began– the new array would have feasibly spread to multiple lineages. This spreading prior to the onset of centromere drive would lead to a “soft selective sweep” (Prezeworski et al. 2005) via centromere drive with little or no footprint of strongly reduced flanking genetic diversity: so there would be little molecular signature a recent episode of centromere drive.

### Intransitive competition is predicted to causes perpetual, rapid, and punctuated evolution at centromeric HORs

Why is evolution at active centromeric sequences so fast and punctuated that current HORs on chimp and human orthologous chromosomes are far more diverged than other regions of the genome (>> 1.2%; Brittan 2002) and commonly do not appear to coalesce to the same recent ancestral sequence (Jorgensen et al. 1992; Archidiacono et al. 1995; Warburton et al. 1996)? This rapid sequence turnover is feasibly explained by a CENP-B-based competitive intransitivity in which A < B, B < C, and C < A (Figure 15). The “A” state is a short HOR made up of a single b/n-box dimer. The “B” state is a moderately longer HOR that retains modular b/n-box dimeric structure. This somewhat longer HOR is formed by two steps as it became an expanded descendant of an A-state dimeric HOR: i) the recruitment of a new b/n-box dimer (via ectopic gene conversion or mmBIR with template switching) into the shorter HOR array (Figure 12), and ii) tandem expansion of the longer HOR within the shorter HOR array until it captures the switch-point and eventually occupies the entire active centromeric array (due to its longer-length-induced faster lateral expansion rate; Figures 7, 10, 13). Steps (i) and (ii) are repeated to make successively longer B-type HORs. The “C” state descends from a B-type HOR once it becomes sufficiently long (Supplemental Figures S13-S14). C-type HORs continue to grow in length (add more monomers) and incrementally lose their modular b/n-box dimeric structure by single mutations in b-boxes and/or incorporation of single monomers (both of which increase lateral expansion rate). Once there is sufficient loss of modular b/n-box dimeric structure, the C-type HOR becomes susceptible to replacement (Figures 12,13) because it recruits substantially less CENP-C (Supplemental Figure S11) and produces: i) weak centromeres that are at a disadvantage in centromere drive and ii) lower PHI-resistance. This susceptibility eventually leads to its replacement by an A-type HOR (Figure 9 [left panel]).

**Figure 15.**
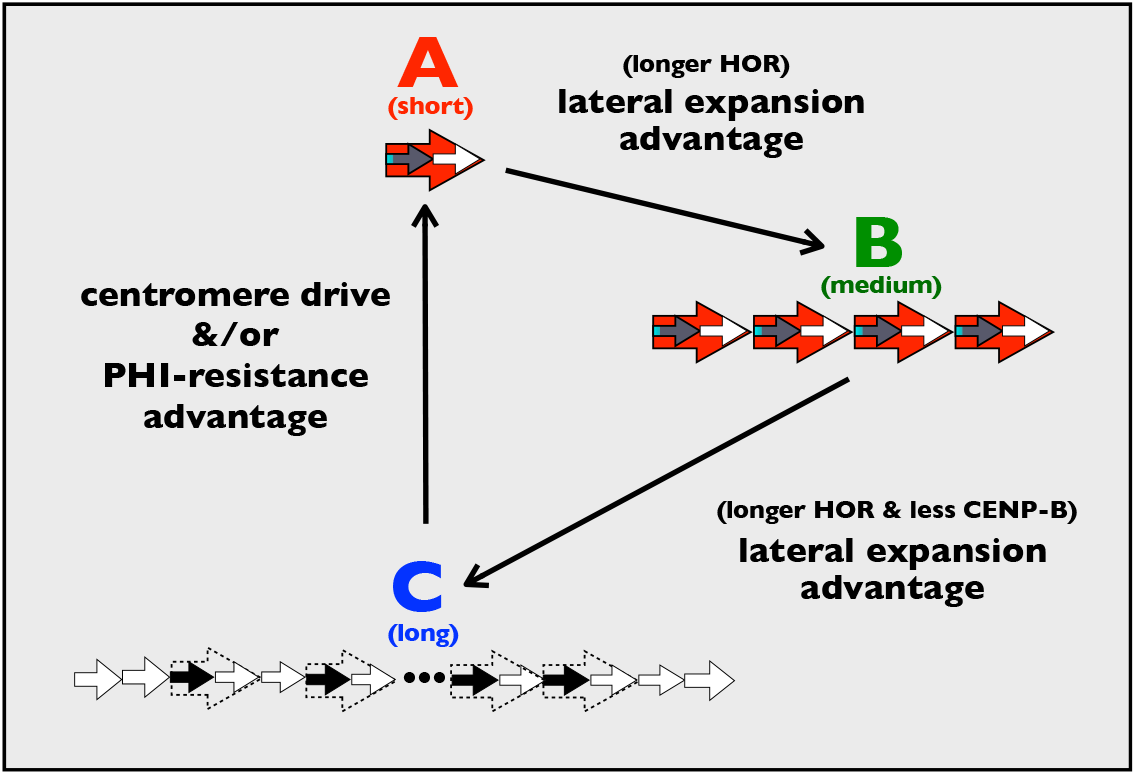
Hypothesis: Intransitivity leads to perpetual, and sometimes punctuated, evolution at centromeric HORs. **A→B.** A new, lengthened HOR that has added one or more b/n-box dimers by transposition (via ectopic gene conversion or mmBIR with template switching, see Figure 10) has a higher lateral expansion rate compared to its parental HORs (see Figure 10): this expansion advantage causes the longer HOR to competitively displace its progenitor HORs (see Figures 6,12-13). Mitotic, non-crossover recombination (via gene conversion or BIR with template switching) between homologous centromere lineages (lineage spreading) enables this transition to spread globally. **B→C.** Once HORs are sufficiently long, they can incrementally lose modular b/n-box dimeric structure by: i) mutations in b-boxes (that prevent binding by CENP-B), and/or ii) recruiting single n-box monomers (see Supplemental Figures 13-14). Both of these features reduce the density of binding by CENP-B (which in yeast reduces fork-stalling/ collapse). Reduced CENP-B binding (and additional HOR lengthening) is predicted to increase lateral expansion rate and provide an intra-array competitive advantage to these HORs: thereby causing them to competitively displace their parental HORs with higher b/n-box dimeric modular structure (and/or shorter length). These HORs with reduced b/n-box dimer modular structure (and/or increased length) can spread globally to all lineages via lineage-spreading. **C→A.** Once HORs lose substantial modular b/n-box dimeric structure, they recruit less CENP-C (Supplemental Figure S11). This reduction is expected to: i) reduce PHI-resistance (see Figure 2 and the main text section: High density of b/n-box dimers within HORs is predicted to increase PHI-resistance and to be favored in intra-array competition), and ii) produce substantially weaker kinetochores (Supplemental Figure S11). Low PHI-resistance is expected to causes long HORs with degraded b/n-box modular structure to be competitively displaced by invading (transposed), single b/n-box dimer HORs (Figures 12-13, 8 [left panel]). The newly invaded b/n-box dimer HOR can spread globally to all lineages via lineage-spreading. Higher kinetochore strength is predicted to provide an additional route to the spread of the single b/ n-box dimer HORs: a segregation advantage via centromere drive. The C→A transition represents a form of punctuated evolution. Arrow key as in Figure 10.

The intransitivity (A < B, B < C, C < A) would be expected to generate a cycle of perpetual evolution at active centromeric arrays (Figures 14,15). The fact that base substitution rates are ~1000-fold higher during BIR replication (Sakofsky et al. 2012) would be expected to speed HOR sequence divergence with time compared to other genomic regions. The invasion of type-C HORs arrays by unrelated A-type dimers (Figures 9 [left panel] and 11) – which would subsequently grow into longer HORs– would lead to the observation that HORs on orthologous chimp and human chromosomes are frequently descended from different ancestral sequences (Jorgensen et al. 1992; Archidiacono et al. 1995; Warburton et al. 1996). The hypothesis that unrelated HORs can and do invade and eventually replace each other (Figures 9, 12, 13) is supported by the observation that the flanking, nonactive HORs arrays observed on some chromosomes are commonly distantly related to the active HOR arrays, and to each other (see Figure 8 in the companion paper, Rice 2019).

### Continuous turnover at centromeric repeat arrays does not require HORs nor CENP-B-binding at b-boxes

The cycle of perpetual evolution described in the above section is driven by the influence of HOR length (number of monomers), composition (level of b/n-box dimeric structure within the HOR), and the phenotypic effects of CENP-B at centromeres (it can double the recruitment of CENP-C to nucleosomes and plausibly influence the rate of fork-collapse as well as the level of PHI-resistance). However, rapid and perpetual evolution at centromeric repeats is feasibly expected to operate without CENP-B binding at centromeres (i.e., without the intransitivity this binding can generate; Figure 15), and without the presence of HORs.

Evidence for HOR replacement outside the context of the CENP-B-associated intransitive competition (Figure 15) can be foun d at chromosome 15. Th is chromosome has two flanking, inactive HOR arrays that completely lack b-box monomers (both to the left [p-arm side] of the active HOR array when viewed in the UCSC genome browser). The levels of standing genetic diversity (for SNPs and TEs, from the UCSC genome browser diversity tracks) indicate that the flanking array most distal to the active HOR array is the oldest, and plausibly replaced by the other (middle) HOR array, which in turn was replaced by the currently active HOR array that is rich in b/n-box dimers. The replacement of one flanking HOR array by the other flanking HOR would have taken place without any influence of CENP-B.

A plausible scenario for this replacement is that the older HOR array was invaded by a short repeat of one to a few monomers which then expanded to form a small repeat subarray (Figures 12-13). All else being equal, this short repeat would be predicted to have a slower lateral expansion rate (Figure 10), but could nonetheless invade the longer HOR array if it had substantially higher PHI-resistance (Figure 9 [left panel]). One factor predicted to contribute to PHI-resistance is the interaction of an HOR with gene products from genetic variation for for the suppression and enhancement of PEV (Su[var] and En[var]; Figure 3). Because in flies (i.e., *Drosophila*, where almost all the PEV studies have been conducted) at least 150 genes are known to contribute to these suppressor and repressor phenotypes (reviewed in Elgin and Reuter 2013), and because high levels of PEV genetic variation has been found to be segregating in natural populations (most of which does not map to these 150 genes; Kelsey and Clark 2017), there appears to be exceptionally high amounts of polygenic variation influencing PEV: and therefore plausibly influencing PHI-resistance (Figure 3).

The replacement of one HOR array on chromosome 15 by another array when both arrays do not bind CENP-B could have occurred because the replacement HOR (or a smaller version of it that initially invaded) better recruited PEV-influencing gene products that substantially increased PHI-resistance (Figure 9 [left panel]). Alternatively, the short HOR array might fortuitously bind a new protein that is unrelated to centromere function –or more tightly bind a protein already binding centromeric HORs– causing it to have substantially higher fork-stalling/collapse/BIR and hence a higher lateral expansion rate despite having fewer monomers per HOR (Figure 7). The key feature in both scenarios is the evolution of a new repeat sequence with superior PHI-resistance and/or lateral expansion rate: features that feasibly require neither CENP-B binding at the centromere, nor HOR structure of the repeat.

### Interactant ‘shifts’ and ‘retreats’ are also predicted to drive perpetual evolution of centromeric repeats

For generality in this section I will not presume HOR structure nor CENP-B-binding at centromeres: but the same logic will apply when these features are present. A new sequence is expected to replace an extant centromeric repeat array by chance, when it fortuitously originates within the sequence spanning the switch-point position (Figure 5), or when it confers a within-array advantage: higher PHI-resistance (Figure 8) and/or higher lateral expansion rate (Figure 7). Opportunity for the within-array advantage route would be expected to diminish with time as evolution incrementally recruited sequences closer to the optimum for PHI-resistance and lateral expansion rates: unless these phenotypes are mutually opposing (as appears to be the case with CENP-B binding) or because the optimum is continually changing.

One reason the optimum may be continually changing is that a subset of molecules that interact with the sequences of the centromeric repeats are continually evolving in a context that is independent of centromere functioning. These molecules are assumed to have cellular functions that are unrelated to the centromere (their primary functions), yet they fortuitously bind centromeric DNA. I will refer to these molecules collectively as ‘interactants’. Their continual evolutionary change is expected because the interactants are continually evolving in the context of their primary cellular functions (and via genetic drift). In this scenario, centromeric repeat sequences have evolved to interact with non-centromere proteins (e.g., bind them) because this interaction increases PHI-resistance and/or lateral expansion rate. In response, the optimal sequence at the centromeric repeats would slowly change over time as the primary cellular functions of the interactants independently evolve. I will refer to this process as an ‘interactant shift’.

Alternatively, the optimal sequence at centromeric repeats may rapidly evolve due to an ‘interactant retreat’. In this case, as a new repeat sequence (that binds a new interactant molecule and gains a lateral expansion or PHI-resistance advantage) expands to many thousands of copies within the centromere of one or more chromosomes, it would reduce the availability of the interactant in its primary cellular functioning. This drawdown would select for increased selectivity of the interact for its primary function, which would reduce its association with the centromeric repeats. In turn, the increased selectivity of the interactant molecule would select for a new sequence at the centromeric repeat that better competed for the interactant –or it would favor a new sequence that binds/associates with a new and different interactant molecule. These interactions would generate a a form of chase-away antagonistic coevolution (Parker 1979; Rice and Holland 1997; Holland and Rice 1998): a positive feedback loop that can drive perpetual evolution.

There is empirical evidence that satellite DNA binds a diverse array of DNA-binging proteins (e.g., transcription factors) and influences the cellular functioning that depends on these proteins. In flies, Lemos et al. (2010) found that polymorphic Y chromosomes substantially influence the transcription rate of thousands of genes in XXY females that do not express structural Y-linked genes. They also found evidence that satellite repeats on the Y bind DNA binding proteins that influence gene expression.

If the polygenic variation influencing PHI-resistance and/or lateral expansion rate is continually turning over through time (due to genetic drift, interactant shifts, and/or interaction retreats), there would be persistent selection for new centromeric HOR sequences that work better (increase within-array competition) in the changing genetic background. This evolution would generate a continuous turnover of repeat sequences at centromeres as new sequences invade and expand due to their superior PHI-resistance and/or lateral expansion rate. A possible example of an interactant molecule is pJα. This protein binds centromeric DNA in humans yet has no known centromeric function (Gaff et al. 1994; Romanova et al. 1996).

### The Y chromosome HOR is predicted to be exceptional

The HOR on the male-limited, human Y chromosome differs from those on the X and autosomes in many ways: i) it is exceptionally long (34 monomers) compared to the longest active HOR on the X and autosomes (19 monomers on chromosome 4), ii) it has no b-boxes whereas these are found on all of the active HORs on the X and autosomes, iii) it binds 50% less CENP-C and is exceptionally weak, with the highest mis-segregation rate of all of the chromosomes (Fachinetti et al. 2015), iv) it is never exposed to centromere drive because it is male limited, and v) it is virtually always hemizygous, a condition that prevents sequence exchange between homologs.

Because the Y chromosome is male limited, the C-to-A transition (long HOR to short HOR) shown in Figure 15 is inoperative via centromere drive. Although centromere drive cannot fuel this transition, it might still occur through lineage-spreading of a short HOR with high PHI-resistance. However, hemizygosity of the male-limited Y chromosome prevents lineage-spreading and thereby impedes the C-to-A transition (and all other transitions in Figure 15) except via genetic drift. As a consequence of the Y chromosome’s male-limited transmission and hemizygosity, most of its centromeric HOR evolution is expected to be driven by intra-array competition (i.e., by faster lateral expansion rate and higher PHI-resistance) in combination with genetic drift among lineages.

The absence of centromere drive is expected to contribute to the evolution of exceptionally long HORs on the Y chromosome because the evolution of progressively longer HORs (transition B-to-C in Figure 15) is not truncated by centromere drive (nor increased PHI-resistance in combination with lineage spreading) replacing a long HOR with a short one (transition C-to-A). Correspondingly, the Y has evolved an exceptionally long HOR.

The lack of centromere drive on the Y chromosome would also preclude one selective mechanism for recruiting b-box monomers (that make stronger centromeres) when these began spreading within the lineage of the great apes. Hemizygosity, male-limited transmission, and lack of recombination markedly reduces the effective size of the Y chromosome (Wilson Sayres et al. 2014), and this reduction would have lowered the efficacy of selection for the stronger centromeres that b-boxes would produce (Fachinetti et al. 2015). Autosomes and the X (in females) have a homolog present in the same cell which creates an opportunity for ectopic gene conversion and/or BIR or mmBIR with template switching, to move sequences between homologs. This opportunity is absent for the hemizygous Y chromosome (note that lineage-spreading cannot occur in rare XYY individuals [about 1 in 1000 human males; Morris et al. 2008] because the Y chromosomes in this karyotype are identical copies inherited from the same male). In sum, impeded natural selection and the absence of both centromere drive and lineage-spreading would be expected to strongly interfere with the recruitment of CENP-B-binding b-boxes to the Y chromosome after they became established within the great ape clade. Correspondingly, the Y is the only chromosome that has not evolved a centromeric HOR that contains b-box monomers despite the higher functioning centromeres that they produce. The weak centromere strength, extreme HOR length and lack of b-box monomers observed at the centromeric HOR on the Y chromosome are consistent with the model of centromeric HOR evolution shown in Figures 14 and 15.

## A Game-of-Thrones at human centromeres

Centromere evolution is a *“A Game of Thrones”* (*GofT*) at the molecular level. Newly arising centromeric HOR lineages resemble regime changes because centromeres are defined by an epigenetic ‘crown’ of CENP-A. New HORs replace old ones when the crown of CENP-A is transferred between them. This transfer can occur: i) within an hereditary line of descent when the new HOR is a mutational modification of the old one that fortuitously arises within the head of the ruling lineage (i.e., at the switch-point within a centromeric HOR array), or ii) between hereditary lines when a related or unrelated ‘usurper’ (new HOR subarray) captures the switch-point of an active centromeric array due to faster expansion of its ‘house’ of tandem repeats to outnumber the old regime (faster lateral expansion) and/or better vanquishes persistent invaders (higher PHI-resistance).

Centromere evolution and the *GofT* also have numerous thematic similarities. Slashing, cutting, and decapitation are predominant interactions in the *GofT*. Similarly, DSBs and fork-collapse (slashing and cutting) and the truncation of sister chromatids during DNA replication (decapitation) dominate the molecular interactions that drive centromere evolution. Frequent promiscuous sex, including that between close relatives, permeates all aspects of the *GofT*. In parallel, frequent promiscuous DNA exchange during copy number expansions (intragenomic reproductive acts) pervades centromere evolution: out-of-register BIR between sister chromatids (close relatives) and ectopic gene conversion and/or mmBIR with template switching (promiscuous procreation of repeat elements). An endless succession of regime changes within and between dynasties, driven primarily by violence (cutting) rather than a hierarchical transfer of power via hereditary succession, is the overarching theme of *GofT*. Similarly at centromeres, epigenetic inheritance leads to a form of perpetual, non-hierarchical regime change. This includes: 1) transfer of the CENP-A epigenetic “crown” from deposed to usurper HORs via cutting (DSBs) and decapitation (mmBIR with template switching of a fork-collapsed [decapitated] sister chromatid) with frequent low genetic similarity between ancestral and descendent sequences, and with deposed HORs exiled to the ‘peripheral, pericentric flanks where they are literally ‘cut to pieces’ by recurrent deletions from SSA-repair of DSBs. Lastly, the huge, flying, fire-breathing dragons that gradually grow from tiny hatchlings into behemoths are a hallmark of the GofT. At the molecular level, they are represented by centromeric HOR arrays that start their lifecycle as tiny, transposed b/n-box dimers (molecular flight to new genomic locations), but gradually grow into behemoths (satellite arrays of many millions of basepairs) with periodic flares of punctuated evolution when new b-n-box dimers invade a different chromosome and usurp their epigenetic CENP-A crowns.

## Conclusions

The new model for centromere evolution developed here is a radical departure from the unequal crossing-over model that is the generally accepted hypothesis for centromere evolution (Smith 1976). A new model is needed because the Smith model does not predict: i) the multiple levels of sequence structure observed at human centromeric HORs, ii) the pattern of centromeric length variation at the sex chromosomes, and iii) the far-larger-than-required size of human centromeric HOR arrays (Rice 2019). The foundation for the new model is a new form of intragenomic differential reproduction (aka molecular drive; Dover 1982) that occurs within centromeric tandem repeats: subarrays with faster lateral expansion rates and/or higher PHI-resistance are favored in intra-array competition, and arrays that form stronger kinetochores are favored in inter-array competition via centromere drive. The new model replaces unequal exchange between sister chromatids in the Smith model with the repair of collapsed replication forks via out-of-register BIR replication. In humans, it predicts rapid, continuous, and sometimes punctuated evolution of centromeric repeats due to a competitive intransitivity between three HOR types: i) single b/n-dimers, ii) small groups of b/n-dimers, and iii) longer HORs with reduced modular b/n-dimeric structure. It also predicts (or is consistent with) all of the patterns and multiple levels of structure observed at human centromeric repeats (Rice 2019). The new model is a straightforward consequence of its four assumptions: i) centromeric HOR arrays are partitioned into a centric core that binds kinetochore proteins and outer flanks that recruit dense cohesin and form the innermost region of the pericentric heterochromatin, ii) the boundary between centric and pericentric chromatin is dynamic and moves inward as the HOR array expands, iii) only the centric core of an HOR array continuously expands via fork-stalling/collapse/ BIR because the pericentric flanks recruit dense cohesin clamps that suppress out-of-register BIR, and iv) the entire HOR array continuously contracts via recurrent deletions (due to SSA repair of DSBs and also rarer longer deletions due to NHNJ repair of pairs of DSBs). There is direct and strong empirical support for assumptions (i) and (ii), while the evidence for assumptions (iii) and (iv) is substantial but indirect and requires further testing. If these assumptions are met, there will be stochastic, bidirectional flow of repeat elements within an array from a central position outward toward the two edges of the centric core. The central array position where the direction of flow reverses (the switch-point) has pivotal evolutionary significance: repeat variants that can capture (i.e., have tandem copies that span) the switch-point will eventually spread to the entire repeat array. The presence of a switch-point generates intra-array selection for sequences (especially in the context of the organization of monomers into b/n-box dimers) that produce phenotypes (higher lateral expansion rate and PHI-resistance) that allow them to capture the switch-point and thereby expand to become the predominant repeat sequence at a centromeric array.

Although the new model directly applies to humans and other great apes, the foundation of the model has wider application for other organisms with regional centromeres: irrespective of the presence of HORs and the binding of CENP-B at the centromeres. As organisms adapt to a continuously changing environment, a persistent turnover of polygenic variation is expected. This turnover would generate ever-changing opportunities for new molecules (interactants) to bind/associate with centromeric repeat sequences and increase their intra-array competitive ability: leading to perpetual evolution of new repeat sequences that take advantage of this changing genetic background. The binding of interactants to centromeric repeat arrays might also reduce their availability for other cellular functions and lead to chase-away antagonistic coevolution that causes continual evolution at centromeric repeats.

## Acknowledgements

I received helpful comments of the early ideas that motivated this paper when I presented a seminar on centromere evolution at the Fred Hutchinson Cancer Research Center, especially from Harmit Malik. Additional comments were provided by Steve Henikoff on an early draft of the manuscript and by Urban Friberg and Andrew Stewart on a near-final draft of the manuscript. Copy-editing assistance was provided by Kathryn Schoenrock.

---

**Supplemental Figure S1.**
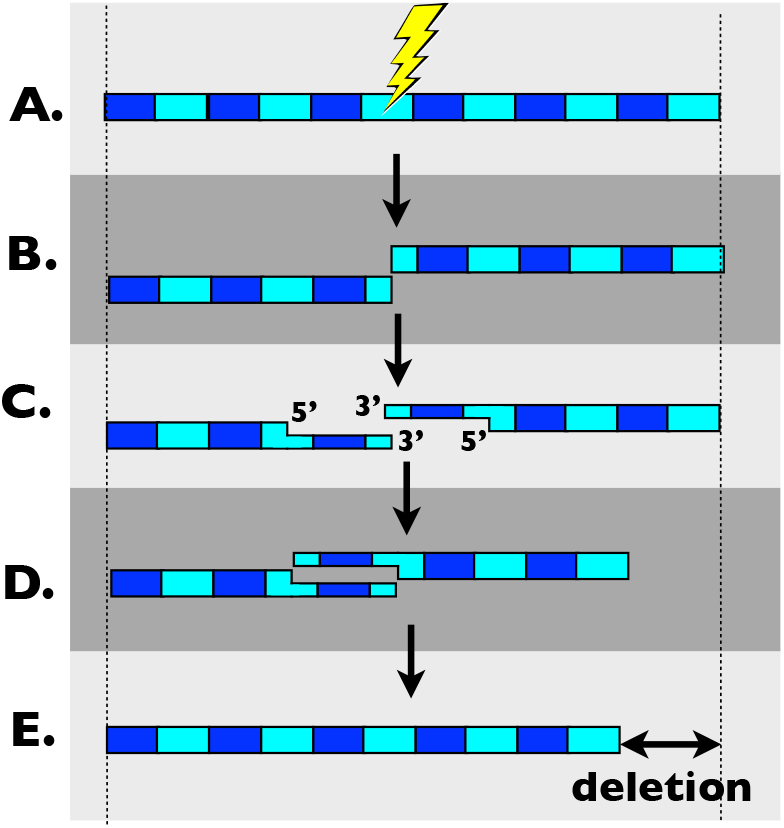
SSA repair of a DSB (lightening bolt) leads to a deletion of a single repeat unit in a tandem array. **A-B.** A DSB occurs in a length of DNA composed of a tandem array of two monomers (blue and turquoise rectangles). **C.** Resection of the 5‘ ends of each DNA strand. **D.** Once resection uncovers homology, strand annealing occurs followed by ligation of the two single strand breaks. **E.** A deletion is produced after repair, (deletion length = length of one repeat unit). If the DSB-generating agent causes damage to the strand ends that is removed prior to repair (e.g., via exonucleases), then the removed bases will be replaced via gap-filling after annealing.

**Supplemental Figure S2.**
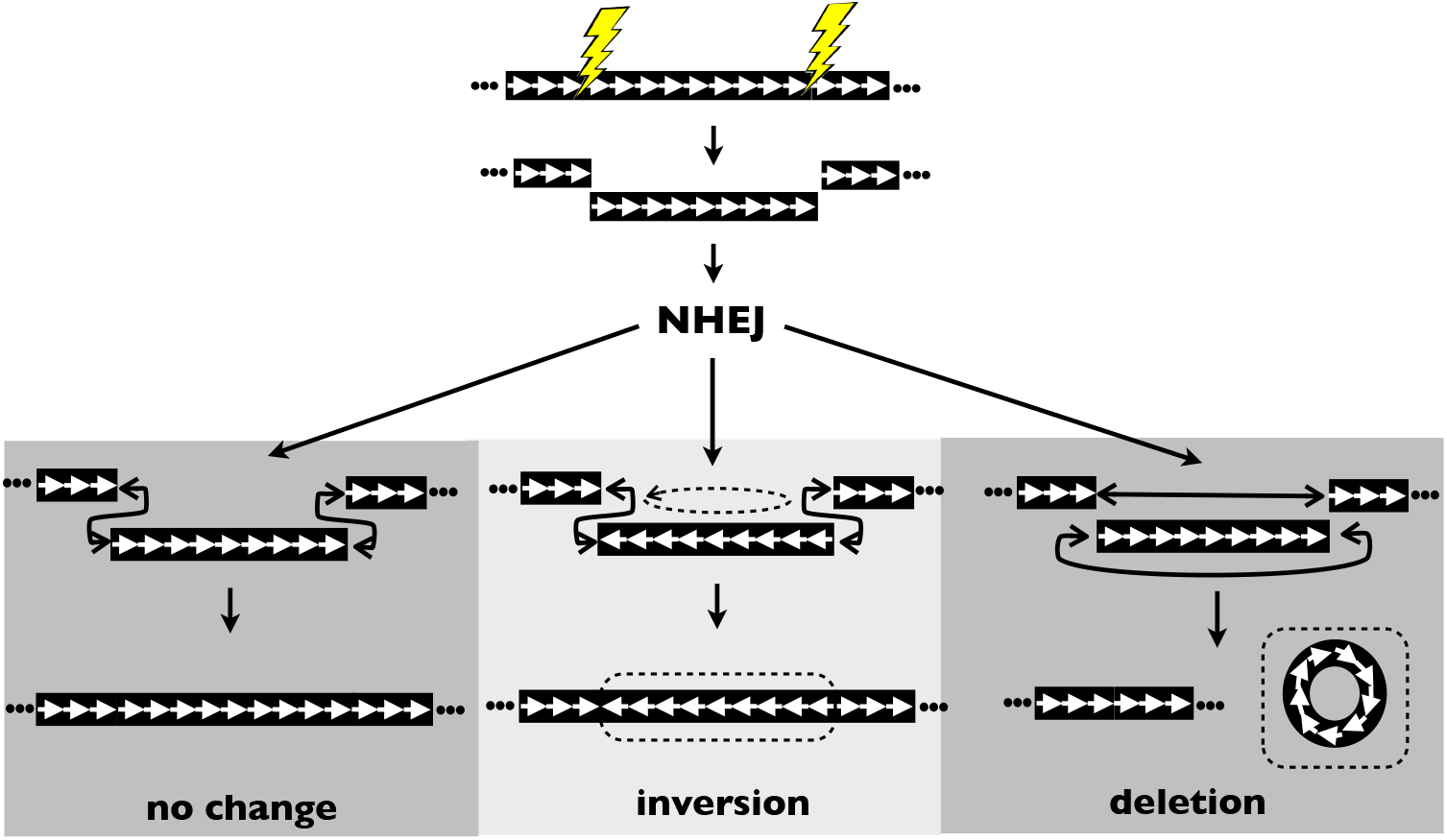
Deletions are one outcome during NHNJ repair of pairs of DSBs.

**Supplemental Figure S3.**
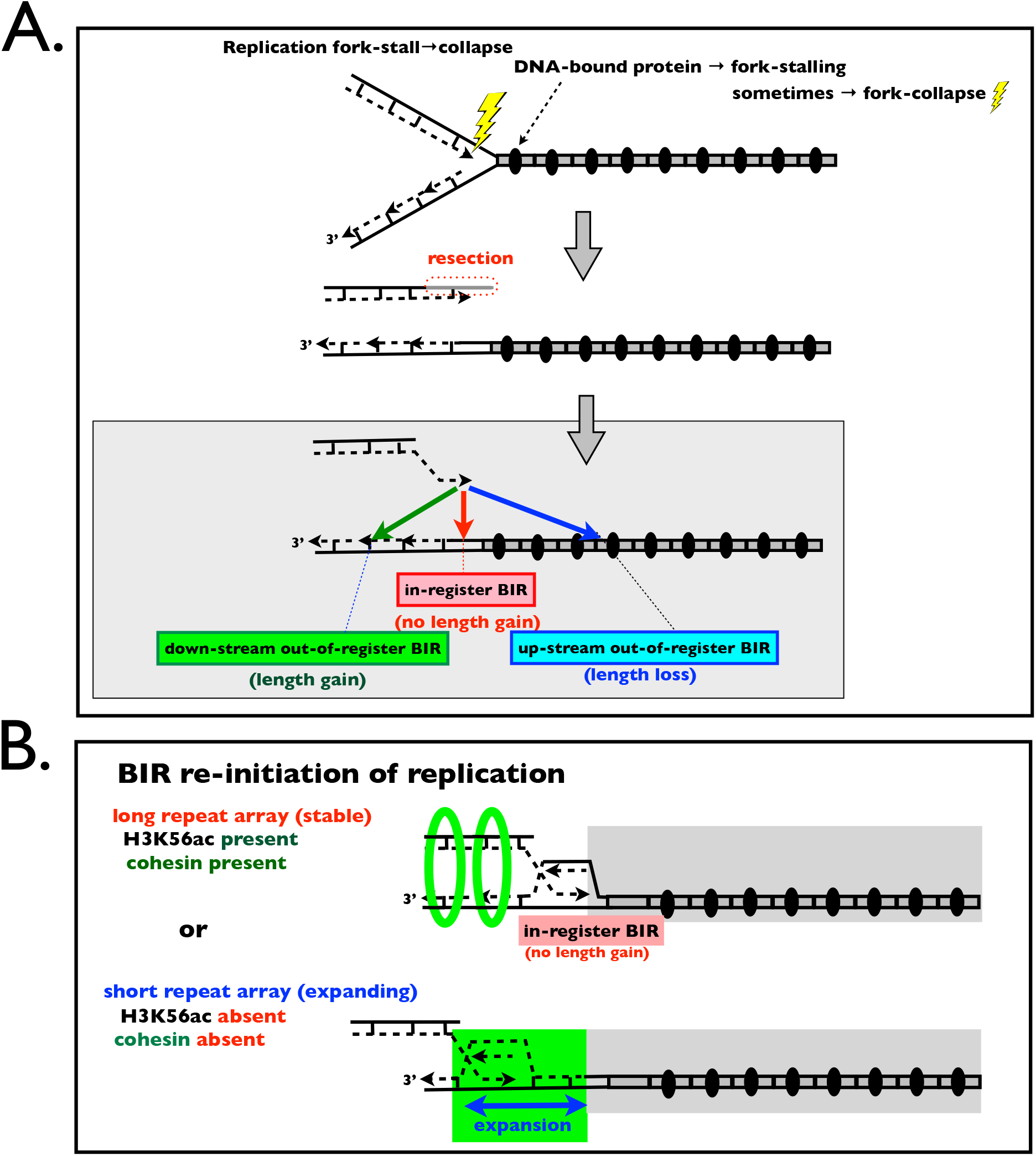
Expansion in yeast rDNA repeats via fork-collapse and BIR repair (see Kobayashi & Ganley 2005). **A.** BIR repair of a collapsed replication fork in a tandem repeat can increase, decrease or have no effect on the length of the resected chromatid. **B.** In yeast, long arrays are stable because dense cohesin is present and prevents out-of-register re-initiation of DNA replication. In short arrays, cohesin is absent or rarefied, permitting out-of-register re-initiation of DNA replication, which is biased toward downstream repeats, and hence, on average, leads to array expansion in length of the upper chromatid.

**Supplemental Figure S4.**
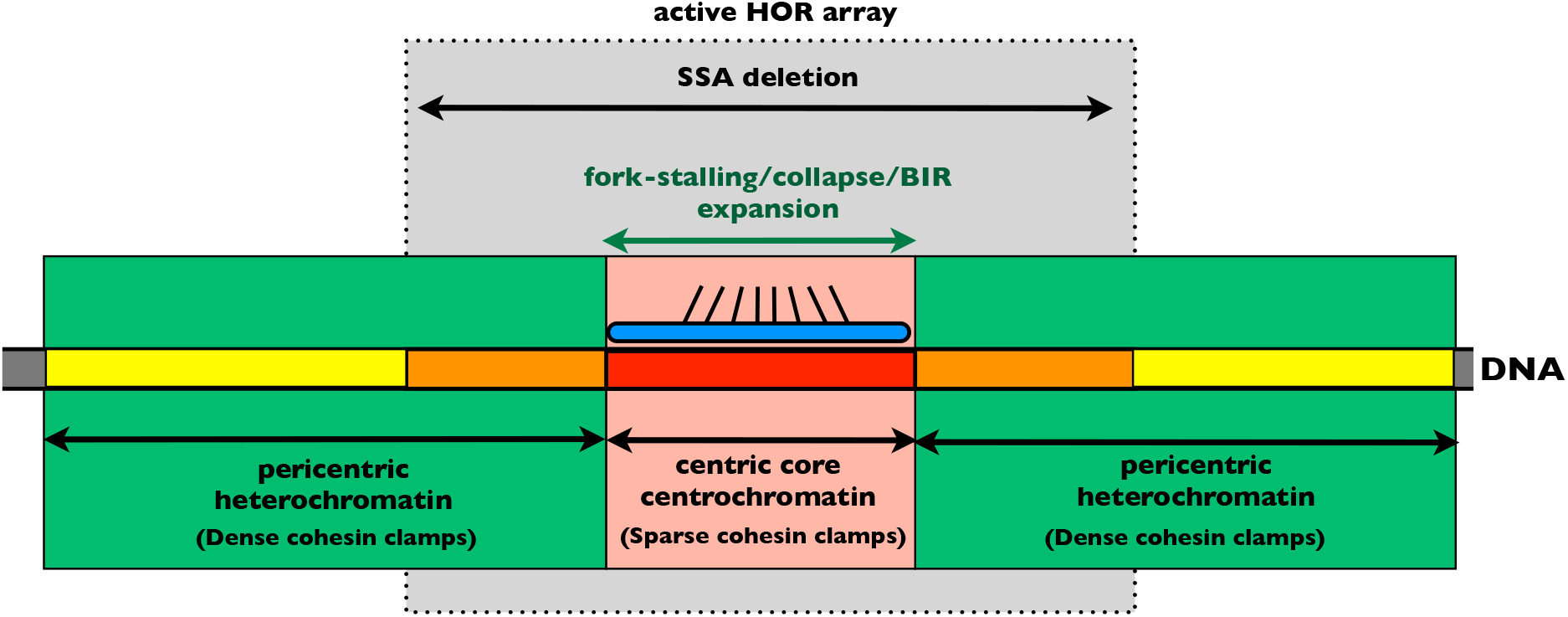
Only the inner, centric core of the active HOR array (dark red central rectangle, ~ one third of the HOR array, composed of centrochromatin) that binds the kinetochore (blue oval) has the attributes required for persistent expansion due to out-of-register BIR, i.e., this region has i) fork-stalling and collapse due to bound CCAN proteins and/or DNA/RNA polymerase collisions, and ii) reduced cohesin recruitment due to the absence of heterochromatin. The outer, pericentric flanks of the HOR array (orange rectangles) are heterochromatic and recruit dense cohesin clamps (green region), as does the more distal pericentric heterochromatin (yellow rectangles; primarily composed of unorganized monomeric repeats) surrounding the active HOR array. Deletion via SSA repair of DSBs is expected to occur across the entire HOR array.

**Supplemental Figure S5.**
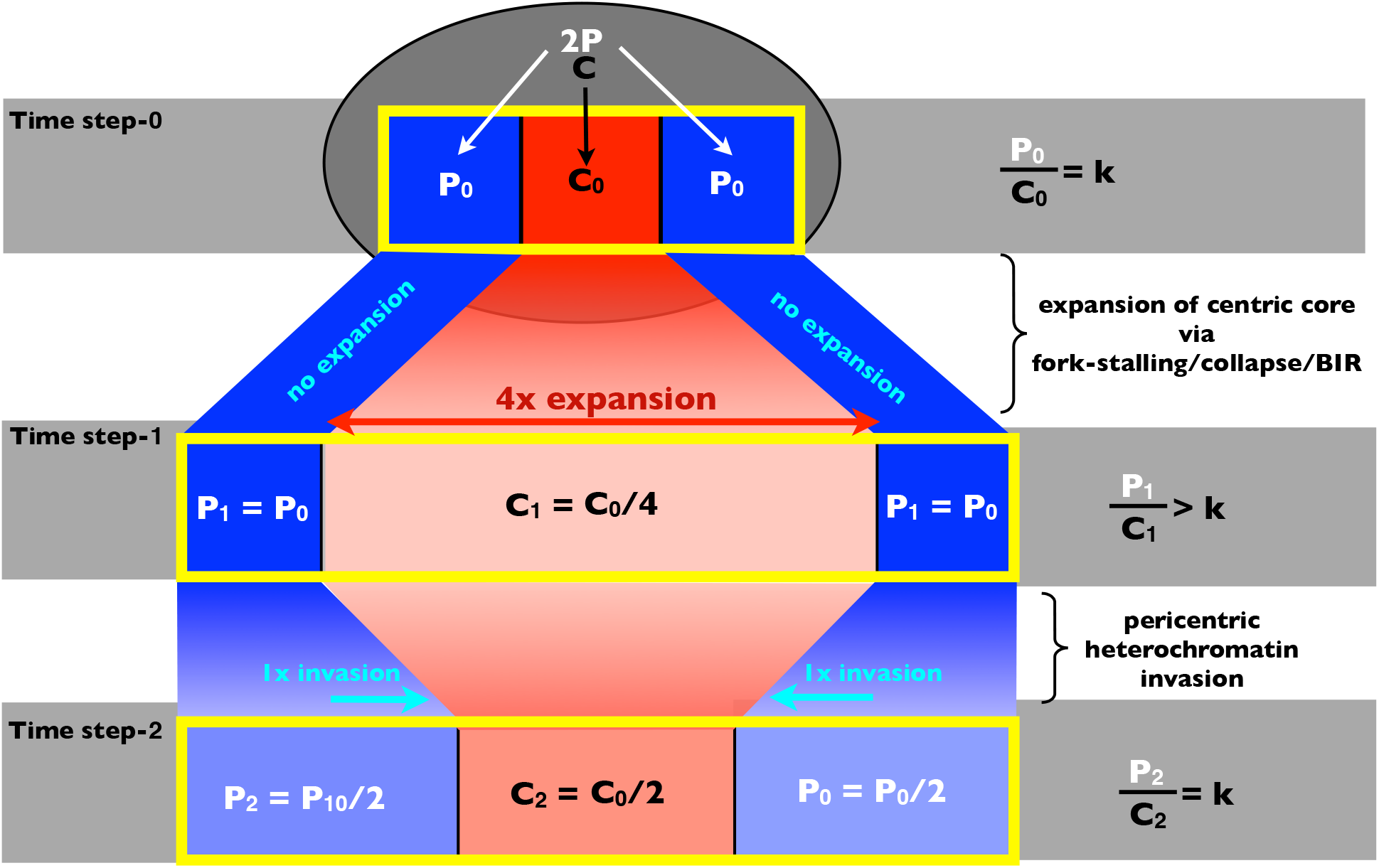
A simple model for the constant proportional size of the centric core as the HOR array expands or contracts in size. The three yellow rectangles represent a centromeric HOR array through three time steps, with blue regions representing the two pericentric flanks and the red region the centric core. The dark ellipse at the top of the figure represents the nucleoplasm in the region surrounding the centromeric HOR array. **Time-step-0**. Assume that the boundary between the centric core and the pericentric flanks (Centric-Pericentric-boundary) of a centromeric HOR array is formed when chromatin assembly spreading outward from the centric core (centrochromatin) is exactly counterbalanced by inward spreading of the pericentric flanks (heterochromatin). Further assume that each of these spreading rates is controlled by a different rate-determining steps that depend on localized concentrations (P and C) of Factor-P (localizing to the pericentric flanks) and Factor-C (localizing to the centric core) that control the spread of the percentric flanks and centric core, respectively. Spreading rates are equal (and each Centric-Pericentric-boundary is stabilized) when P = kC (or equivalently P/C = Ratio_P/C_ = k), where k is a constant. **Time-step-1**. Fork-stalling/collapse/BIR (in the absence of dense cohesin clamps) causes the centric core alone to expand, and this dilutes the concentration of Factor-C from C0 to C0/4 within the centric core. An exaggerated 4-fold expansion in one time step is shown to make this step, and the following inward flow of the Centric-Pericentric-boundary, more visually apparent. **Time-step-2**. The reduced concentration of Factor-C within the centric core allows the pericentric flank to progressively invade the centric core until RatioP/C = k. This cycle of centric core expansion and proportional invasion of the pericentric flanks would explain: i) the approximate constant relative size of the centric core across HOR arrays of vastly different sizes and sequence (as found by Sullivan et al. 2011; Ross et al. 2016), and ii) the expansion of the proportionate size of the centric core when CENP-A concentration (a feasible candidate for Factor-C) was experimentally increased (as found by Sullivan et al. 2011). This same reasoning would apply to the context of a large deletion removing a substantial part of a centromeric HOR array: the size of the centric core (in bp) is expected to decline but its proportional size should remain unchanged.

**Supplemental Figure S6.**
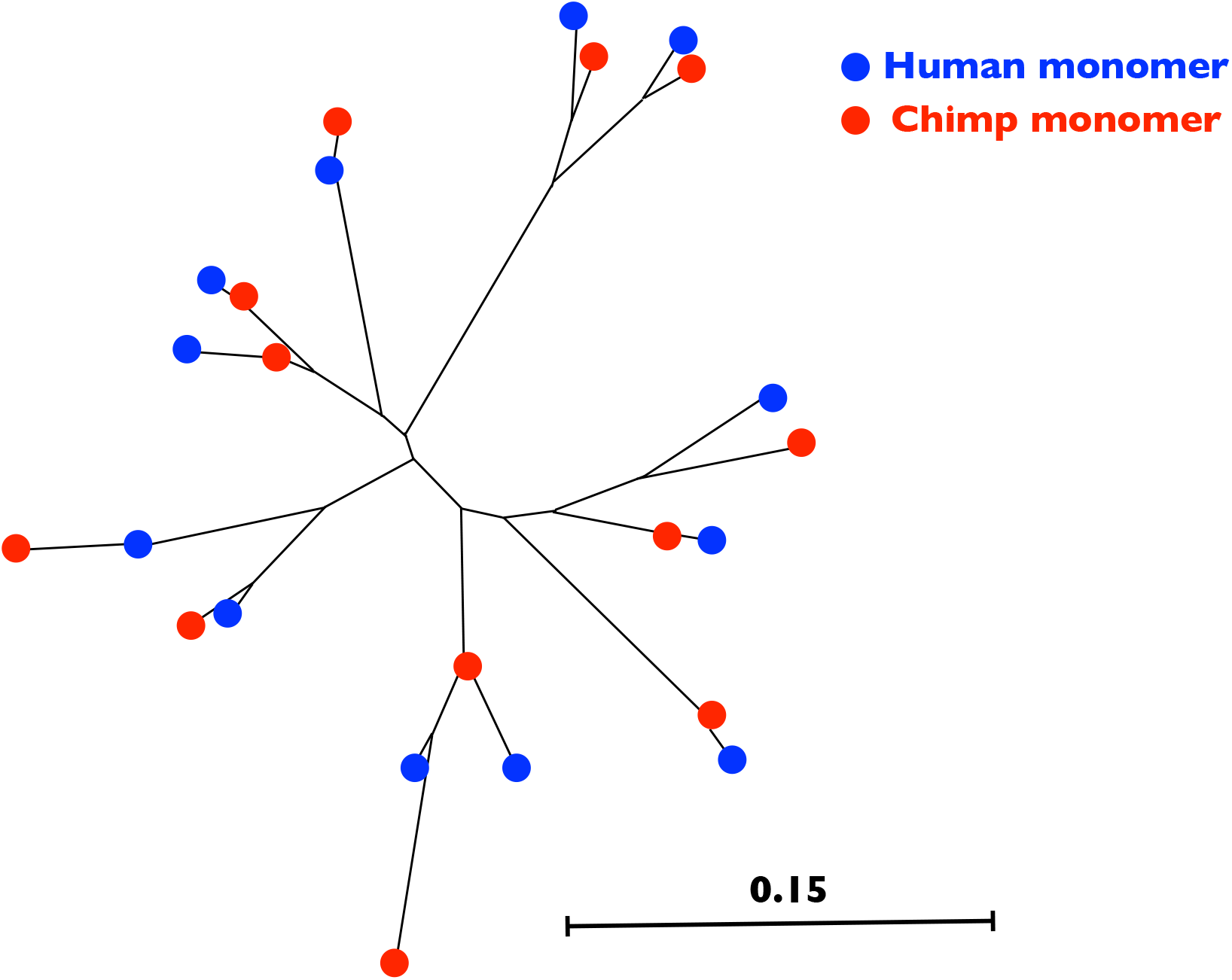
A neighbor-joining tree of the 12 monomers from the X-linked centromeric HOR arrays of humans and chimps. Human sequence is from Supplemental Table 1 of the companion paper, Rice 2019. The chimp sequences were obtained from PacBio reads using the same procedures described in Rice (2019). The average divergence between the sequences (ignoring the 14 bp insert in one of the human monomers and summing across the two consensus 12-monomer HORs) is 6.7% (136 mismatches out of 2038 bp) and the Jukes-Cantor divergence is 6.2%.

**Supplemental Figure S7.**
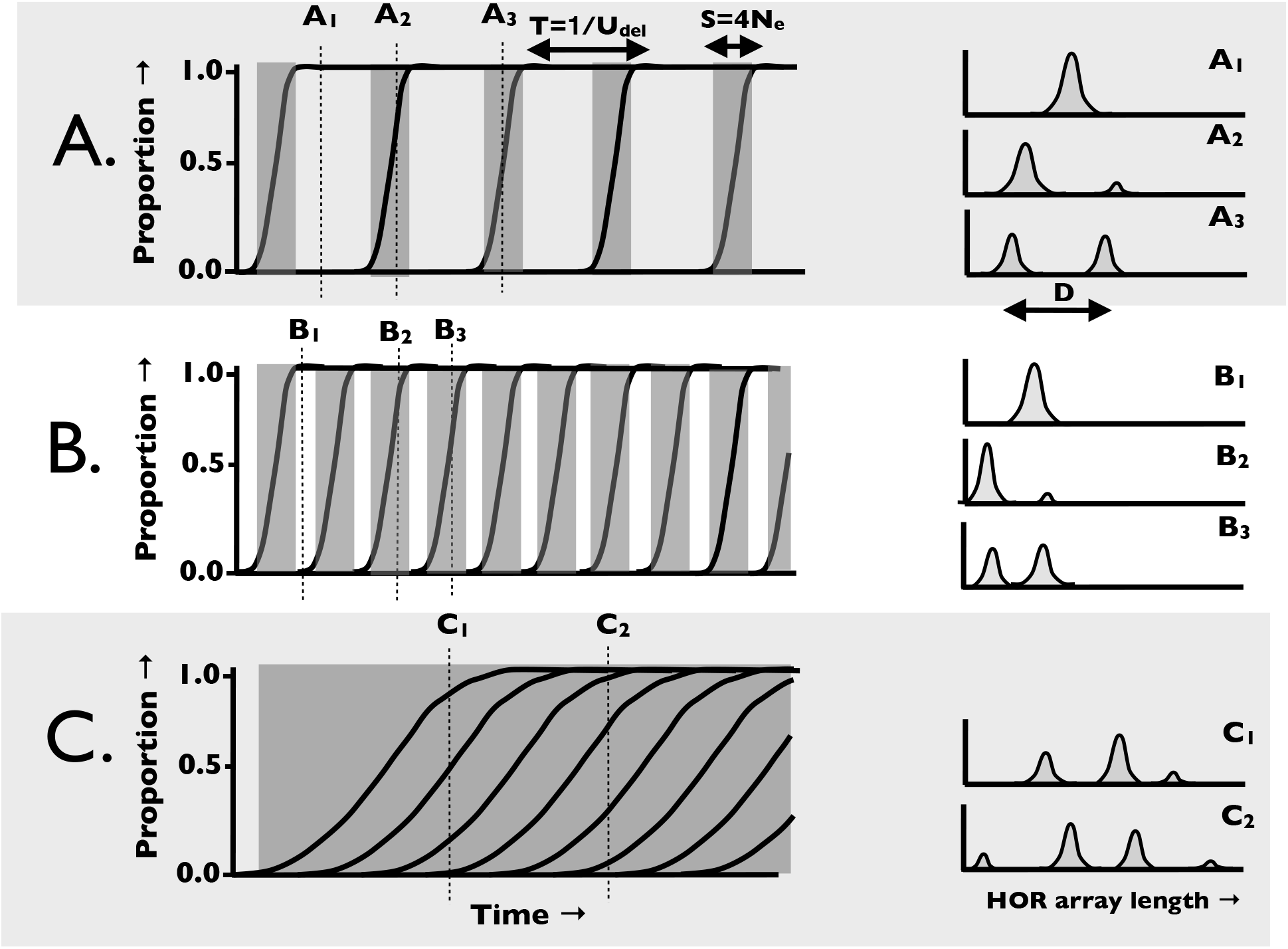
A simplified graphical model of genetic drift of large deletions (of average size D). For neutral mutations: i) the average time between allelic replacements (T) is 1/U_del_ generations where U_del_ is the mutation rate to large neutral deletions, ii) the average time from mutation to fixation (S) is 4N_e_ generations, where N_e_ is the effective population size (Kimura 1983). In this simplified analysis, I ignore sampling error in D, T and S and use their expected values. Allele frequency graphs are on the left side of the figure and size distribution graphs on the right side. Alleles are large deletions that (when not lost early-on by sampling error) grow back to larger size by fork-stalling/collapse/BIR. Stochastic variation in growth of alleles (and deletions from repair of DSBs via SSA repair) generates size variation around their average size. Shaded regions denote periods of polymorphism for alleles. Size graphs depict the distribution of allele sizes at the time points indicated at the top of each allele frequency graph. **A.** Reference population where N_e_ is small relative to 1/U_del_ so that S << T and the population is monomorphic most of the time. **B.** Increasing the mutation rate (by 2X in this example) reduces the average size of HOR arrays because turnover of deletion alleles is faster. If the turnover rate is sufficiently fast, HOR arrays will have insufficient time to grow to large size. **C.** Increasing N_e_ (by 5X in this example) increases the number of segregating alleles (modes in graphs to the right) and increases the largest mode size.

**Supplemental Figure S8.**
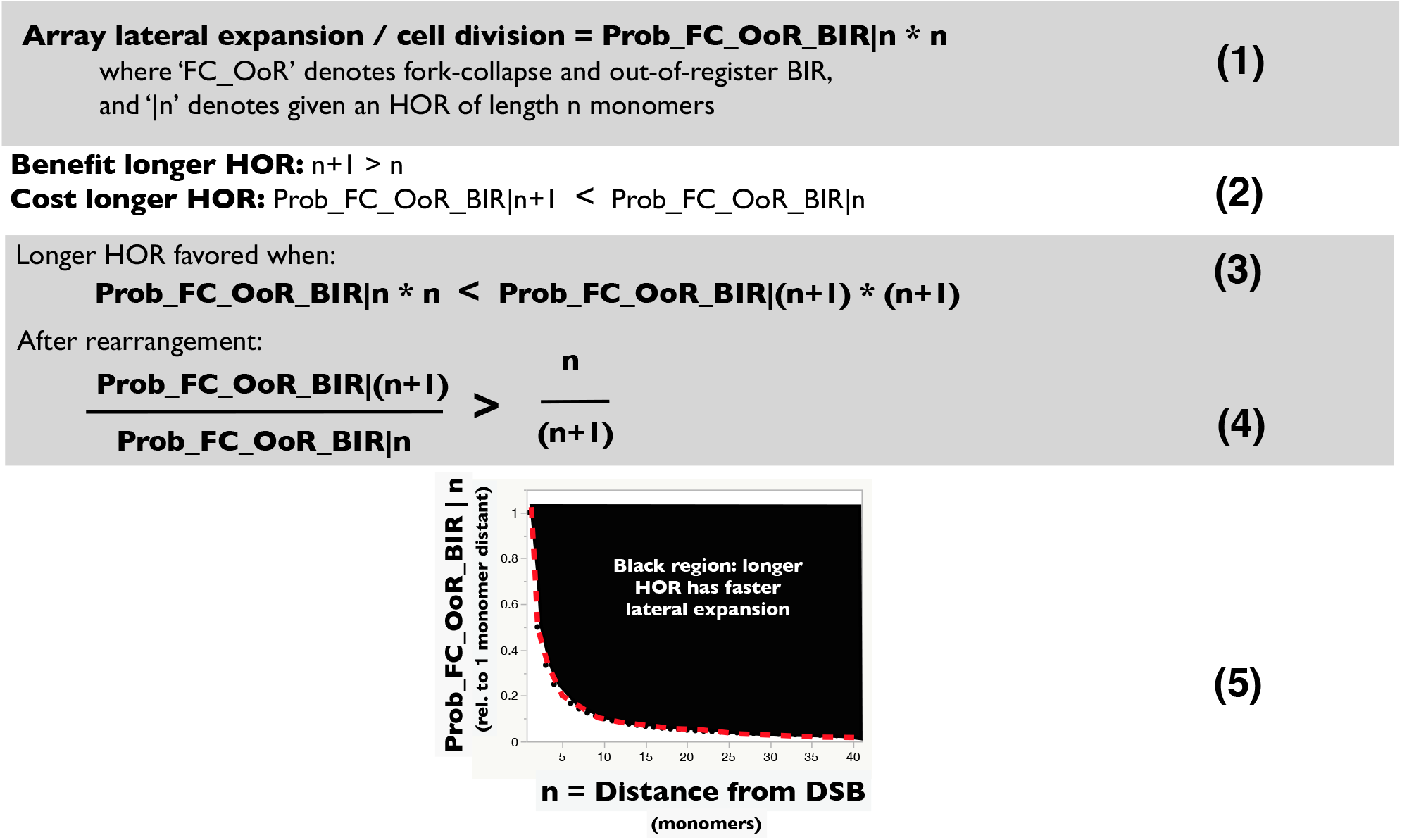
Conditions leading to a longer HOR (length n+1 monomers) having a faster lateral expansion rate compared to a shorter HOR (length n monomers). When fork-collapse within an HOR array is rare per cell division, its rate of lateral expansion per division can be expressed as the product of the probability of a fork-collapse followed by out-of-register BIR (Prob_FC_OoR_BIR) multiplied by the number of monomers in the HOR (n) [equation **(1)**]. This calculation implicitly assumes that BIR will reinitiate DNA replication at the point of closest downstream homology, i.e., n or (n+1) monomers downstream (see last sentence for other re-initiation points). Longer HORs lead to both costs and benefits with respect to lateral expansion rate **(2)**. Expansion **Cost of a longer HOR**: increased distance between the DSB and the closest position of downstream homology (n+1 vs. n monomers), which is expected to reduce the efficacy of out-of-register homology search (Renkawitz et al. 2013) and thereby increase the probability of in-register BIR (causing Prob_FC_OoR_BIR I (n+1) / Prob_FC_OoR_BIR I n) < 1). **Expansion Benefit of a longer HOR**: increased number of monomers added each time a downstream, out-of-register BIR occurs (n+1 vs. n monomers). Longer HORs will have an expansion advantage when inequality **(3)** (or equivalently **(4)** is met, i.e., when the cost of a longer HOR (reduced probability of out-of-register BIR) is less than the benefit (expanding by an additional monomer). These inequalities are met (black region in graph (5) whenever the probability of a downstream, out-of-register BIR declines (with distance from the DSB) more slowly than the red dashed curve (Prob_FC_OoR_BIR I n = 1/n) which is the equality point. Most of the parameter space supports a lateral expansion advantage for longer HORs. ChIP-chip data with antibodies against Rad51 (a protein mediating homology search) indicate a slow decay in the efficacy of homology search with distance from a DSB (much less steep than the red curve in graph (4), indicating a low cost to a longer HOR (Renkawitz et al. 2013) and hence a faster lateral expansion rate for longer HORs. Although this model assumes BIR reinitiates DNA replication at the closest downstream point of homology, the same logic applies to re-initiation of BIR replication at any arbitrary pair of positions of homology on the short (n) and long (n+1) HOR arrays (2n vs. 2(n+1), 3n vs. 3(n+1), and so on).

**Supplemental Figure S9.**
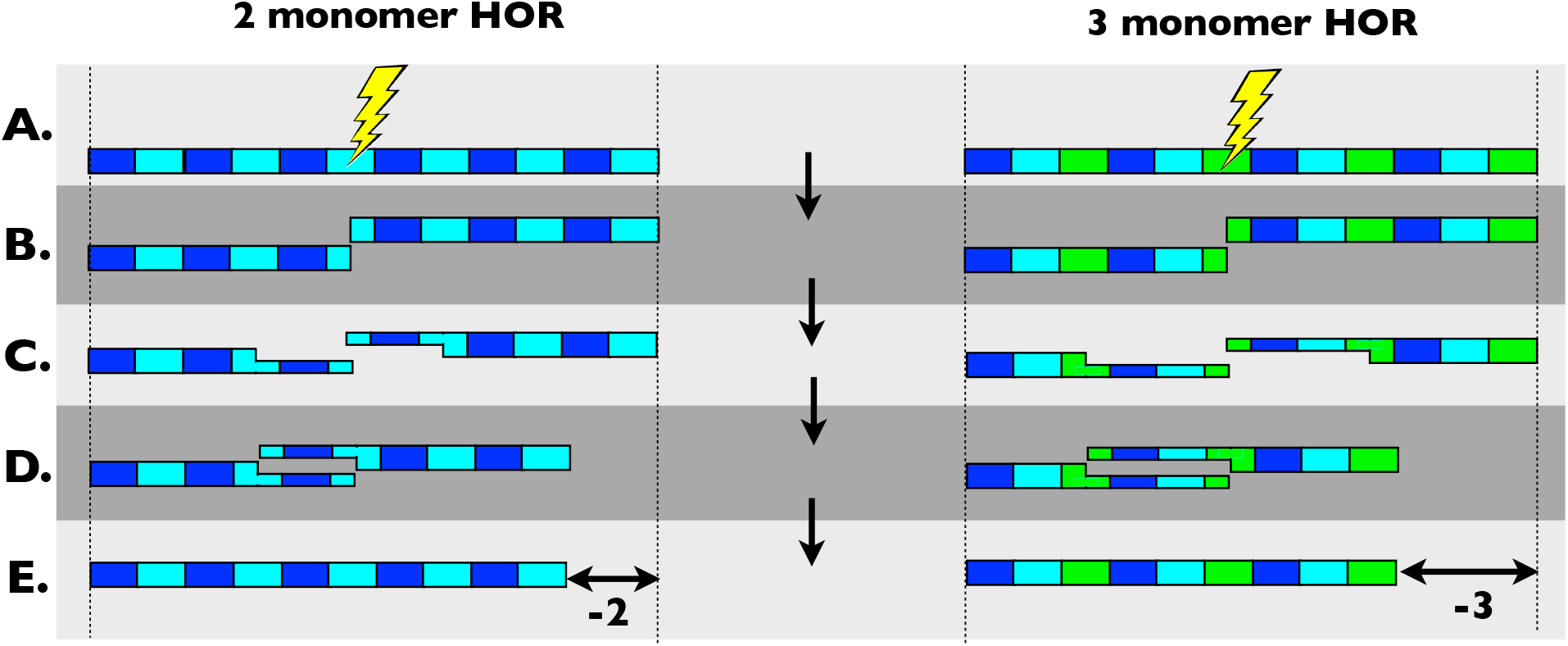
SSA repair of a DSB leads to a larger deletion in a longer HOR. Rectangles represent repeat elements, and those with the same color have the same DNA sequence. **A-B**. DSB (lightning bolt) is generated in the HOR array. **C.** Resection continues until homology of the two ends is found. **D.** Annealing of the two ends and ligation. **E.** The SSA repair creates a longer deletion in the longer HOR array (right; 3 monomers) compared to the shorter HOR array (left; 2 monomers). If the DSB-generating agent causes damage to the strand ends that are removed prior to repair (e.g., via exonucleases), then the removed bases will be replaced via gap-filling after annealing.

**Supplemental Figure S10.**
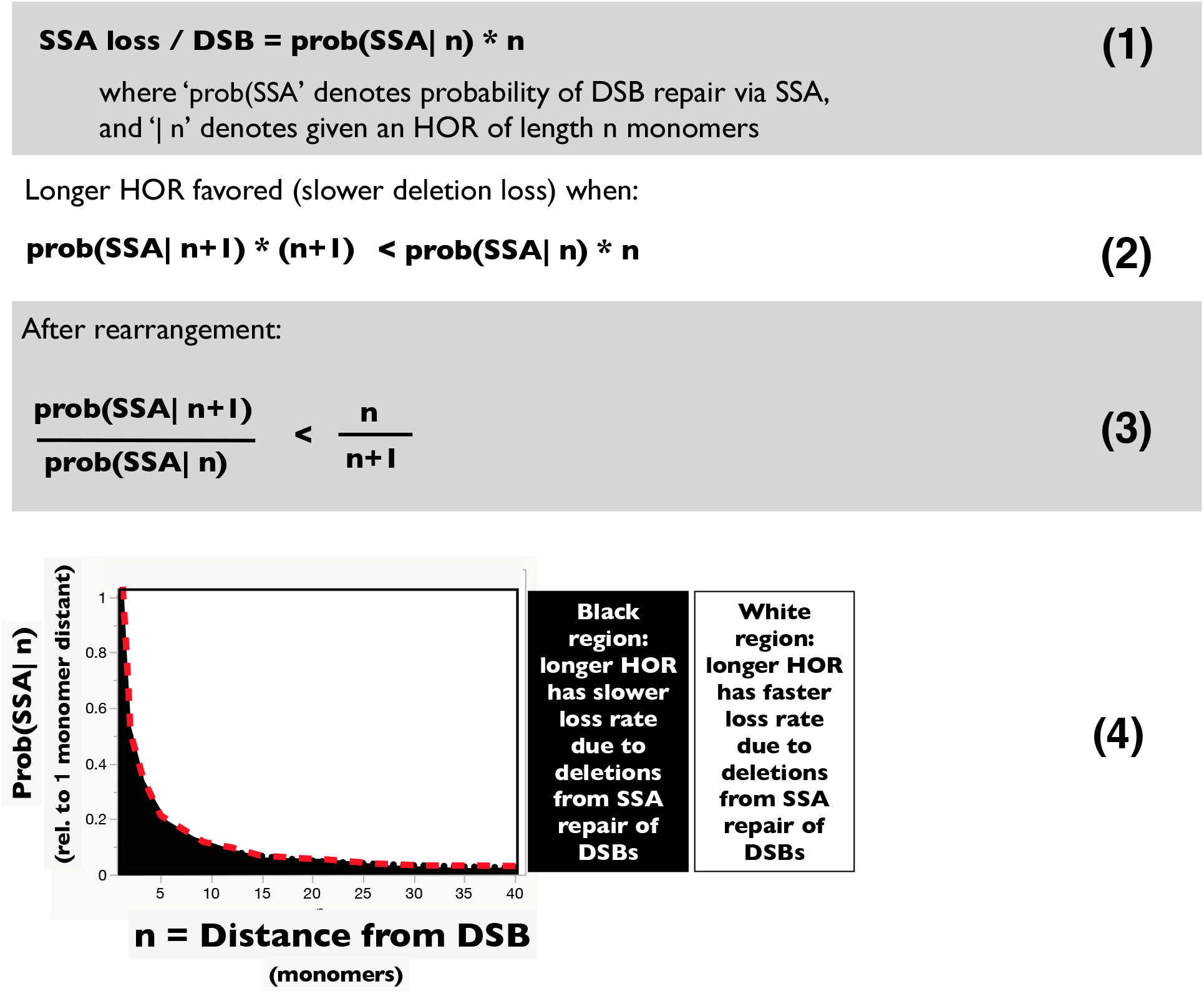
Conditions leading to a longer HOR (length n+1 monomers) having a slower shrinkage rate compared a shorter HOR (length n monomers) during SSA repair of DSBs. Given that a DSB occurs within an HOR array of length n monomers, the loss per SSA-repair is expected to be n monomers (i.e., the length of the HOR = distance between the start points of neighboring repeats) (Sugawara et al. 2000). Let the probability of repair via SSA (compared to deletion-free repair via **H**omology **D**irected **R**epair [HDR], including the SDSA pathway and repair via double Holiday structures). This probability declines with the distance separating repeats (Schildkraut et al. 2005), so the prob (SSA | n+1) < prob (SSA | n). The expected loss by deletion (in monomers) per DSB is equal to prob(SSA | n) * n **(1)**. A longer HOR is favored (fewer deleted monomers per DSB) when the inequality shown in **(2)** [or equivalently **(3)**] are met. This relationship is shown graphically in the figure **(4)**. Longer HORs are favored only when the prob(SSA | n) drops precipitously with distance (dark region below dashed red curve). Data from Schildkraut et al. (2005) show a linear decline in prob(SSA) with distance and a slope that is too shallow to meet the constraints shown in the graph (4), indicating that longer HORs are disfavored via SSA-repair of DSBs.

**Supplemental Figure S11.**
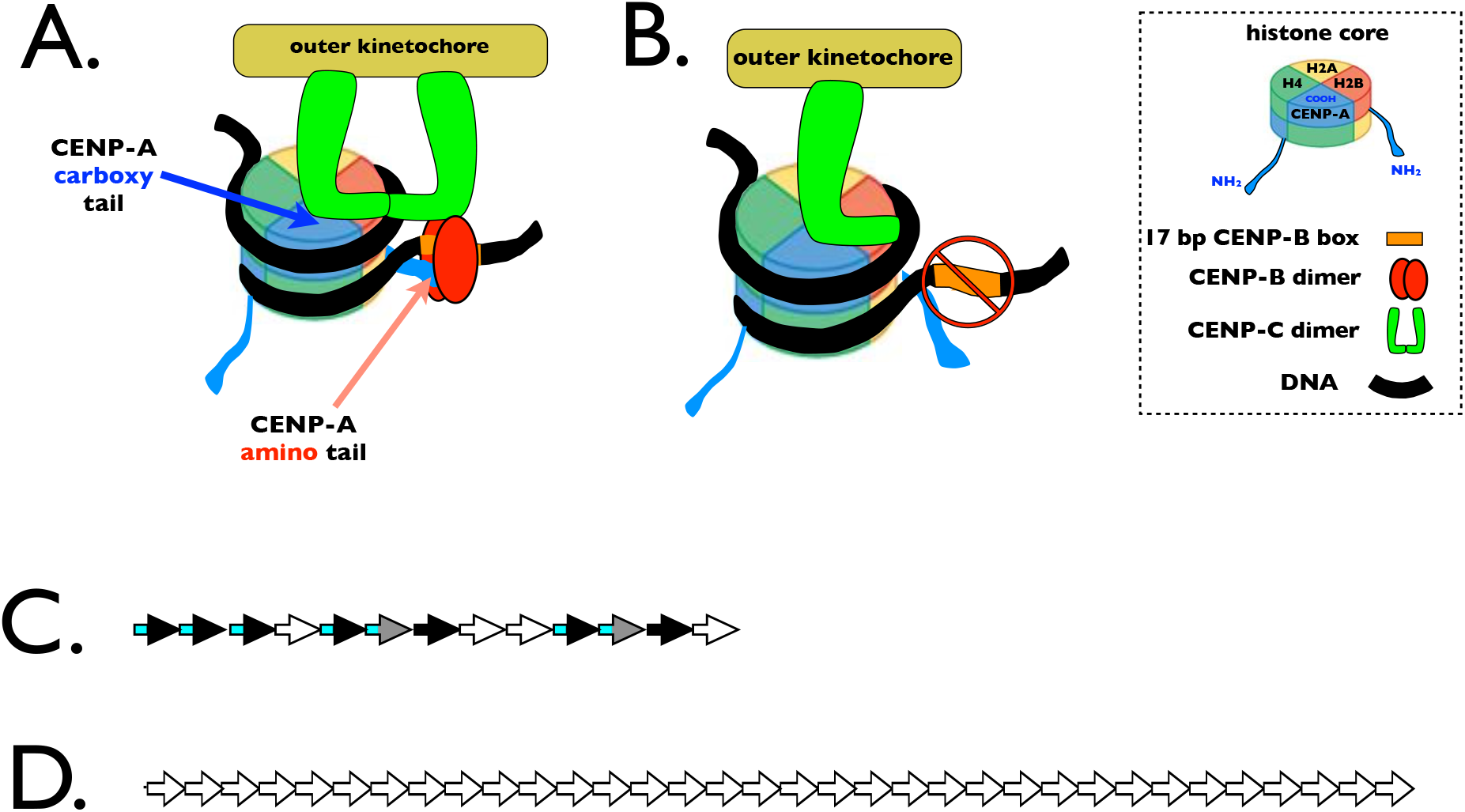
**A.** CENP-C binds to CENP-A-containing nucleosomes at two places: at the the carboxyl end of CENP-A and also near CENP-A’s amino end: with the attachment near the amino-end dependent on the presence of CENP-B. The arrangement shown here follows that proposed by Fachinetti et al. (2015), but the CENP-B might also attach to linker DNA between other nucleosomes that are juxtapositioned in 3D space. **B.** When CENP-B cannot bind the linker DNA because of a mutated b-box (or a complete absence of a b-box on n-box monomers), only one CENP-C can bind each CENP-A nucleosome. (Adapted from Fachinetti et al. 2015). A single CENP-B per nucleosome will feasibly also increase lateral expansion rate because in yeast this protein, when bound to DNA, reduces fork-stalling/collapse (Nakagawa et al. 2002). **C.** A 13 monomer HOR variant found on human chromosome 17 that has a higher density of b-box-containing monomers (7/13) than any other active human HOR: but all of these are arranged in runs of 2-3 b-box-containing monomers. Despite the high density of b-box-containing monomers, the strong deviation from modular b/n-box dimeric structure was associated with 60% less recruitment of CENP-C and 40% less CENP-A compared to the average from all other chromosomes (Aldrup-MacDonald et al. 2016). **D.** The HOR of the human Y chromosome has no b-box monomers, recruits 50% less CENP-C, and typically recruits ~15-20% less CENP-A (Fachinetti et al. 2015; Bodor et al. 2014; Irvine et al. 2004; but see Ross et al. 2016). Small black arrow with blue tail = b-box monomer (blue tail = b-box); Small black arrow with no blue tail = mutated b-box monomer that does not bind CENP-B; Small white arrow = n-box monomer; grey arrow = monomer with a sequence intermediate between the consensus for n-and b-box monomers.

**Supplemental Figure S12.**
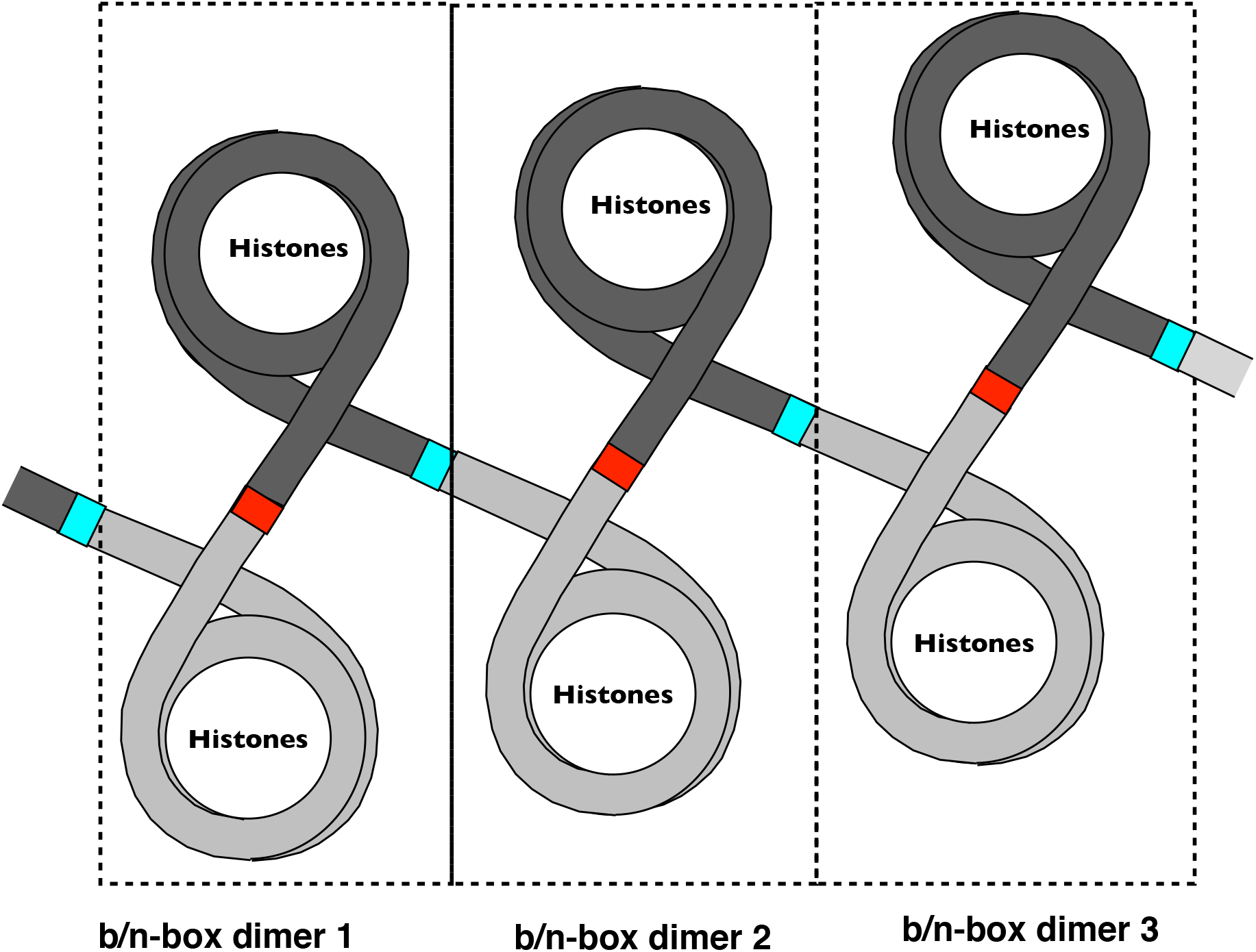
Modular b/n-box dimeric structure positions every nucleosome next to a linker containing: i) a b-box than can bind CENP-B (red region of light grey monomers), and ii) a linker containing an n-box (blue region of dark grey monomers), which feasibly enables every CENP-A nucleosome to recruit two CENP-C per nucleosome (Supplemental Figure S11A,B) and maximizes kinetochore strength. The dimer structure also places non-CENP-B-bound linker DNA adjacent to each CENP-A nucleosome, which –because the large CENP-B molecule (160 kD dimer) would block access to the linker DNA– may facilitate nucleosome positioning, RNA transcription/binding and transient binding of kinetochore proteins during the assembly of kinetochore units.

**Supplemental Figure S13.**
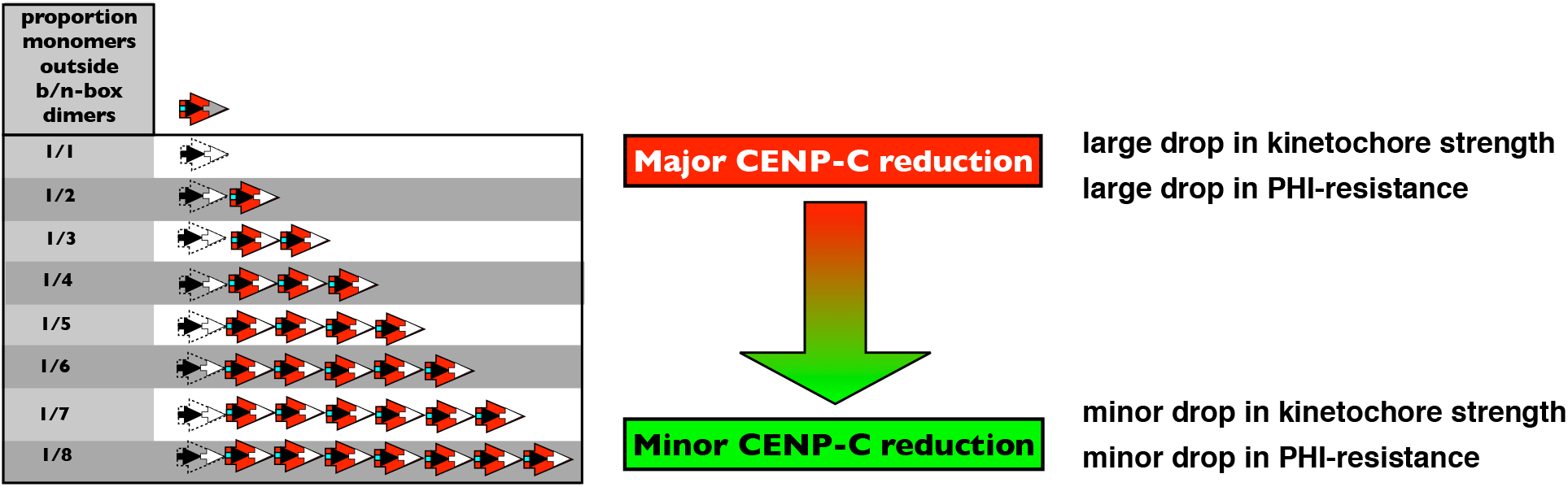
The effect of a mutation that inactivates a single b-box (so it no longer binds CENP-B) within a modular b/n-box HOR of differing lengths. An inactive b-box will feasibly interfere with flanking nucleosomes recruiting CENP-B and cause only one CENP-C (instead of 2) to be recruited to these nucleosomes (see Supplemental Figure S11). When the HOR is short, loss of a single functional b-box per HOR would cause all or a high proportion of nucleosomes to recruit one rather than two CENP-C molecules. The percent loss in CENP-C per nucleosome is expected to be only high in short HORs, so this loss will only be substantially opposed by centromere drive and/or PHI-resistance when the HOR is short. Arrow key as in Figure 10.

**Supplemental Figure S14.**
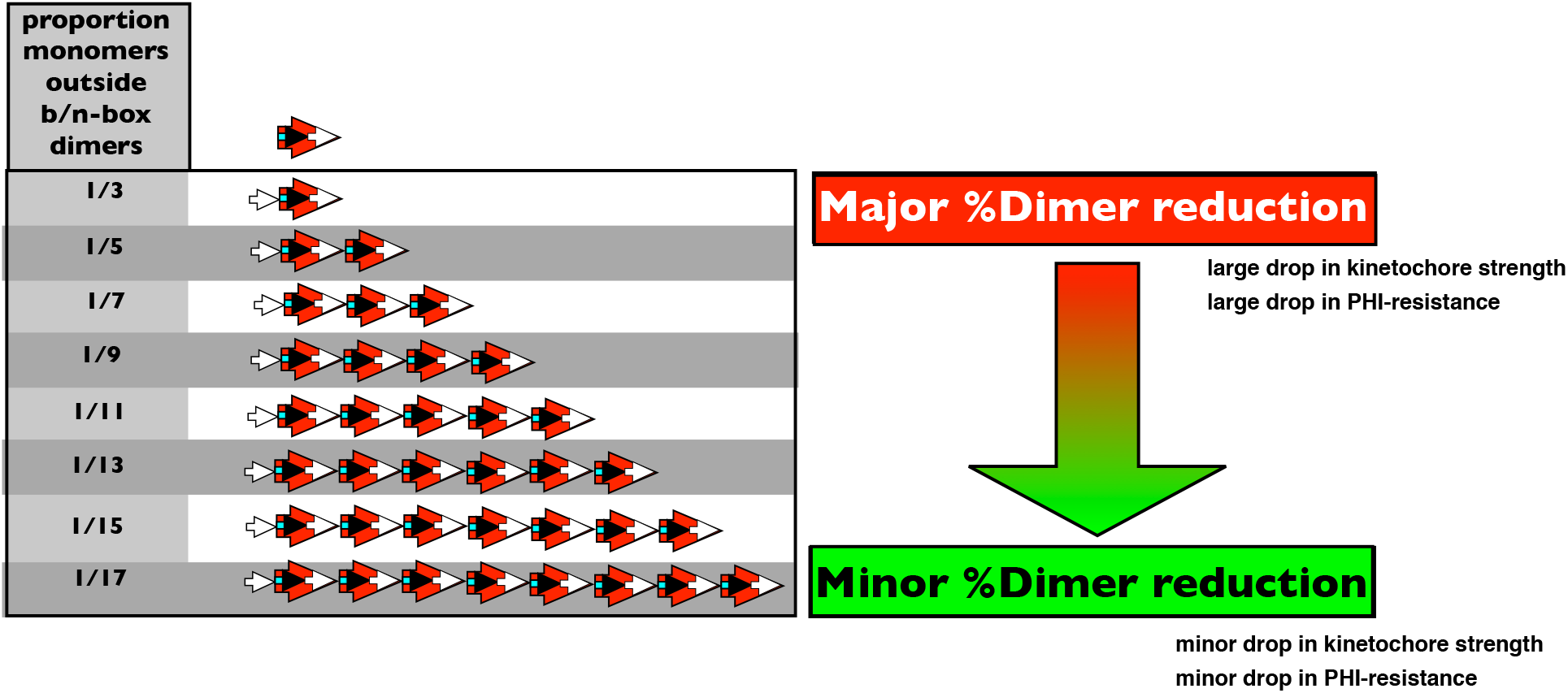
The effect of adding a single n-box monomer to a modular b/n-box HOR of differing lengths. The single n-box monomer dilutes the density of canonical n/b-box dimers to a greater degree in shorter HORs. Assuming b/n-box dimeric structure facilitates robust kinetochore recruitment and robust centrochromatin recruitment and spreading, adding a monomer to a short HOR will have a large effect in reducing both kinetochore strength and PHI-resistance, but this effect will be greatly reduced in long HORs.

